# Self-organization of spermatogenic wave coordinates sustained sperm production in the mouse testis

**DOI:** 10.1101/2024.11.03.621757

**Authors:** Toshiyuki Sato, Yuting I. Li, David J. Jörg, Mitsuru Komeya, Hiroyuki Yamanaka, Hiroko Nakamura, Kodai Hirano, Yohei Kondo, Kazuhiro Aoki, Masahide Takahashi, Hiroki Nakata, Hiroshi Kimura, Takehiko Ogawa, Benjamin D. Simons, Shosei Yoshida

## Abstract

Spermatogenesis takes place in the testis, relying on the ordered turnover of differentiating cells supplied from stem cells. Classic histological analyses have revealed that this process shows hierarchical spatiotemporal patterning known as the spermatogenic cycle, wave, and descent of segmental order, indicative of currently underexplored mechanisms of tissue- and organ-scale homeostasis. Here, using mice, we conducted high-resolution, wide-field, and ‘ultra’ long-term live imaging studies *in vivo* and *ex vivo*, combined with whole-organ mapping of differentiation stages. Such trans-scale measures demonstrate how stereotypic local cell turnover is coordinated into characteristic phase waves propagating along the seminiferous tubules, further organized into organ-scale patterning over the tubule loops. Minimal mathematical modeling shows that such higher-order dynamics can emerge from the local coupling of autonomous oscillators, which are rooted in delayed feedback interplay between stem and differentiating cells via retinoic acid signaling. These findings highlight a self-organization mechanism underpinning organ-scale homeostasis and constant sperm production.

## INTRODUCTION

In long-lived organisms, tissues and organs maintain homeostasis over extended periods, during which multiple generations of differentiating cells are supplied by stem cells and turnover. Through intensive studies, substantial knowledge has been gained into stem cell properties and niche-based regulation. However, the mechanisms that ensure the consistent supply and turnover of differentiating cells remains underexplored.

Mouse spermatogenesis is an ordered process of cell turnover that supports continual sperm production, presenting an ideal system to address this question^1,2^ (Figure 1). In the mouse testis, spermatogenesis takes place within the seminiferous epithelium. The seminiferous epithelium is arranged three-dimensionally into convoluted seminiferous tubules, whose diameter is 150-200μm. Seminiferous tubules, which reach up to 2 meters in length and are split into approximately 10-11 loops, make up the entire testis. Each testis produces around 5 million sperm per day, translating to over 10^9^ sperm per individual over a lifetime.^3^

**Figure 1.**
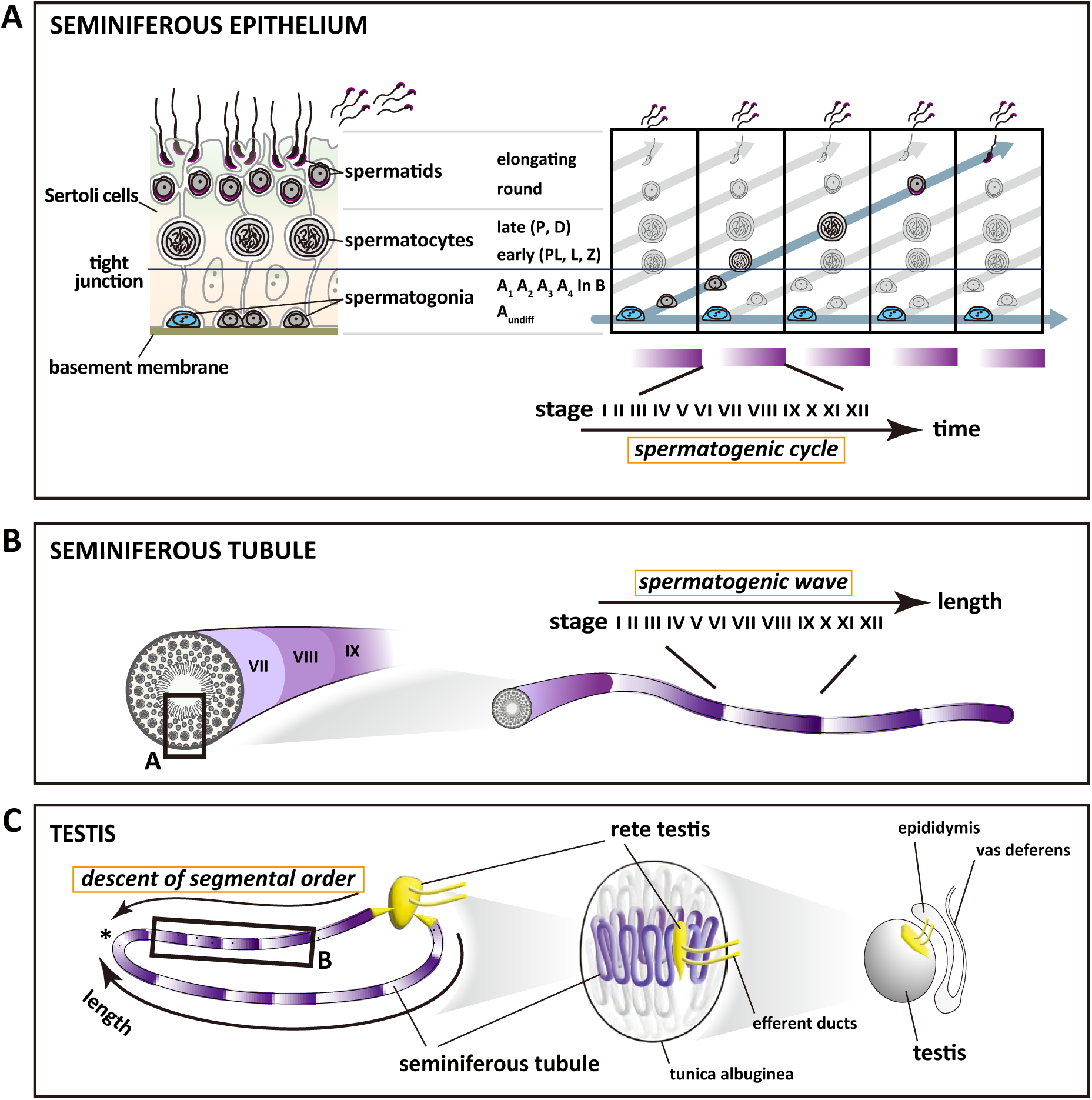
Classic views of the spatiotemporal coordination of germ cell differentiation in mouse testis at distinct scales. **(A)** In the seminiferous epithelium, A_undiff_ (blue) on the basement membrane form the stem cell compartment, supplying differentiating cells that mature into sperm via spermatogonia, spermatocytes, and spermatids. This differentiation, accompanied by cell movement toward the lumen, proceeds periodically to establish the layered architecture (left). As germ cells turn over each generation, the cell association cycles, a phenomenon known as the spermatogenic cycle (right, rectangles, and purple gradient). Based on cell associations, this cycle is morphologically divided into stages I to XII (see Figure S1 for details). PL, preleptotene; L, leptotene; Z, zygotene; P, pachytene; D, diplotene. **(B)** On larger scales, the seminiferous epithelium is shaped into seminiferous tubules. The rectangle indicates the scale shown in (A). In the seminiferous tubules, each spermatogenic cycle stage extends over a cylindrical segment (left). Along the tubule length, the stages typically follow their chronological order (I to XII and back to I) as illustrated by purple gradients (right). This organization is known as the spermatogenic wave. **(C)** At the organ scale, seminiferous tubules form long loops connected at both ends to the rete testis, the common outlet for sperm (left). The rectangle indicates the scale described in (B). About 10 loops compose a single testis (middle, right). Typically, the spermatogenic wave is arranged so that the stage number descends away from the rete testis, known as the descent of segmental order (shown by the purple gradient on the left side of the panel). The asterisk marks the site of reversal of the wave direction.

In the seminiferous epithelium, the entire process of stem cell self-renewal and commitment, differentiation, maturation and sperm production occurs within a sheet of somatic Sertoli cells, which form tight junctions (also called blood-testis barrier) separating the basal and adluminal compartments (Figure 1A). A minority population of undifferentiated spermatogonia (A_undiff_) residing on the basement membrane functions as stem cells, balancing self-renewal and the production of committed A_1_ spermatogonia. A_1_ spermatogonia experience a series of mitoses on the basement membrane, progressing through A_2_, A_3_, A_4_, intermediate (In), and B spermatogonia before initiating meiosis. During meiosis, early spermatocytes detach from the basement membrane and translocate to the adluminal compartment through exquisite reorganization of Sertoli cells’ tight junction.^4,5^ After meiosis, haploid cells gradually move toward the lumen, transforming from round spermatids to elongating spermatids before being released through spermiation. Spermatogonia, spermatocytes, and spermatids form syncytia, connected by intercellular bridges that result from incomplete cytokinesis during mitotic and meiotic divisions. A_undiff_ includes singly isolated (A_s_) cells and syncytia containing as many as 16 cells, while tens to hundreds of spermatocytes may form a syncytium.^6,7^

Notably, the A_undiff_-to-A_1_ transition and subsequent differentiation proceed in a locally synchronized, temporally periodic manner (Figure 1A, right). This leads to a layered organization of the seminiferous epithelium, with maturing cell generations positioned more centrally. Vertically aligned germ cells display definite associations, indicative of stereotypic coordination between cell generations^8,9,10^ (Figure S1). As each generation of sperm is released into the lumen, younger cell generations transition into a more mature state, restoring the tissue periodically to the same composition. This periodic change in the germ cell composition is known as the “spermatogenic cycle” (or cycle of seminiferous epithelium) with each cell association defining a certain “stage” of the cycle. In mice, S-phase labeling studies have estimated the cycle period to be 8.6 to 8.9 days, which is divided into 12 distinct stages (I to XII), with stage-specific durations varying from around 8 to 22 hours.^11,12^ During adult homeostasis, Sertoli cells are post-mitotic and long-lived, while their gene expression and morphology change in synchrony with the spermatogenic cycle.^13,14,15,16,17,18^

In mouse, each stage of the cycle extends uniformly over cylindrical segments of the seminiferous tubule (Figure 1B). Along the tubule length, such segments are typically found in an ordered sequence from stage I to XII and then back again to stage I (Figure 1B, right). Given the ordered temporal transition of the stages, this spatial arrangement indicates that the stage sequence moves along the tubule, a phenomenon termed the “spermatogenic wave” or “wave of the seminiferous epithelium”.^8,19,20,21,22^

At larger length scales, the seminiferous tubule loops, each of up to 20 cm in length, are minimally branched, with ends open to the outlet of sperm, called rete testis^1,23^ (Figure 1C). Histological observations show that the stages of the cycle are typically arranged in decreasing order (e.g., XII, XI, X, and so on) in the direction away from the rete, with the order reversing around the middle of the loop. This large-scale patterning is known as the “descent of segmental order”.^24,25,20,26^

The spermatogenic cycle, wave, and descent of segmental order have shown that seminiferous tubules exhibit well-coordinated cell turnover, captivating scientific interest. However, the trans-scale nature of this process—from the micrometer scale of individual cells to the tens of centimeters scale of tubule loops—has hindered attempts to achieve a mechanistic understanding of the underlying cell behaviors. Critically, the dynamics of this system have been inferred only indirectly from static measurements obtained from the analysis of fixed specimens, limiting insights into the core dynamical principles and regulatory mechanisms. In addition, static observations have revealed that the spatial wavelength (the tubule length spanning the entire sequence of 12 stages) is highly variable. Moreover, although the stages generally follow a sequential order, there are short stretches—known as ‘modulations’—where the stage order is reversed.^25,20,26^ These irregularities make capturing the spatiotemporal dynamics of mouse spermatogenesis more challenging.

To address the mechanism of coordinated cell turnover, it is crucial to determine how the A_undiff_-to-A_1_ transition is regulated. Among various factors regulating the fate of spermatogenic stem cells, retinoic acid (RA) is known to be a critical driver of this process.^27,28^ Local RA concentration within seminiferous tubules varies in synchrony with the spermatogenic cycle, showing pulse elevations at stages VII to VIII.^29,30^ This RA pulse induces the A_undiff_-to-A_1_ transition, as well as other differentiation events such as initiation of meiosis and spermiation.^31,30^ Further, when germ cell differentiation is halted by RA depletion, its restoration synchronizes the stage of spermatogenic cycle across the testis, followed by gradual stage dispersion to recover the original distribution.^32,33,34^ Another cue arises from a rat-to-mouse germ cell transplantation study: When rat germ cells are nourished by mouse somatic cells, the reconstructed spermatogenesis exhibits a 13-day cycle, consistent with the natural cycle of the rat.^35^ These experiments indicate that changes in RA concentration are crucial to control the germ cell differentiation timing, and that germ cells play a crucial role in establishing the tempo of the spermatogenic cycle. However, their specific functions remain to be elucidated either at the cell biological level or from a dynamical systems perspective.

To overcome current limitations and capture the spatiotemporal coordination of germ cell differentiation, continuous live imaging across broad temporal and spatial scales is vital. Previously, an intravital (*in vivo*) live-imaging platform was developed that enabled the observation of fluorescence-labeled cells within the testis of adult mice kept under anesthesia for up to 3 days.^36^ Yet, this duration represents only a fraction of the spermatogenic cycle. By contrast, advances in the *ex vivo* culturing of seminiferous tubules using a gas-permeable microfluidic device allows spermatogenesis to be maintained for several months,^37^ providing the opportunity to develop continuous live imaging over periods greatly in excess of the cycle time. At the organ scale, the anatomy of all constituent seminiferous tubule loops and the stage distributions have been reconstructed in 3D from serial sections of whole testes.^23,26^

Here, we refined live-imaging platforms to continuously film germ cell turnover within mouse seminiferous tubules for up to 9 days *in vivo* and one month *ex vivo*. Combining these observations with precise spatial mapping of the stage distribution across tubule loops, we captured quantitatively the spermatogenic wave dynamics across multiple length and time scales. Based on these measures, we developed a minimal mathematical framework to understand the nature of the local cell turnover and its large-scale coordination along the seminiferous tubules. Together, these results suggest that the robust phase wave dynamics emerges as a self-organization phenomenon within the context of spatially homogeneous seminiferous tubules, independent of external positional cues or scale-specific mechanisms.

## RESULTS

### Trans-scale live-imaging of spermatogenesis *in vivo* and *ex vivo*

To visualize germ cell differentiation and the dynamics of the spermatogenic cycle and wave, we used *Stra8-EGFP* mice harboring a modified BAC expressing EGFP-tagged Stra8 (Stimulated by retinoic acid gene 8) protein.^38^ Endogenous Stra8 is expressed in early spermatocytes during stages VII to IX and in spermatogonia during stages VII to I in response to the RA pulse that occurs in stages VII-VIII. These Stra8 expressions drive meiosis initiation and the A_undiff_-to-A_1_ transition, respectively^29,31,39^ (Figure 2A; Figure S2A). *Stra8-EGFP* mice showed consistent GFP fluorescence (Figure 2B; Figures S2B-F). In whole-mount preparations, seminiferous tubule segments were categorized as belonging to stages VII to IX based on GFP expression of both spermatocytes and spermatogonia (magenta), X to I based on GFP expression of spermatogonia only (green), and II to VI based on the absence of GFP expression (cyan) (Figure 2C; Figures S2G-J).

**Figure 2.**
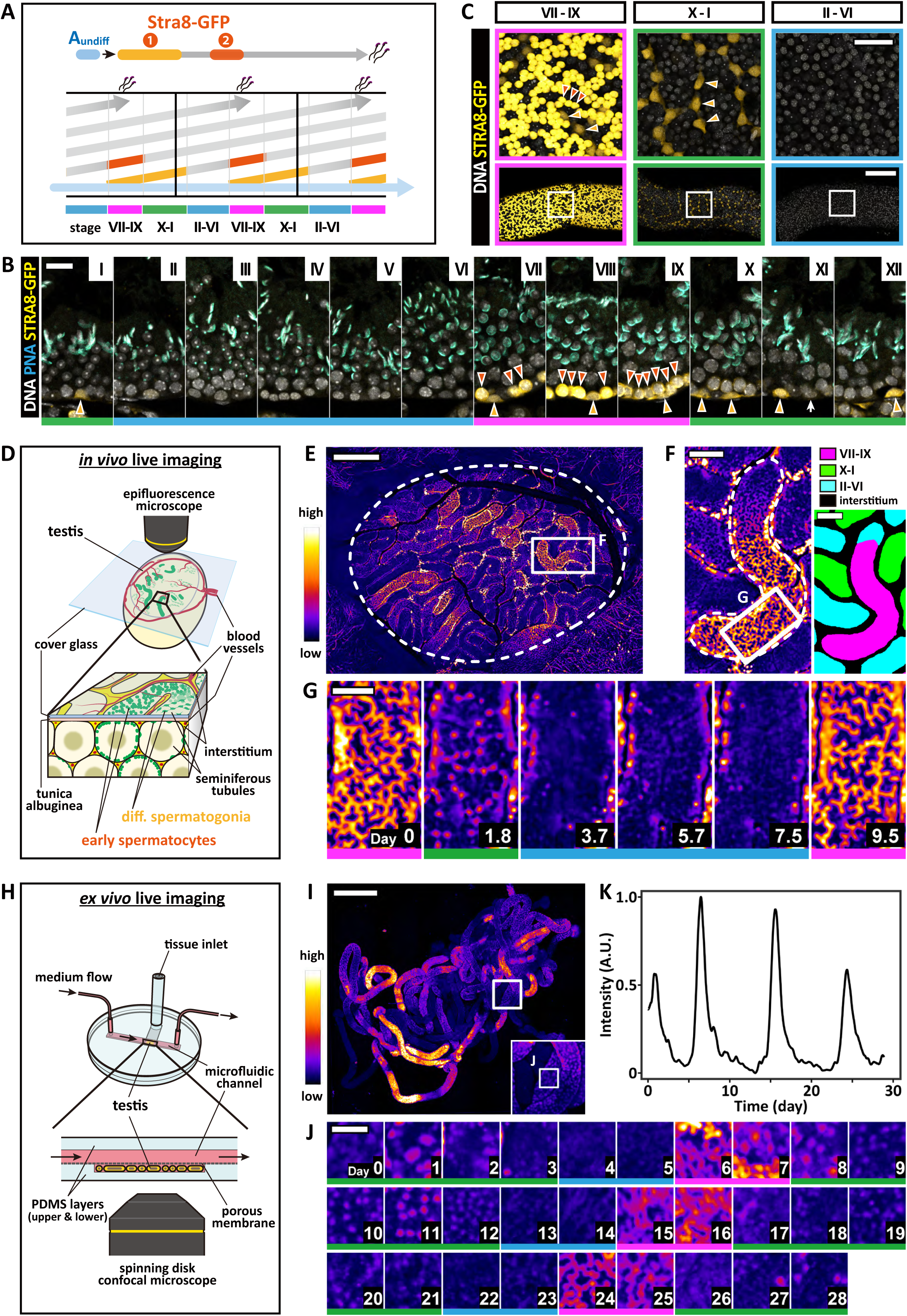
*In vivo* and *ex vivo* live-imaging studies using *Stra8-EGFP* mice visualize the spatiotemporal coordination of germ cell differentiation. **(A-C)** Characterization of *Stra8-EGFP* mouse testis. (A) Stra8 is expressed in early differentiating spermatogonia (1, light orange) and early spermatocytes (2, dark orange), dividing the spermatogenic cycle into three phases: Stra8^+^ spermatocytes and spermatogonia (stages VII-IX; magenta), Stra8^+^ spermatogonia only (stages X-I; green), and no Stra8^+^ cells (stages II-VI; cyan); see Figure S2A for details. (B) Cross sections of Stra8-EGFP seminiferous epithelia show Stra8-GFP^+^ spermatogonia and spermatocytes (light and dark orange arrowheads), defining stages VII-IX (magenta), X-I (green), and II-VI (cyan). White arrow indicates undifferentiated spermatogonia; see Figure S2B for details. (C) Whole-mount seminiferous tubules stained for Stra8-GFP, representing the three phases described in (A-B). Upper panels magnify the rectangular regions in the lower panels. Scale bars: 20 µm (B), 50 µm (C upper), and 200 µm (C lower). **(D-G)** *in vivo* imaging of the *Stra8-EGFP* mouse testis. (D) Schematic of the observation strategy. (E) Compiled image of the entire testis surface at t=0, showing GFP intensity by pseudocolor. (F) Magnified image of the rectangle region in (E), segmented into three phases based on the Stra8-GFP pattern (right). (G) Selected frames of time-lapse images in the rectangle region in (F) (Video S2). Scale bars: 1000 µm (E), 200 µm (F), and 100 µm (G). **(H-K)** *ex vivo* live-imaging of *Stra8-EGFP* mouse testis explants. (H) Culture and observation scheme. (I) Representative compiled image covering the entire explant of a single Stra8-EGFP testis at t=0, with a magnified image of the rectangular region in the lower right inset. (J) Selected frames of time-lapse images in the square area of the inset in (I) at indicated elapsed days (Video S3). (K) Time course of the normalized average intensity in (J). Scale bars: 500 µm (I) and 50 µm (J).

To measure the dynamics over an entire spermatogenic cycle, we needed to extend the imaging period from our previous *in vivo* live-imaging settings^36^ (Figure 2D and Methods). By refining the procedures, including the anesthesia and nutrient supply, we achieved to live-image the testes of *Stra8-EGFP* adult mice in time lapse, at a rate of one frame per 20 minutes for up to 9 days (Figures 2E-G; Figure S3A; Video S1). Time-lapse images taken at multiple XY positions were assembled into movies that cover one side of the testis surface of about 6 x 5 mm, displaying dozens of tortuous tubule segments (70-80mm in summed length, comprising about 4% of a whole testis) (Figure 2E). The observed transition between Stra8-GFP-based stages (magenta, green, and blue) verified that the spermatogenic cycle progressed consistently in this setting (Figure 2G; Video S2).

To observe longer tubule regions over multiple cycles, we further developed an *ex vivo* live-imaging platform based on a previously established culture method capable of supporting spermatogenesis over months using a gas-permeable, polydimethylsiloxane (PDMS) microfluidic device^37^ (Figure 2H and Methods). For reasons currently unknown, this method requires the use of juvenile tissue as source material. We therefore started with 10 to 11-day-old mouse testes in which germ cells have developed as far as spermatocytes.^40,41^ In culture devices modified for live imaging, the explant was supplied continually with fresh medium through a porous membrane from the top, while imaged intermittently from the bottom at one frame per 20 minutes for around one month. The early age of the starting material, along with the potential for culture artifacts, requires careful extrapolation of the *ex vivo* observations to address adult homeostasis *in vivo*.

Using this *ex vivo* setting, we first questioned whether the spermatogenic cycle occurs, persists, and can be imaged continuously. Strikingly, we observed periodic changes of Stra8-GFP expression and its propagation along the explant tubules, mirroring the *in vivo* situation in the absence of the organ context, e.g., blood flow, innervation, or hormonal regulation (Figures 2I-K; Figure S3B; Video S3).

### Locally coordinated germ cell turnover underpins the spermatogenic cycle

Based on the acquired live-imaging data, we questioned how germ cell differentiation is coordinated over time, across different length scales. We first zoomed into submillimeter-scale small regions of the seminiferous epithelium using our *in vivo* platform (Figures 3A-B; Video S4). We found that in regions where Stra8-GFP^+^ cells were initially absent, intense fluorescence developed sharply in early spermatocytes, indicating the initiation of meiosis triggered by the RA pulse in stage VII. The Stra8-GFP signal showed strikingly synchronous changes over tens to hundreds of spermatocytes, which likely belong to one or a few syncytia. The Stra8-GFP signal in these cells lasted for up to 60 hours, consistent with the duration of stages VII to IX.^11,12^ Fading of the signal in chains of spermatocytes, accompanied by their deformation, was consistent with the initiation of their vertical translocation from the basal to adluminal compartments mediated by reorganization of the tight junction between Sertoli cells, a process starting in stage IX^5^ (Figure 3C; Figure S4A; Video S5).

**Figure 3.**
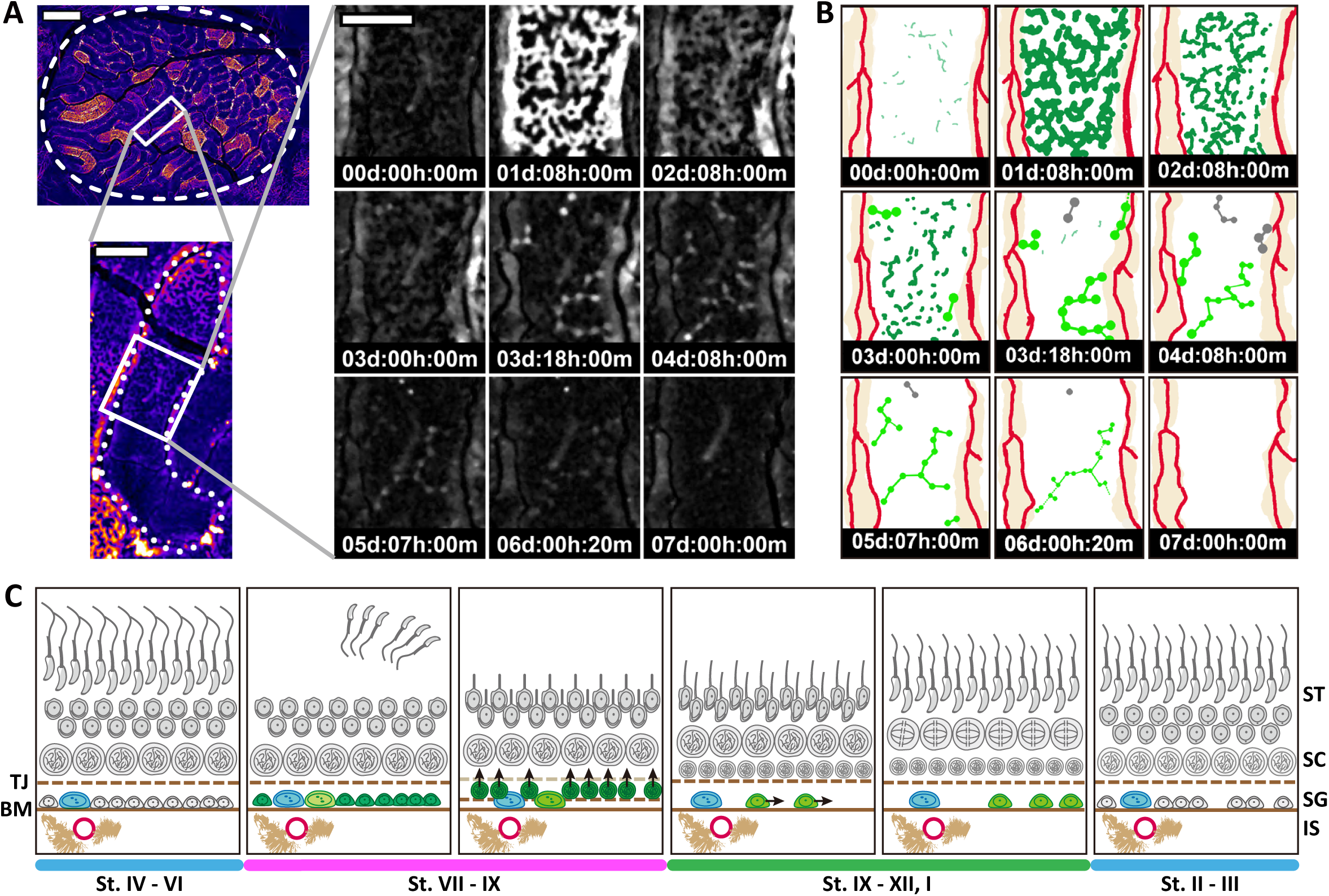
Behaviors of Stra8-GFP^+^ spermatocytes and spermatogonia are locally synchronized and coordinated across cell generations. **(A)** Selected *in vivo* time-lapse images zoomed into a small region of the seminiferous epithelium at indicated time points as day:hour:minute (Video S4). Scale bars: 1000 µm (left upper), 200 µm (left lower), and 100 µm (right). **(B)** Trace of the images in (A): dark green indicates Stra8-GFP^+^ spermatocytes; light green and gray connected circles indicate Stra8-GFP^+^ spermatogonia surviving and dying during the filming period, respectively. Blood vessels and interstitium are traced in red and beige, respectively. **(C)** Scheme showing the turnover of germ cell generations during the spermatogenic cycle in the seminiferous epithelium, reconstructed from static information. Stra8^+^ spermatocytes (dark green) and spermatogonia (light green), as well as A_undiff_ (blue, not visible in live imaging), are highlighted. Consistent with (B), red circles and beige objectives represent blood vessels and interstitial cells (e.g., Leydig cells and macrophages), respectively. Arrows schematically indicate the directions of cell movements. SG, spermatogonia; SC, spermatocytes; ST, spermatids; TJ, tight junction; BM, basement membrane; IS, interstitium.

As the Stra8-GFP^+^ spermatocytes undergo vertical translocation, the next generation of Stra8-GFP^+^ spermatogonia spread horizontally throughout the basal compartment from regions near the interstitium and vasculature (Figures 3A-C; Figure S4A). We reasoned that this migration coincides with the A_undiff_-to-A_1_ transition, based on its consistency with a previous *in vivo* live-imaging study using Ngn3/EGFP mice.^36^ Stra8-GFP^+^ A_1_ spermatogonia were observed to undergo one or two mitotic divisions, giving rise to A_2_ and A_3_ spermatogonia while extending their syncytial size, before losing the GFP signal altogether. Some spermatogonial syncytia underwent cell death (Figures 3A-B), corroborating the abundance of cell death suggested to occur in the A_2_ and A_3_ spermatogonia.^42^ Finally, although not directly visible in *Stra8-EGFP* mice, A_3_ spermatogonia are known to undergo mitotic divisions to become A_4_, In, and B spermatogonia and give rise to Stra8-GFP^+^ spermatocytes in the next cycle (Figures 2A and 3C). These observations demonstrate exquisite spatiotemporal coordination of germ cell turnover, which underpins the stereotypic progression of the spermatogenic cycle.

### Seminiferous tubules comprise an array of stage-synchronized cell clusters separated by small time shifts

We then asked how locally synchronized germ cell differentiation is coordinated over larger length scales along the tubules. From *in vivo* live-imaging data, we found that the Stra8-GFP expression was highly synchronized among clustered spermatocytes extended over a tubule length of around 100-400 μm (likely comprising one or a few syncytia), showing a small time shift of about 1 to 6 hours from neighboring clusters (Figure 4A; Video S6). For quantitative verification, we mapped the time-lapse images onto a square grid and quantified the temporal change in GFP intensity in each square, largely reflecting the signal from spermatocytes (Figures 4B-C). Subsequent hierarchical clustering analysis detected discrete regions in which the signal intensity followed highly synchronized changes, with distinct but variable time shifts between neighboring regions along the tubule.

**Figure 4.**
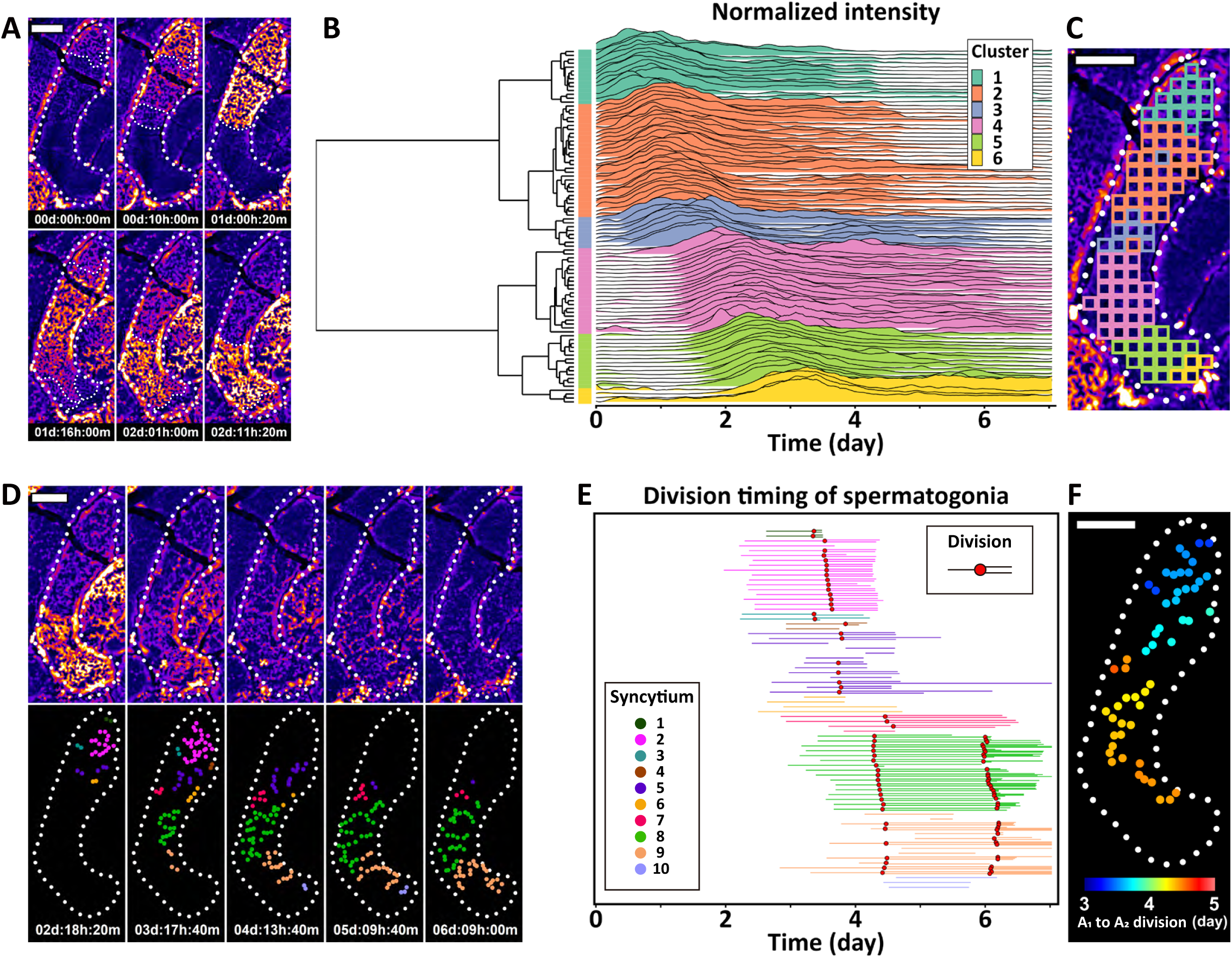
Locally synchronized germ cell clusters are aligned along the seminiferous tubules with small time shifts. **(A-C)** Local coordination of Stra8-GFP expression timing in spermatocytes. (A) Time-lapse *in vivo* images of a representative seminiferous tubule segment (outlined by rough dotted lines; the same segment in Figure 3A). See Video S6 for the full series. Note the highly synchronous Stra8-GFP expression within local clusters of spermatocytes, bounded by fine dotted lines. (B, C) Quantitative verification of the observation in (A). The tubule segment was divided into 60 small grids (C), and the change in GFP intensity averaged in each grid, reflecting Stra8-GFP expression in spermatocytes, was profiled over the imaging period in the ridge plot (B). Synchrony and heterochrony between grids were evaluated using hierarchical clustering, revealing 6 major clusters (dendrogram in B) shown by different colors in the ridge plot (B). The distribution of these clusters is indicated by the same colors in (C). Note the high synchrony within grids of the same cluster, their spatial proximity, and the variable time shift (up to 6 hours) between clusters, reinforcing the visual observations in (A). Scale bars: 200 µm in both (A) and (C). **(D-F)** Spatial coordination of Stra8-GFP^+^ spermatogonia in motion and division. (D) Selected time-lapse images following those in (A) (upper) and traces of spermatogonia, color-coded for each presumptive syncytium (lower; Video S6). (E) Division profiles of Stra8-GFP^+^ spermatogonia per syncytium, with horizontal lines showing the observation period of individual cells, color-coded as in (D); red dots indicate the timing of cell divisions. (F) Position and timing (color-coded) of A_1_-to-A_2_ divisions. Scale bar: 200 µm in both (D) and (F).

Following the progression of Stra8-GFP expression domains in spermatocytes, we consistently observed coordinated migration and division of Stra8-GFP^+^ spermatogonia (Figure 4D; Video S6). In particular, spermatogonia belonging to the same syncytium migrate collectively from the proximity of vasculature/interstitium to spread over the basement membrane and undergo synchronous cell divisions. The timing of these events varies systematically along the tubule length (Figures 4E-F). Notably, a syncytium of GFP^+^ spermatogonia often extended across the borders between synchronized clusters of the spermatocytes (Figure S4B), suggesting that the spatial extent of synchronized cell clusters is not perfectly aligned vertically across cell generations.

Altogether, these results show that local clusters of cells whose differentiation timing is highly synchronized, at least in part due to forming syncytia, are aligned along the tubule length showing systematic time shifts. These clusters are significantly smaller in size than the typical area that represents one of the 12 stages of the cycle. In accordance with this, the observed typical time shifts between adjacent clusters (1-6h) accounts for only a few percent of an entire cycle (9d), much shorter than that required to complete a single histologically recognized stage (8-22h). These observations imply a delicate and systematic interaction between neighboring synchronized cell clusters, a behavior hidden from previous observations based on the study of fixed specimens.

### Spermatogenesis progresses in phase waves with characteristic irregularities

Then, we shifted our focus to the dynamics of cell turnover over larger length scales. From *in vivo* imaging of tubule segments up to 2mm long, we observed that regions with Stra8-GFP expression in spermatocytes propagate along the tubule, followed by those with the migration and division of Stra8-GFP^+^ spermatogonia (Figure 5A; Video S7). These dynamics were captured by mapping the GFP signal intensity, predominantly reflecting the high GFP expression in spermatocytes, as a kymograph. The results exhibited a stepwise progression, corresponding to the arrayed configuration of synchronized clusters with time shifts as described earlier, and varying wave velocities between separate tubule segments (Figure 5B; Figures S5A-C). In *ex vivo* imaging, we observed the propagation of traveling waves over sequential cycles, with kymographs typically showing parallel oblique lines with steps (Figures 5C-E; Video S8). These observations provide a vivid demonstration of how local cell turnover (spermatogenic cycle) is organized into traveling waves (spermatogenic wave).

**Figure 5.**
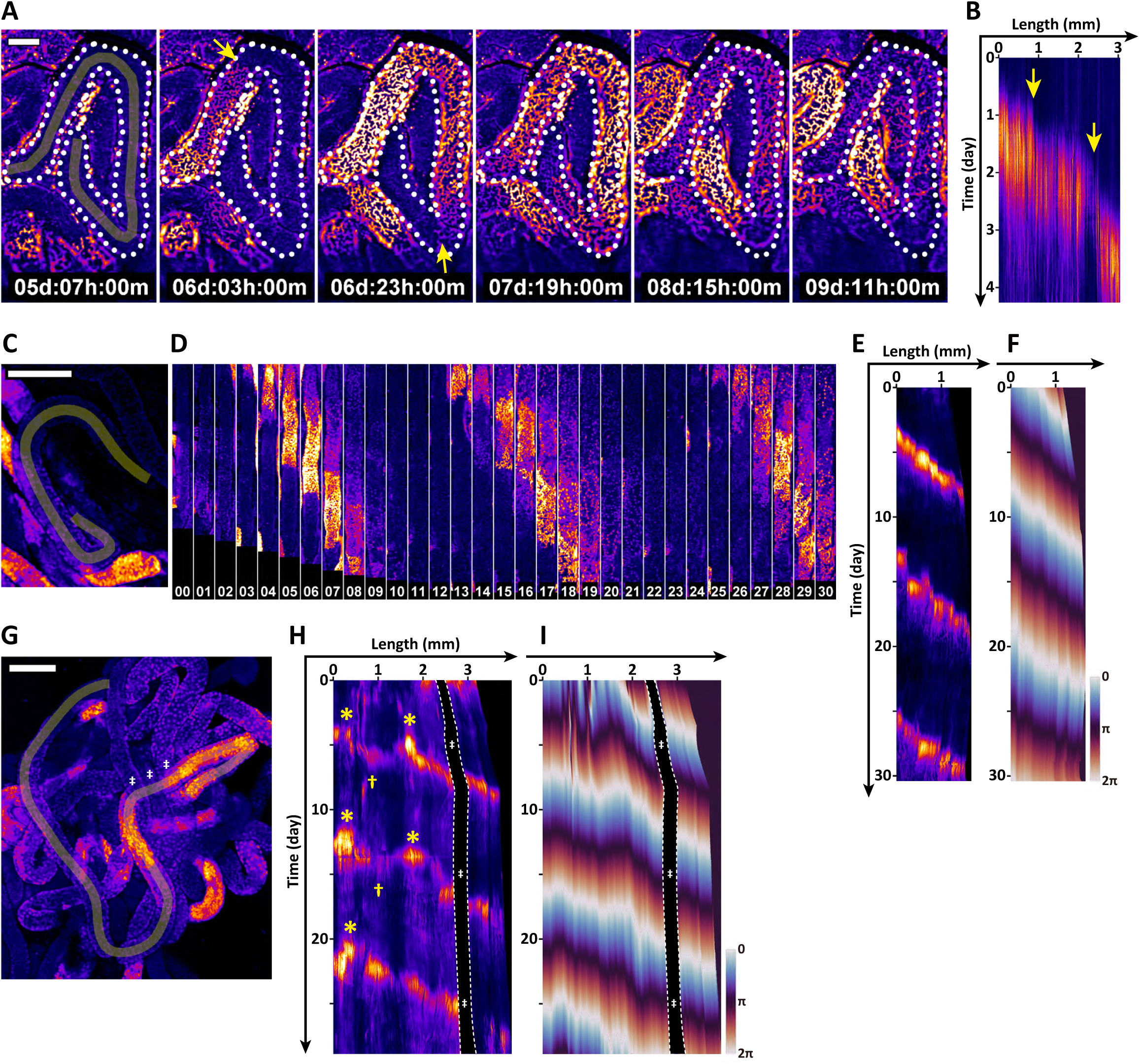
Germ cell differentiation is organized into phase waves propagating along the seminiferous tubules with particular irregularities. **(A-B)** Wave propagation observed in *in vivo* live imaging (Video S7). (A) Selected time-lapse images of a lengthy tubule segment outlined by dotted lines, showing the propagation of Stra8-GFP signal along the tubule length. (B) Kymograph showing the time evolution of GFP signal intensity along a yellowish line on the tubule segment in (A). Yellow arrows in (A) and (B) indicate the points showing significant changes in wave speed. Scale bar: 200 µm in (A). **(C-F)** Repeated traveling phase waves observed by *ex vivo* live imaging (Video S8). The seminiferous tubule region marked by a yellowish line in (C) was computationally straightened and tiled in (D), with elapsed time indicated in days. The kymograph of the GFP signal intensity (E) and its transformation into the phase of oscillation (F). Scale bar: 200 µm in (C). **(G-I)** Irregular and changing phase-wave dynamics observed *ex vivo* (Video S9). A seminiferous tubule marked by a yellowish line in (G) was computationally straightened and processed for kymographs in GFP intensity (H) and after phase transformation (I). Note the irregular pattern, with yellow asterisks and daggers indicating points of wave emergence and annihilation, respectively. Further, in the third round of the wave, a pair of emerging and annihilating points disappeared; the wave direction reverted between these points. Artifact signals caused by overlapping tubules are marked by white double daggers in (G-I) and masked between dotted lines in (H-I). Scale bar: 200 µm in (G).

While observations and measures emphasize the wave-like behavior, it is important to recognize that what propagates is only the “phase” of the spermatogenic cycle –or regions where cells are in particular differentiation stages–, but not the constituent cells. In particular, although the live-imaging has shown that, in addition to Stra8 ^+^ spermatogonia, GFPRa1^+^ and Ngn3^+^ A_undiff_ actively migrate within the seminiferous tubules^36,43^ (Figure 4D; Video S6), their migration is limited within local ranges and does not travel over a long distance to create the spermatogenic wave. In the language of dynamical systems, these hallmark properties identify the spermatogenic wave as a *phase wave*. Therefore, to quantitatively characterize such dynamics, the GFP signal intensity profile was converted into phase values ranging from 0 to 2π (Figure 5F).

From these measurements, we found that the phase waves of spermatogenesis showed significant irregularity. In some regions, waves emerge and propagate in opposite directions while, in others, two waves collide from opposite directions and vanish (Figures 5G-I; Video S9). Such singular points, where the wave direction reverses, are defined as *topological defects* in the parlance of dynamical systems. This feature explains the “modulations” observed histologically in fixed tissue, regions with reversed stage order in short stretches of the tubule.^25,20^

Strikingly, we further found that the wave pattern could change over time; for instance, wave emergence and collision sites could annihilate, causing the local wave direction to reverse and resulting in a unidirectional traveling wave (Figures 5H-I). Altering wave patterns were accompanied by fluctuations in the local cycle period. Whether such wave pattern alterations observed in *ex vivo* culture also occur during homeostasis *in vivo* requires careful future assessment, as *in vivo* live-imaging over multiple cycles remains infeasible. Nevertheless, we will see that the observed temporal variations in the phase-wave pattern, at least *ex vivo*, provides an important clue into the mechanism on which it relies.

### Organ-scale patterning of spermatogenic wave sustains constant sperm production

Having characterized the phase-wave behavior of spermatogenesis in millimeter-scale segments of seminiferous tubules, we then questioned how these dynamics integrate into the 10-20-centimeter-long tubule loops, which connect at both ends to the rete testis (Figure 1C). Live imaging of over the entire length of tubule loops was unfeasible. Therefore, we conducted instead precise spatial mapping of the 12 histologically recognizable stages, using 3D reconstruction from serial sections of a fixed whole testis^26^ (Figures 6A-B; Video S10).

**Figure 6.**
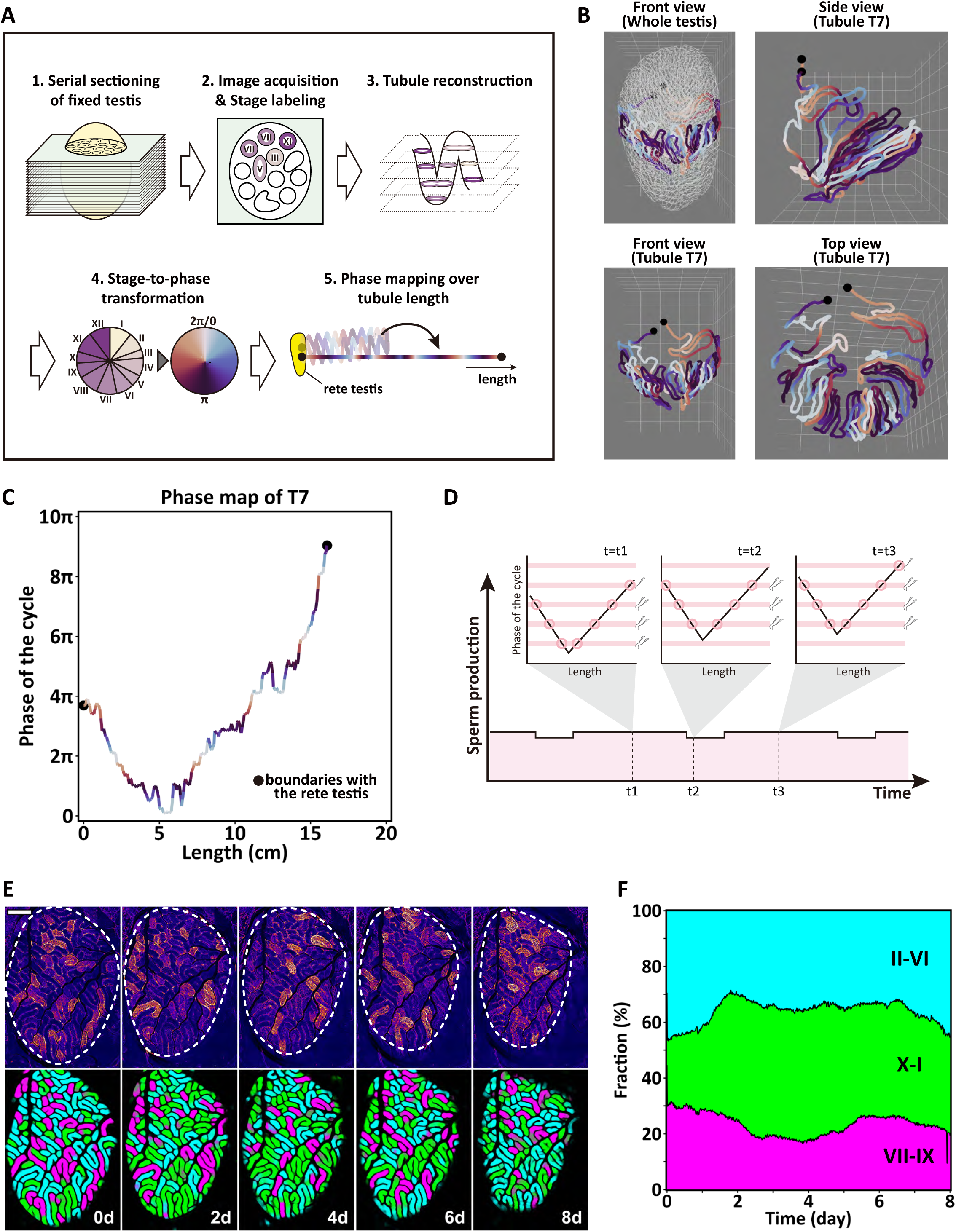
Phase waves of mouse spermatogenesis are patterned macroscopically over the seminiferous tubule loops, underpinning organ-scale homeostasis. **(A-C)** Stage arrangement of the spermatogenic wave over the seminiferous tubule loops. (A) Workflow of higher-resolution reanalysis of testis specimens from Nakata et al. 2017. (B) 3D-reconstructed seminiferous tubules from serial testis sections, highlighting one of the nine tubules (T7) (Video S10). Stages of the spermatogenic cycle (I to XII) were converted to phase information (0-2π) and mapped onto the reconstructed tubules with the color code in (A). (C) Phase map of the spermatogenic cycle over the full length of a tubule loop (T7). Phases are indicated beyond the 0-2π range given the stage continuity; black dots mark the boundaries with rete testis. Note the global V-shape pattern with local jaggedness, representing the descent of segmental order. See Figure S6A for other tubules. **(D)** An explanatory illustration for temporal variation of the summed sperm production (the pink area) from a hypothetical tubule loop showing a definitive V-shape phase profile. Insets indicate the relationship between the phase profile (black V-shape lines) and the phase of spermiation (stage VIII; shown by horizontal pink lines) at different time points (t1, t2, and t3). As the phase profile moves upwards over time, the sites of spermiation (intersections of the black and pink lines; indicated by pink circles) shift along the tubules. Note that the total sperm production fluctuates minimally, contrasting with non-V-shaped phase profiles (Figure S6B). **(E-F)** Spatial phase distribution and its temporal change over a testis *in vivo*. (E) Selected frames of *in vivo* time-lapse images of the entire surface of a *Stra8-EGFP* mouse testis (upper), categorized into stages: GFP^+^ spermatocytes and spermatogonia in magenta (stages VII-IX), GFP^+^ spermatogonia only in green (stages X-I), and no GFP^+^ cells in cyan (stages II-VI) (lower). See Video S11 for the full time series. Scale bar: 1000 µm. (F) Kinetics of the summed proportion of these three phases over the imaging period.

After the stage information was converted into phase values (0-2π) and plotted along the tubule length, the spermatogenic waves were manifest as oblique lines with jagged features reflecting the irregular features of the spermatogenic wave observed in live-imaging (i.e., the stepwise progression, variable wave velocities, and short stretches of reversed wave direction) (Figure 6C). Notably, at the length scale of the tubule loop, the lines showed a consistent V-shape pattern, with the phase decreasing away from the rete and reversing at some point in the middle of the tubule (Figure 6C). Such jagged V-shaped phase patterns were conserved in all loops measured, including those that branched (Figure S6A). These provide a comprehensive and quantitative characterization of the “descent of segmental order,” updating its qualitative classic descriptions^20^.

The spermatogenic wave is thought to ensure the steady production of sperm over time by averaging the local stage variance.^44,21^ Indeed, formation of the macroscopic V-shaped phase pattern serves to consolidate this averaging effect, stabilizing at any given time the fraction of tubule segments at stage VIII, when spermiation occurs (Figure 6D; Figure S6B). To gain insight into how this patterning translates into gross sperm production at the organ-scale, we analyzed our *in vivo* live-imaging films, which cover the entirety of the observed testis surface, over periods corresponding to a complete cycle. Using a U-Net neural network,^45^ we segmented all recognized tubule images into three phases based on the Stra8-GFP pattern (Figure 6E; Figure S6C; Video S11). The results demonstrated that, while the cycle phase varied from segment to segment, the overall proportion of the three phases remained stable as the cycle progressed (Figure 6F; Figure S6D). In particular, the constant contribution of stages VII to IX (magenta) evidences stable daily sperm production.

These measurements demonstrate collectively how germ cell differentiation is spatiotemporally coordinated across multiple length scales. Specifically, periodic cell turnover is arranged into local phase oscillations within the seminiferous epithelium (spermatogenic cycle) that integrate into spatial phase waves along the seminiferous tubules (spermatogenic wave), and which further organize into macroscopic V-shaped phase patterning over the length of tubule loops (descent of segmental order). Such trans-scale cellular coordination supports homeostatic sperm production.

### A coupled oscillator model predicts spermatogenic wave dynamics

Having captured the dynamic and trans-scale organization of spermatogenesis in seminiferous tubules, we next considered how local phase oscillations become coordinated into the larger-scale wave organization. Such phase ordering could be driven by external cues such as spatial gradients of signaling molecules. Alternatively, it could arise spontaneously as a self-organization phenomenon within the context of a functionally homogeneous tissue—a hypothesis resonant with the preservation of the spermatogenic waves in tubule explants (Figures 2I-K, 5C-I). Therefore, we questioned whether a minimal mathematical model could predict the observed intricate dynamics –the robust emergence of traveling phase waves, their characteristic irregularities, and the macroscopic V-shaped pattern across the tubule loops– without assuming the existence of extrinsic influences.

Given the observed clusters of locally synchronized cells separated by small phase differences along the seminiferous tubules (Figure 4), we modeled the system as a one-dimensional array of phase oscillators with weak coupling between neighbors (Figure 7A; for details of the model, see Supplementary Theory). In this framework, the dynamics take the form of a Sakaguchi-Kuramoto model, a minimal theory widely used in the study of coupled oscillator dynamics.^46,47^ In this framework, the local dynamics of a phase oscillator is given by the kinetic equation

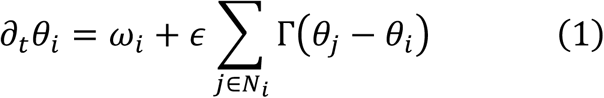

**Figure 7.**
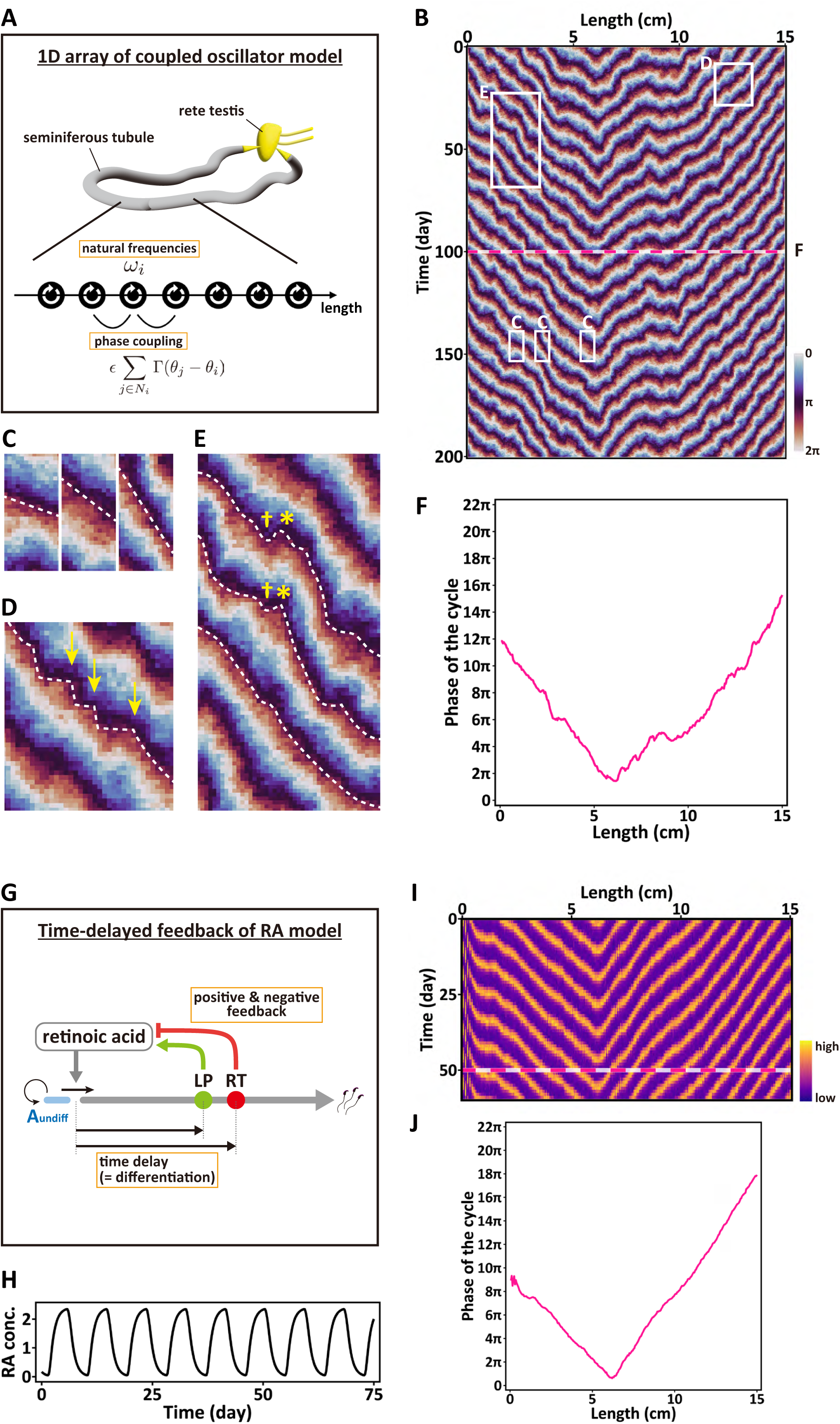
The spermatogenic wave emerges as a self-organization phenomenon based on a model of coupled autonomous oscillators. **(A)** Minimal model of the spermatogenic wave. In this framework, seminiferous tubules are represented as a one-dimensional array of discrete oscillators that interact weakly with their neighbors – a Sakaguchi-Kuramoto-type model. (For details of the model and notation, see main text and Supplementary Theory.) **(B-F)** Stochastic simulation of the model with *η* = −0.44, *σ*/*∈* = 0.09, 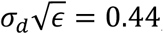, *g* = 1.3, *ω̅* = 2*π*/(8.6 days) and *∈* = 1/(5 days). (B) A representative kymograph of the model dynamics in long-term, color-coded as indicated. (C-E) Magnified images of the rectangular regions highlighted in (B), showing irregular wave patterns similar to that observed in live-imaging observations (Figure 5 and S5), including variant wave velocity (C), abrupt changes in wave speed causing stepwise progression (D, yellow arrows), and defect points where waves emerge and annihilate (E, yellow asterisks and daggers, respectively), as well as the disappearance of these defect pairs (E). (F) A cross-sectional phase profile over the tubule length at time point shown by the magenta dotted line in (B). **(G-J)** Modeling of potential biological processes underpinning the coupled oscillators as the time-delayed feedback of RA metabolism (for details of the model and parameters, see main text and Supplementary Theory). (G) A candidate regulatory framework to explain the origin and robustness of the local autonomous oscillator. A set of RA metabolic enzymes expressed in LP and RT create time-delayed positive and negative feedback loops. (H) Simulation of the model dynamics for RA concentration of a single oscillator in steady state. (I) A representative kymograph of the RA concentration in long term obtained from the oscillator dynamics in (G) when weakly coupled to neighbors in a one-dimensional array with temporal and spatial fluctuations, showing the same features as that observed in the stochastic Sakaguchi-Kuramoto model (A-F). **(J)** A cross-sectional phase profile of local RA oscillators along a tubule loop, converted from the fluctuation of RA concentration in (I), at time point indicated by the magenta dotted line.

where 0 ≤ *θ_i_* < 2*π* denotes the phase of the oscillator at “site” *i* (representing a synchronized cell cluster), Γ(*θ*) = sin *θ* + *η*(1 − cos *θ*) represents the phase coupling between neighboring sites, *j* ∈ *N_i_*, with *∈* its strength, and *η* parameterizing the degree of non-reciprocal phase coupling. Following the classical form of the Sakaguchi-Kuramoto model, we begin by assuming that the natural frequencies of the local oscillators, *ω_i_* = *ω̅* + Λ*_i_*, are variable between sites but unchanging over time, represented by an average frequency *ω̅* with local fluctuations Λ*_i_*. For simplicity, frequency variations between sites, Λ*_i_*, were taken to be statistically independent and drawn at random from a normal distribution with zero average and variance 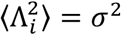. Since the connection with the rete testis likely provides a distinct context that may slow down or speed up the local cycle rate, we imposed a fixed gradient boundary condition at both ends of the oscillator array: Formally, we set *θ*_1_ – *θ*_0_ = *g*_1_ and *θ*_*N*+1_ − *θ*_*N*_ = *g*_2_, where coordinates *n* = 0 and *n* = *N* + 1 index the boundary with the rete and *g*_1_ and *g*_2_ are fixed constants, with *g*_2_ = −*g*_1_ = *g* based on the anatomical symmetry of the two ends of a tubule.

Within this framework, we performed maximum likelihood fitting and found that the model (1) could generate the observed dynamics only partially: While it could reproduce phase waves showing local irregularities and macroscopic V-shaped patterning, we could not find a parameter regime where both features could match quantitatively with the experimental data (Figures S7A-C; Supplementary Theory). However, extending the classic Sakaguchi-Kuramoto model, we found that the spatial patterning and dynamics could be reproduced quantitatively by accounting for temporal variations in the oscillation frequencies. Specifically, setting 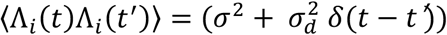, varying the strength of dynamic fluctuations σ_*d*_, the stochastic Sakaguchi-Kuramoto model could reproduce quantitatively the large-scale wave dynamics, patterning and modulations. Significantly, parameter estimation for the measured phase profile yielded consistent values across all nine tubules measured, regardless of their varying loop lengths, indicating homogeneous local properties across and along the seminiferous tubules (see Supplemental Theory for detailed estimation; Figure T10). The ability of the stochastic Sakaguchi-Kuramoto model to generate an irregular V-shape pattern was robust, while the configuration (e.g., the position of the wave direction reversals) depended on the pattern of spatial fluctuations in local frequencies, with minimal sensitivity to the initial phase distribution (Figures 7B-F; Figure S7D-E; Supplemental Theory Figure T11).

The fidelity of the stochastic Sakaguchi-Kuramoto model supports the proposition that seminiferous tubules are functionally uniform, with the characteristic phase-wave dynamics emerging as a self-organization phenomenon without recourse to external cues. The model also provides insights into the origin of the spermatogenic wave and its hallmark irregularities. First, instead of locking to the same phase across the tubule, the formation of persistent traveling waves implicates a mechanism that skillfully adjusts the timing of cell differentiation in response to the stage differences between neighboring sites (i.e., asymmetric phase coupling; Supplemental Theory for detail). Second, local intrinsic variation in the natural cycle period could lead to the nucleation of topological defects at which the wave direction alters. Finally, by constraining the direction and gradient of the phase shift only at the end of the loop, the boundaries with the rete can influence distant regions through asymmetric coupling. This effect and the asymmetric coupling ultimately form the macroscopic V-shape pattern over homogeneous tubule loops.

### Coupled oscillators can be generated by RA-mediated delayed feedback regulation

Based on the validity of the stochastic Sakaguchi-Kuramoto model, we turned to explore the biological processes underpinning the presumed oscillators and their phase-coupling. From a theoretical perspective, feedback regulation with a time delay is a common mechanism for a system to function as an oscillator.^48^ While, experimentally, as described in the Introduction, RA is known to be a crucial player: RA levels vary in synchrony with the spermatogenic cycle, with pulse increases at stages VII-VIII driving the A_undiff_-to-A_1_ transition and other steps of differentiation.^29,31,30^ In addition, rat-to-mouse germ cell transplantation experiment, in which rat germ cells nourished by mouse somatic cells exhibits spermatogenic cycle with a rat tempo, indicates that germ cells are key components that determine the oscillator cycle.^35^

Therefore, we reasoned that the oscillator likely involves a delayed feedback mechanism where both germ cells and RA play critical roles. In this context, we focused on RA metabolism, which involves enzymes expressed in multiple cell types that constitute the seminiferous tubules.^49,17^ Notably, the key enzyme promoting RA synthesis (Aldh1a2) is predominantly expressed in late pachytene spermatocytes (LP), while enzymes that down-regulate RA synthesis by promoting the storage of RA precursor (Adfp and Lrat) are highly expressed in round spermatids (RT). These expression patterns suggest the formation of positive and negative feedback loops between differentiating germ cells and the stem cells, mediated by RA. In this scheme, the time delay is provided by the duration required for A_undiff_ to differentiate into LP and RT, each taking around 17.2 days (2 cycles) and 22.5 days (2.5 cycles), respectively (Figure 7G; Figures S7F; Supplementary Theory Figure T13).

To test whether this hypothesis could explain the dynamics of local oscillation and phase waves observed in the Sakaguchi-Kuramoto-type coupled oscillators, we developed a minimal model for the dynamics of local RA concentration based on delayed positive and negative feedback loops provided by germ cell differentiation, together with a natural decay (see Supplementary Theory for details). Without coupling between neighboring sites, our analysis showed that the model functions as a self-sustained limit-cycle oscillator, where RA concentration varies periodically in concert with the periodic progression of germ cell differentiation, without any external rhythmic input (Figure 7H; Figures S7G-H; Supplementary Theory Figure T14). Interestingly, with the observed time delays for positive and negative feedback loops – 17.2 and 22.5 days, respectively– the model showed an oscillation period half the time delay of the positive feedback, i.e., duration for stem cell to differentiate into LP. This is consistent with the observed period of the spermatogenic cycle, reinforcing the solidity of the model (Figures S7G; Supplementary Theory Figure T14).

Then, by imposing a minimal coupling of RA concentration between neighboring sites, we extended the model to a one-dimensional geometry resembling the seminiferous tubules (see Supplementary Theory for details). Such coupling may arise from mechanisms such as local RA diffusion within the seminiferous tubules, while RA diffusion between neighboring tubules is assumed to be suppressed by an RA inactivating enzyme (Cyp26b1) expressed in peritubular myoid cells.^49,17^ From stochastic simulations, we found that the 1D-extended RA model could reproduce the intricate behaviors of the stochastic Sakaguchi-Kuramoto model, showing irregular phase waves superimposed on a global V-shape pattern (Figures 7I-J; Supplementary Theory Figure T16).

These findings support the hypothesis that the known feedback mechanisms between stem cells and differentiating cells via RA can generate a coupled oscillator, ensuring the robustness of the spermatogenic cycle. Further, through neighborhood coupling, this mechanism also explains the intricate phase wave behaviors along the tubules, recapitulating the dynamics of the stochastic Sakaguchi-Kuramoto model. Our model only defines minimal essential components to capture the critical property of higher-order dynamics. However, at molecular and cellular scales, many other factors must also be involved in proceeding with this feedback mechanism. These include additional RA metabolizing enzymes^49,17^ and other signaling molecules. In addition, Sertoli cells nourish the germ cells and coordinate their differentiation through exquisite interactions within close proximities^18^.

To conclude, our minimal modeling scheme –the stochastic Sakaguchi-Kuramoto model and RA-based feedback model– shows a high capacity for capturing a range of observed intricate dynamics of spermatogenesis across different scales. This finding provides an insight that ordered germ cell behaviors (spermatogenic cycle, wave, and descent of segmental order) can emerge as a self-organization phenomenon rooted in local feedback between stem cells and differentiating germ cells, independent of external instructions or scale-specific mechanisms.

## DISCUSSION

First described over 150 years ago and detailed in the 1950s and 60s, the patterns of spermatogenesis in rodents—namely the cycle, wave, and descent of segmental order—have long fascinated researchers in the field.^8,9,20^ However, insights into the mechanisms that mediate the spatiotemporal dynamics of cellular differentiation across scales have remained limited. Here, to capture the long-term dynamics of mouse spermatogenesis across length scales, we developed ‘ultra’ long-term live-imaging techniques. By linking these observations with precisely mapped macroscopic wave patterns using mathematical modeling, we demonstrated that the intricate dynamics of the spermatogenic wave can be understood as a self-organizing phenomenon, emerging from uniform seminiferous tubules that form coupled oscillators, without the recourse to external guidance cues. We further propose how RA-mediated feedback interactions between stem cells and differentiating cells can generate an autonomous local oscillator and phase wave dynamics along the seminiferous tubules. These results provide mechanistic insights into the spatiotemporal coordination of spermatogenesis. More generally, the current study extends the traditional view that homeostasis relies on tissue stem cells and their niche.^50^ Instead, these results point at a broader paradigm, where stem cells and differentiating cells are tightly coordinated across cell, tissue, and organ scales, facilitating persistent cell turnover through a self-organization mechanism.

Self-organization phenomena in biological systems have been extensively explored, particularly in the context of morphogenesis during development.^51,52^ Specifically, local oscillations and phase wave formation have been observed in processes such as vertebrate somitogenesis, neurogenesis, and fruiting body formation in *Dictyostelium*.^53,54,55,56,57^ These morphogenetic patterns occur on spatiotemporal scales orders of magnitude smaller than those in mouse spermatogenesis. For instance, mouse somitogenesis exhibits a 2-hour cycle and spatial wavelengths of hundreds of micrometers (i.e., tens of cells), forming within millimeter-scale presomitic mesoderm tissue.^58^ In these microscopic dynamic processes, the underlying physical principle and its biological basis have been intensively studied–though still not fully understood. By contrast, the macroscopic coordination of cell dynamics during homeostasis has remained underexplored, primarily due to the challenges posed by their large spatiotemporal scales. The current study highlights that, while the spermatogenic wave may rely on the same class of physical mechanisms as developmental morphogenesis, it exhibits distinct features tailored to the functional requirements for tissue and organ-scale homeostasis.

First, the origin of the massive differences in time and space scales is an intriguing problem. In somitogenesis, individual cells primarily act as oscillators, with intracellular levels of gene products and metabolites fluctuating.^59^ In this scheme, time-delayed feedback regulation is rooted in molecular processes such as transcription, translation, protein maturation, localization, and degradation. By contrast, the current study suggests that, in adult spermatogenesis, a group of cells within a local tissue region, including both stem cells and differentiating cells, function collectively as a phase oscillator. Herein, cell differentiation proceeds periodically, with the levels of intercellular signaling molecules such as RA oscillating. In this system, differentiating cells can mediate feedback coupling by regulating the fate of the stem cells from which they originate, with time delays arising from the duration required for stem cells to differentiate into certain stages. This lengthy process explains the characteristically long 9-day rhythm. These findings highlight how the same physical mechanism can operate across different biological hierarchies, functioning on diverse spatiotemporal scales and serving distinct roles—in this case, morphogenesis and homeostasis.

Second, building on the previous point, it is worth emphasizing that while phase waves in developmental patterning are transient, the adult spermatogenic wave is persistent. In the former, phase waves provide spatiotemporal cues for progenitor cells to differentiate irreversibly into specific cell types. At this point, oscillation and phase waves terminate, fixing the spatial pattern. However, in adult spermatogenesis, although the phase waves similarly dictate the differentiation of stem cells, these cells robustly maintain a self-renewing compartment that produces differentiating cells in subsequent cycles if exposed by RA.^60,2^ Consequently, the oscillations and traveling phase waves do not terminate but persist. This difference underscores their distinct biological functions: phase waves in developmental patterning imprint a blueprint for future structure while, in homeostasis, they sustain long-term cell turnover and sperm production.

Finally, it is important to note that spermatogenic waves exhibit significant irregularities in their spatiotemporal patterns. This feature contrasts with developmental morphogenesis, where phase wave dynamics typically generate spatially ordered patterns from unstructured conditions, as seen in somitogenesis from the presomitic mesoderm. Indeed, this spatial irregularity in spermatogenesis becomes even more pronounced in non-rodent species like humans and birds, where the spatial arrangements within the seminiferous tubules are more intricate and less organized, despite being based on similarly stereotypic local cycles.^61,62,63,64,65^ While the mechanisms underlying the species difference remains to be explored, this suggests that, instead of generating spatially regular patterns, phase waves in mouse spermatogenesis serve two key functions in homeostasis: maintaining local phase coherence and facilitating phase dispersion over greater length scales. Microscopically, phase coupling minimizes gaps in differentiation timing between neighboring sites. This mechanism effectively prevents stage-specific signaling molecules from diffusing into adjacent regions at distant stages of the cycle, or a single Sertoli cell from nurturing germ cells at distinct stages. Macroscopically, the formation of phase waves effectively disperses stages of the spermatogenic cycle, which is further averaged by the V-shape pattern over the tubule loops. This property underpins organ-scale homeostasis and stable sperm production.

Mouse spermatogenesis is an ideal system for characterizing tissue-to-organ scale patterning, thanks to the dramatic changes in cell morphology during differentiation and the prominent local synchrony resulting from syncytia formation. However, such macroscopic patterns, if present, might remain unrecognized in other cycling tissues and organs during adult homeostasis. A close analogy can be drawn with the hair follicle, which undergoes periodic bouts of growth and regression, with cycles extending over a couple of months (the hair cycle).^66^ Reminiscent of spermatogenesis, hair cycles can become globally desynchronized under regenerative conditions and certain genetic backgrounds, leading to phase wave-like propagation.^67,68,69^ It remains a vital question for future research whether the dynamics of coupled oscillators caused by feedback interplay between stem cells and differentiating cells, or other self-organization mechanisms, are employed in the regulation of stable homeostasis in skin or other organ contexts over similarly extended spatiotemporal scales. Equally, it will be intriguing to see how these mechanisms may alter in non-homeostatic contexts such as growth, aging, regeneration, and disordered states. Trans-scale live imaging technologies may provide essential insights for addressing these questions and advancing our understanding of a wide range of biological processes.

### Limitation of the study

The *ex vivo* live-imaging studies conducted in this study used juvenile, rather than adult, testis explants as source material, due to current technical limitations. In addition, as is generally true for organ culture, the integrity of the explant tissue is somewhat compromised. These limitations required careful interpretations of the results beyond concluding that the spermatogenic cycle and wave can occur and propagate. Thus, we verified the wave properties observed *ex vivo* by consistency with histological or *in vivo* imaging observations whenever possible (e.g., stepwise wave progression or reversed wave direction). While particular care must be taken in interpreting the observed wave pattern alteration over multiple cycles, currently such effects cannot be verified *in vivo*.

Our RA-based minimal model, relying on just a few experimentally-demonstrated essential components (i.e., RA and germ cell differentiation), effectively explains the oscillatory dynamics of adult homeostatic spermatogenesis. While these results suggest that this model can capture the core oscillation mechanism, other components are also involved and play auxiliary and redundant roles. These include RA actions and metabolism not described in the model explicitly, extracellular signals other than RA, and somatic cells such as Sertoli cells. Further, while focusing on cell turnover during homeostasis, our model does not account for other contexts in which the spermatogenic cycle is established from conditions where differentiating germ cells (e.g., LP and RT) are absent, including the first-wave of spermatogenesis in postnatal mice or regeneration following insult or transplantation. Understanding the dynamics in these non-homeostatic contexts requires future experiments and potentially extensions of the model.

## Acknowledgments

We are grateful to Yumiko Saga for providing the *Stra8-EGFP* mice and Kei-ichiro Ishiguro for anti-Stra8 antibody. We thank Kagayaki Kato and Yoshitaka Kimori for suggestions on image analysis, and the current and former Yoshida Lab members, including Yoshiaki Nakamura for technical instructions and fruitful discussions. We acknowledge the support of the Model Organisms Facility (NIBB Trans-Scale Biology Center) for animal care and the Equipment Development Center, the Institute for Molecular Science for culture device manufacturing.

## Author contributions

T.S. and S.Y. designed the research framework; T.S. improved *in vivo* live-imaging settings and performed all *in vivo* and *ex vivo* live-imaging experiments, immunofluorescence experiments, image processing, and quantification; T.S., M.K., H.Y., K.H., and T.O. developed the *ex vivo* organ culture method for imaging, with H.Nakamura and H.K. designing and fabricating culture devices. T.S., Y.K., and K.A. performed image analysis using machine learning; H.Nakata and T.S. conducted fine stage mapping of reconstituted seminiferous tubules; Y.I.L., D.J.J., and B.D.S. performed theoretical model analyses. S.Y., M.T., and B.D.S. supervised the project. T.S., Y.I.L., B.D.S., and S.Y. wrote the manuscript with input from all other authors.

## Methods

### Animals

*Stra8-EGFP* mouse strain (ICR.Cg-Tg(Stra8-EGFP)1Ysa/YsaRbrc, RBRC02498) was developed by Yumiko Saga (National Institute of Genetics, Japan) and provided by the RIKEN BRC through the National BioResource Project of the MEXT/AMED, Japan.^38^ *Stra8-EGFP* mice maintained in the ICR background were backcrossed to C57BL/6J females (Japan SLC or CLEA Japan) for four or more generations before experiments, with the transgene most likely located on Y chromosome. C57BL/6J adult male mice were purchased from Japan SLC or CLEA Japan. All animal experiments but that described below were conducted with approval by the Institutional Animal Care and Use Committee of the National Institutes of Natural Sciences.

For detailed stage mapping of the spermatogenic cycle on 3D-reconstructed seminiferous tubules, we reanalyzed the image data of serial sections of a whole testis of C57BL/6N mouse (purchased from Japan SLC) described previously,^26^ in which animal experiments were conducted with approval by Kanazawa University and conducted in accordance with the Guidelines for the Care and Use of Laboratory Animals in Kanazawa University.

### Immunofluorescence (IF)

#### Testis sections

Wild-type (C57BL/6J) and *Stra8-EGFP* adult mice (3-4 months old) were sacrificed by cervical dislocation, whose testes were removed and fixed overnight in 4% PFA in PBS at 4°C, embedded in paraffin, and sectioned at 5 μm thickness. After deparaffinization, rehydration, and antigen retrieval by heating at 98°C for 15 min in 20 mM Tris–HCl (pH 9.0), sections were blocked with Can Get Signal™ solution 1 containing 4% donkey serum and Blocking One Histo (1 drop per 500 μL) for 1h at room temperature (RT), followed by incubation with the primary antibodies diluted with the same blocking solution (3h, RT). Then, sections were washed with PBS containing 0.04% Tween20 (PBST) and incubated with secondary antibodies, Hoechst 33342, and fluorescent-labeled PNA-lectin diluted with Can Get Signal™ solution 2 containing 4% donkey serum and Blocking One Histo (1 drop per 500 μL) (2h, RT). PNA-lectin was fluorescent-labeled by mixing PNA-lectin (10 mg/mL) and Alexa Fluor 750 NHS Ester (10 mg/mL) in 200 μL H_2_0 and 20 μL DMSO. To additionally stain GFP in *Stra8-EGFP* specimens, the above procedure was followed by washing with PBST, incubation with Can Get Signal™ solution 1 containing 4% goat serum and Blocking One Histo (1 drop per 500 μL) (1h, RT), and that containing Alexa Fluor 647-labelled anti-GFP antibody (2h, RT). Finally, sections were washed with PBST and mounted with PermaFluor Aqueous Mounting Medium for fluorescence microscopy.

#### Whole-mount seminiferous tubules

Whole-mount IF was conducted as previously described^70^ after some modifications. Untangled seminiferous tubules from *Stra8-EGFP* were fixed in 4% PFA in PBS (2h, 4°C), dehydrated and rehydrated through graded methanol in PBST, washed in PBST, and incubated with PBST containing 4% donkey serum and Blocking One Histo (1h, RT). Then, tubules were incubated with primary and secondary antibodies (2h each, RT), each followed by washing with PBST. Hoechst33342 was also contained in the incubation buffer for secondary antibody reaction. The specimens were further stained for GFP, through blocking with 4% goat serum and Blocking One Histo (1 drop per 500 μL) in PBST (1h, RT), incubation with Alexa Fluor 647-conjugated anti-GFP antibody (2h, RT), and wash with PBST. The specimens were mounted with Fluoro-KEEPER Antifade Reagent for fluorescence microscopy.

#### Image acquisition

IF-stained section and whole-mount specimens were photographed using an upright laser scanning confocal microscope equipped with a motorized stage (TCS SP8, Leica Microsystems), through a 20x oil immersion objective lens (HC PL APO 20x/0.75 IMM CORR CS2, #506343, Leica Microsystems). Tiled images were stitched together to generate a composite image of the specimen using LAS X software (Leica Microsystems). In the case of section specimens, given the arrangement of detectors, Alexa Fluor 750, Alexa Fluor 647, and Alexa Fluor Plus 555 signals were first acquired sequentially, followed by another sequential acquisition of Alexa Fluor 488, Brilliant Violet 480, and Hoechst 33342 signals. Bright-field images of whole-mount samples were taken with a BX51 upright wide-field microscope (Olympus) with a 4x objective lens (UPlanSApo 4x, Olympus) using cellSens software (Olympus), and stitched using Fiji software.^71^

### *in vivo* (intravital) live imaging

#### Surgery and anesthesia

A previously described setting^36^ was modified in the current study to achieve up to 9 days-long *in vivo* live imaging. In brief, three-month-old male *Stra8-EGFP* mice were anesthetized by intraperitoneally administrating a mixture of 0.75 mg/kg medetomidine, 4 mg/kg midazolam, and 5 mg/kg butorphanol (MMB). After removing hair from the abdomen and neck using a hair-removing cream, a testicle was gently pulled to locate outside the abdominal wall with the associated gubernaculum cut, while blood vessels were kept intact. The mouse was held in a holder; the pulled-out testicle was placed in a custom-made stainless-steel chamber whose ceiling was composed of a cover glass. The chamber was kept in a constant temperature (32.5°C) environment using attached heat plates (Tokai Hit). For prolonged filming, anesthesia was switched from MMB, which lasts for approximately 6h, to isoflurane inhalation (0.5% or lower in oxygen-concentrated air) using a TK-7 evaporator (Biomachinery) and a TK-5A oxygen concentrator (Taiyo Electronic). In addition, to reduce the dose of isoflurane and its respiratory depressant effect, continuous intraperitoneal administration of anesthetics was combined as described below.

Water containing nutrients (i.e., carbohydrates, amino acids, lipids, vitamins, minerals, and trace elements) were delivered by intraperitoneal infusion of 4:1 mixture of Hicaliq NC-H (Terumo) and Aminoleban (Otsuka Pharmaceutical) via an Indwelling tubing (Hibiki polyethylene tubing No4, Sansyo) and by enteral nutrition (Peptamen AF, Nestlé Health Science) via a gastric tube (SP tube SP10, Natsume Seisakusho). Using KDS200 syringe pumps (KD Scientific), intraperitoneal and enteric infusions were adjusted between 10-40 μL/h and 20-80 μL/h, respectively. The intraperitoneal infusion also contained 0.4 μg/mL dexmedetomidine (Maruishi Pharmaceutical), 75μg/mL butorphanol (Meiji Seika Pharma), and 1% penicillin and streptomycin solution (Thermo Fisher Scientific).

Heart rate, respiratory rate, and percutaneous oxygen saturation were continuously monitored in the throat using a MouseOx PLUS vital monitor (STARR LifeSciences). Doses of isoflurane, oxygen, and intraperitoneal and enteral infusions were adjusted based on the vital signs, respiratory movement, and defecation during the imaging period. An eye ointment (white petrolatum) was applied to prevent dryness. When imaging was completed, mice were euthanized by deep anaesthetization with a high concentration of isoflurane (5%) followed by cervical dislocation; filming was terminated in case the mice died or emaciated based on their appearance and vital signs, as well as if the observed testis was damaged typically due to impaired blood supply.

#### Image acquisition

Anesthetized mouse in a holder was placed on a BX61WI upright microscope (Olympus) equipped with an MD-XY30100T-META motorized XY stage (Sigmakoki) and an iXon+ DU-888 EMCCD camera (Andor), so that the surface of *Stra8-GFP* mouse testis was imaged with an epifluorescence optics using a metal halide light source (U-PS50MH, Olympus) and a 10x dry or 30x silicone immersion objectives (UPlanApo10x or UPLSAPO30XS, Olympus). For time-lapse imaging, multipoint tiled images with Z-series (9 images per position with 2 μm interval) were acquired every 20 minutes. The imaging system was controlled by Metamorph software (Molecular Devices).

### Ex vivo imaging

#### Culture device

*ex vivo* testicular organ culture method used for fluorescent time-lapse imaging was developed based on previously reported microfluidic device made of gas-permeable material, polydimethylsiloxane (PDMS).^37^ The culture chamber consisted of an upper layer of medium channel (2mm width and 400 μm height) and a lower layer of tissue chamber (3 mm width, 2 mm length, and 160 μm height), separated by a porous polycarbonate membrane (Isopore™ membrane filter with 10 μm pore size and 5–20% porosity, Millipore). To prepare the upper and lower layers of the culture device, PDMS and curing reagent were mixed at the 10:1 ratio, poured into a photolithographically fabricated SU-8 master mold, and cured at 70°C for >2h. Then, the upper and lower PDMS parts, activated by oxygen plasma with a PDC-32G plasma cleaner (HARRICK), and a porous membrane, which were plasma-activated and silanized using an amino silane, were bonded together and cut into a circular shape (ϕ30 mm) to fit a fixture for microscopy. A ϕ2×ϕ3 mm silicone tubing (6-586-05-01, AS ONE) and two ϕ0.5×ϕ1 silicone tubings (6-586-01-01, AS ONE) were connected to the tissue inlet and the medium inlet/outlet of the device, respectively. A culture medium bag, made of a plastic bag (Unipac B4-ST, Seisan Nipponsha) attached with a 30G needle (NO.30, Dentronics), was connected to the medium inlet of the culture device via a polyethylene tubing (SP-10, Natsume Seisakusho), while a 5 mL syringe (SS-05SZ, Terumo) with a 30G needle was connected to the medium outlet to be pulled by a Legato111 syringe pump (KD Scientific). Devices were sterilized by UV irradiation for at least 15 minutes, immediately before use.

#### Explant culture

*Stra8-EGFP* mice at 10- or 11-days post-partum were deeply anesthetized with isoflurane and sacrificed by cervical dislocation, whose testes were harvested and immersed in a culture medium (α-MEM containing 10v/v% KnockOut Serum Replacement and 1v/v% Penicillin-Streptomycin) and removed of the tunica albuginea with the seminiferous tubules mildly untangled with forceps. Then, some 1/10 of the total tubules of a testis were introduced into a culture device filled with the medium above, through the tissue inlet. The culture device was incubated in a STX-IX3WX stage-top incubator (Tokai Hit) attached to an IX83 inverted microscope (Olympus) equipped with a BIOS202T motorized XY stage (Sigmakoki), or in a CV1000 confocal scanner system (Yokogawa Electric), and kept in a 34°C and 5% CO_2_ condition. The culture medium was perfused through the device at a constant flow rate of 0.2 μL/min using a syringe pump from the culture medium bag, which was kept at <10°C in a D-CUBE S small refrigerator (Twinbird) and refilled with fresh medium every two weeks.

#### Image acquisition

Seminiferous tubule explant in the culture device incubated on an IX83 inverted microscope was imaged using a CSU-W1 spinning disk confocal scanner unit (Yokogawa Electric) and a Prime95B sCMOS camera (Photometrics) using the denoising mode of the camera. For time-lapse imaging, multipoint tiled images with Z-series (31 images with 7 μm intervals for each position) were acquired every 20 min using a 10x UPlanSApo objective lens (Olympus), controlled by Metamorph software (Molecular Devices). In some cases, images were acquired in the same condition using CV1000.

### Image processing

Most image processing, including movie construction, was conducted using Fiji software unless otherwise described.

#### IF images

To prepare IF images of testis sections (Figure 2B; Figures S2A-F), regions of seminiferous tubules at particular stages were cropped to create montages. For whole-mount IF images (Figure 2C; Figures S2G-J), seminiferous tubule regions were cropped based on the GFP patterns. Selected color channels were shown in pseudo-colors, with brightness adjusted for enhanced visibility. In Figure S2G, the low-magnification whole-mount IF image was prepared by downsizing the original image (bin=8, method=average); the bright-field image was rotated and resized to match the IF image (interpolation=Bilinear) with brightness adjusted for improved visibility.

#### *in vivo* (intravital) imaging data

Images obtained using a 10x objective (Figures 2E-G, 3A, 4A, 4D, 5A, and 6E; Figure S3A; Videos S1, S2, S4, S6, S7, and S11) were converted from 16 to 32-bit format, followed by reducing shot noise and background using a median filter (radius=1 pixel) and Subtract background plugin (radius=10 pixels). Then, Find focused slices plugin developed by Qingzong Tseng (https://sites.google.com/site/qingzongtseng/find-focus) was utilized to identify blurred images (images with >50% of maximum variance, variance threshold of 0, apply an edge filter, pick up only slices consecutive) and eliminate them from the Z-series. Subsequently, depth information was integrated using Z-projection (Average Intensity). Further, to reduce the image curvature aberration, 10% of the view margins were trimmed and stitched in the XY direction from multiple images using the BigStitcher plugin.^72^ After the images were converted to 16-bit format, brightness was adjusted along the time direction using the Histogram Matching algorithm in Bleach correction plugin,^73^ given that the composition of the stages of spermatogenesis was almost unchanged over time (see main text and Figures 6E-F). Pseudo-color setting and brightness adjustment were applied for enhanced visibility. The processed data were subjected to movie construction, signal intensity quantification, cell tracking, kymograph generation, and semantic segmentation by machine learning (see below).

As for images acquired with a 30x objective (Figure S4A; Video S5), the best-focused slices were visually selected from the Z-series, registrated with the rigid transformation of StackReg plugin,^74^ and shot noise was removed with a median filter (radius=1 pixel). Brightness was temporally adjusted using the Simple Ratio algorithm of the Bleach correction plugin (background=700). After converting to 32-bit format, background was removed using Subtract background plugin (radius=100 pixels). Brightness was adjusted to enhance the visibility.

#### Semantic segmentation of *in vivo* imaging data

To conduct segmentation of the seminiferous tubule images based on the stages of the spermatogenic cycle, we used U-Net,^45^ one of the most well-tested deep learning models for semantic segmentation of biomedical images. For preparing data for annotation, 18 images of the entire testis surface selected from six compiled movies, following preprocessing procedures described above in the “*in vivo* (intravital) imaging data” section, were manually annotated into the tubule regions of stages II-VI (where no GFP signal was observed), VII-IX (where prominent GFP signal was visible in spermatocytes) and X-I (where signal in spermatocytes weakened, making GFP^+^ spermatogonia clearly recognizable), and off-tubule regions including interstitium and blood vessels. From each of 15 out of these 18 manually annotated images, we created 3,000 derivative images (768 x 768 pixels) through random cropping, rotation, flipping, and z-score normalization, summing up to 45,000 teacher images. From the remaining three manually annotated images, a dataset of 900 images was generated for validation purpose, by creating 300 derivative images from each image using the same procedures of random cropping, rotation, flipping, and z-score normalization.

We adopted the Keras framework (Tensorflow backend) to train the U-Net architecture on our data, with its original parameter size (30M) reduced to 3.5M for computational efficiency (see Figure S6C for details). Starting from randomly initialized weights, the Adam optimizer (learning rate: 1e-3, batch size: 8) was used to minimize the categorical cross-entropy loss. We divided the 45,000 training images into five subsets for retrospective assessment of their heterogeneity and conducted five successive training rounds (10 epochs each) using these subsets. The model showing the lowest validation loss in each round, calculated using a validation dataset described above, was transferred to the next round. The validation loss showed similar values across the five rounds, suggesting that the model was effectively trained based on unbiased teacher datasets. The model with the lowest validation loss in the final round was used for further analysis.

For segmentation of the time-lapse movies in full length, one thousand image stacks were generated in randomly cropped regions of 768×768 pixels. After Z-score normalization, each cropped image was processed by the trained U-Net, and the segmented images were averaged to produce the segmented movies (Figure 6E; Video S11).

#### *ex vivo* imaging data

After conversion to a 32-bit format, the acquired images were processed by an adaptive median filter (radius=7 pixels, threshold=3), an originally implemented algorithm for fast and efficient shot noise removal inspired by a method available at https://imagej.net/plugins/adaptive-median-filter, and a median filter (radius=1 pixel). Then, depth information was integrated using Z-projection (Max Intensity). After trimming 10% of the field edges (not applied to data acquired using CV1000), images were stitched in the XY direction using BigStitcher plugin. These processed data were used to quantify GFP signal intensity, generate kymographs (see below). In generating movies, only for better visualization, after converting to a 16-bit format, the intensity was adjusted in the time direction using Bleach correction plugin (Simple Ratio; background=20410 and 3850 for images acquired by IX83 and CV1000, respectively), followed by pseudo-color and intensity adjustment (Figures 2I-J, 5C-D, and 5G; Figure S3B; Videos S3, S8, and S9).

### Image data assessment and quantification

#### Germ cell types and stage of seminiferous tubules

In IF-stained testicular sections (Figure 2B; Figures S2A-F), undifferentiated spermatogonia (A_undiff_), differentiating spermatogonia, early and late spermatocytes are defined visually as PLZF^+^/KIT^−^/SYCP3^−^, PLZF^+/-^/KIT^+^/SYCP3^−^, PLZF^−^/KIT^+^/SYCP3^+^ and PLZF^−^/KIT^−^/SYCP3^+^ cells, respectively. Although our multi-color detection setting inevitably causes weak leakage of PLZF signal using Brilliant Violet 480 into Alexa Fluor 488 channel used for KIT detection, we reason that the true KIT signal is observed on the cell surface membrane in contrast to the nuclear localization of PLZF. Staging of seminiferous tubules cross-sections (from stages I to XII) was conducted according to the established morphological criteria based on constituent cells’ nuclear morphology, acrosomal system visualized by PNA-lectin staining, and their association.^10,1,75,76,77^

In whole-mount IF images of Stra8-GFP seminiferous tubules (Figure 2C; Figures S2G-J), differentiating spermatogonia and early spermatocytes are defined as GFP^+^/KIT^+^/SYCP3^−^ and GFP^+^/KIT^+^/SYCP3^+^ cells, respectively. These cells’ stage dependence, determined by transillumination microscopy,^78,79,80^ agreed with that based on testis sections as described above.

As for live-imaging data, Stra8 ^+^ differentiating spermatogonia and early spermatocytes was identified as GFP^+^ cells showing consistent morphology with those observed in whole-mount IF.

#### Kinetics of Stra8-GFP signal intensity

To quantify the temporal changes in the GFP signal intensity within a specific region of the *ex vivo* live-imaging data set processed as described above (Figure 2K), using Fiji software, we manually defined a rectangle ROI of 50×50 pixels at virtually the exact position of growing and deforming seminiferous tubules over time, measured its average signal intensity for each frame. Using R, the obtained time-series data were processed by Min-Max normalization (normalize function of BBmisc package) and Savitzky-Golay smoothing (sgolayfilt function of signal package; filter order=4, filter length=51, 0-th derivative), and visualized using the line plot procedure (ggplot2 package).

For *in vivo* time-lapse imaging data (Figure 4B-C), the target seminiferous tubule was enlarged 16-fold in length using Fiji software (interpolation method =Bilinear), and tiled systematically with ROIs of 309×309 pixels (50×50 μm), and the average intensity within each ROI was extracted for every frame. After conducting Min-Max normalization and Savitzky-Golay smoothing (filter order=4, filter length=31, 0-th derivative) using R, the kinetics in signal intensity for each ROI were hierarchically clustered, using the Euclidean distance between the intensity profiles as the dissimilarity structure, by means of hclust function of stats package (method = ward.D2). The cluster number was adjusted to be visually interpretable. The result of clustering was visualized as a dendrogram using ggplot2 and dendextend packages, while the ridge plots of corresponding ROIs’ intensity profiles were generated using ggplot2 and ggridges packages.

#### Motion tracking of Stra8-GFP^+^ spermatogonia

GFP^+^ spermatogonia live-imaged *in vivo* were tracked using the semi-automatic tracking tool of TrackMate plugin^81^, Fiji (Figure 4D). Then, using R, the XY coordinate and time information of all tracked cells (i.e., spots with a track ID) were linked with the syncytium ID. Syncytium IDs were provided by subjective identification of individual syncytia based on the short distance and coherent migration shown by cells belonging to the same syncytium and the reported length distribution of syncytia upon A_undiff_-to-A_1_ transition.^82,83^ Cell division events were detected as branching points of each track. From these measurements, we used Fiji software to create a tracking movie (Video S6), project the complete trajectories into a single panel (Figure S4B), and visualize the position/timing of the first observed divisions of Stra8-GFP^+^ spermatogonia, i.e., A_1_-to-A_2_ transition (Figure 4F). We also generated a genealogical chart using R’s ggplot2 package, aligned in a positional order of their trajectories’ gravity center (Figure 4E).

#### Kymographs of spermatogenic wave

For *in vivo* imaging data, using the StackReg plugin of Fiji, images of seminiferous tubule segments of interest were registrated, and polyline ROIs were drawn manually along the tubule center (Figure 5A). Then, using KymoResliceWide v.0.5 plugin (https://github.com/ekatrukha/KymoResliceWide), kymographs of the GFP signal intensity along the tubule length were generated by averaging the signal intensity over 25 pixels-width along the ROIs, and rescaled without interpolation to the resolution of 1 μm x 1 min per pixel (Figure 5B; Figures S5A-C).

For *ex vivo* imaging, in which the tubules changed in shape and length during imaging periods, polyline ROIs were drawn manually at certain frames of the movie (Figures 5C and 5G) and interpolated to other time points, to generate linearized tubule images for Figure 5D, movies (Videos S8-9), and kymographs averaged over a 25-pixel width across the ROI (Figures 5E and 5H), using LOI Interpolator of Time Lapse plugin.^84^

To convert the obtained intensity kymographs of the GFP signal into phase maps of the spermatogenic cycle, wavelet conversion was applied within a Python environment, as detailed below.^84^ First, changing tubule lengths were normalized to that in the time point showing the maximum length, by linearly interpolating and resampling the intensity data at each timepoint (scipy.interpolate.interp1d class), and applied for Savitzky-Golay smoothing over the length (scipy.signal.savgol_filter function; filter order=4, filter length=101, 0-th derivative). In such length-normalized intensity kymographs, time-series intensity data at each spatial point was transformed into a continuous wavelet using a complex Morlet wavelet as the mother wavelet (scipy.signal.morlet2 function), from which a phase time series (0 to 2π) was obtained as the angle of the complex argument (numpy.angle function). The wavelet scales of 550 and 500 were used for Figures 5F and 5I, respectively. The phase was shifted so that stage I starts approximately at 0/2π, based on the fact that the highest GFP intensity, which points π in the algorithm used, occurs around stage VIII. Finally, for each time point, the linear spatial phase map was converted back to its original length scale, using the same methods of linear interpolation and resampling. The resultant 2-dimensional phase maps were presented at a resolution of 1 μm and 5 min per pixel, using Fiji (interpolation=None).

#### Contributions of the phases of spermatogenic cycle segmented by U-Net

To quantitate over-time contributions of tubule regions in stages VII-IX (magenta), X-I (green), and II-VI (cyan), the U-Net derived images were binarized using Li’s Minimum Cross Entropy Thresholding method of Fiji. Each channel of phase was calculated so that these and off-tubule (black) regions become mutually exclusive, and the area of each phase at all time points was obtained. Then, using R, the proportion of the three phases was calculated from the above time-series data and visualized on a percent stacked area chart using the ggplot2 package (Figure 6F; Figure S6D).

### Stage-mapping on 3D reconstructed seminiferous tubules

#### Whole testis reconstruction

A microscopic digital image dataset of whole-testis serial sections obtained in our previous study^26^ was reanalyzed in the current study. In that previous study, a Bouin-fixed, paraffin-embedded testis from a 90-day-old C57BL/6N male mouse was serially sectioned with 5 μm thickness and collected every 50 μm.

After PAS-H (periodic acid Schiff and hematoxylin) staining, all sections were photographed and digitized using a whole slide scanner (Nanozoomer 2.0-HT; Hamamatsu Photonics). In the current study, 3D reconstruction was performed as previously described^26^ with slight modifications. Namely, the slide images were converted into JPEG format using Photoshop software (Adobe). Then, using Amira software (Thermo Fisher Scientific), all seminiferous tubules were segmented, aligned, and reconstructed in 3D, whose core lines were extracted at a resolution of 10 μm x10 μm x10 μm per voxel.

#### Stage mapping on seminiferous tubules

For stage mapping on the 3D-reconstructed tubules, the stage of each tubule cross-section was determined histologically based on the established criteria,^1^ in which stages II and III were not classified. Voxels on the core lines of the reconstructed tubules (described previously) were labeled with the stage information by providing an intensity value ranging from 1 to 12 (stage II and III are collectively indicated as 3), using Fiji.

Then, the XYZ coordinates and the stage information of the voxels (10 μm x10 μm x10 μm each) on the core lines were extracted. To remove shot noises, most of which are likely due to staging errors, a rolling median protocol was applied to stages in 5 voxels, using rollapply function of zoo package of R, while missing stage information were complemented using those in neighboring regions. Then, the stage was converted to phase value (0 to 2π) based on the frequency of stages measured in testis sections.^11^ Specifically, we set the onset of stage I to be at the phase 0 (precisely, integer multiples of 2π), with all data points having the phase value of the very middle of the duration of the particular stage. For example, all points that are in stage I, which consist of 10.73% of all the tubule sections, were converted to a phase value of 2π x 0.1073/2. To avoid the gap between 0 and 2π phases, appropriate multiples of 2π were added using the unwrap function of signal package.

For three-dimensional stage visualization upon the seminiferous tubules (Figure 6B, Video S10), phase information of the voxels on the core lines were further smoothened by Savitzky-Golay method (sgolayfilt function of signal package; filter order=4, filter length=151, 0-th derivative) and visualized as a 3D line plot using plotly package of R.

For one-dimensional stage mapping along the tubule lengths (Figure 6C; Figure S6A), phase information on the core lines were phase-averaged every 50 voxels for smoothing. Additionally, the distance of each voxel from the boundary connecting to the rete testis along the tubules’ core lines was calculated from the XYZ coordinates of the voxels by cumulatively summing the linear distances between adjacent voxels’ centers (10, 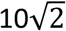, or 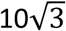 μm for immediately adjacent voxels, diagonally adjacent voxels, or diagonally adjacent voxels in a different direction, respectively). Due to microscopic zigzags, the distances measured in this manner became some 10% longer than previously reported values using Amira software.^26^ Finally, phase distribution along the tubule lengths was visualized as a line plot using ggplot2 package.

---

**Figure S1.**
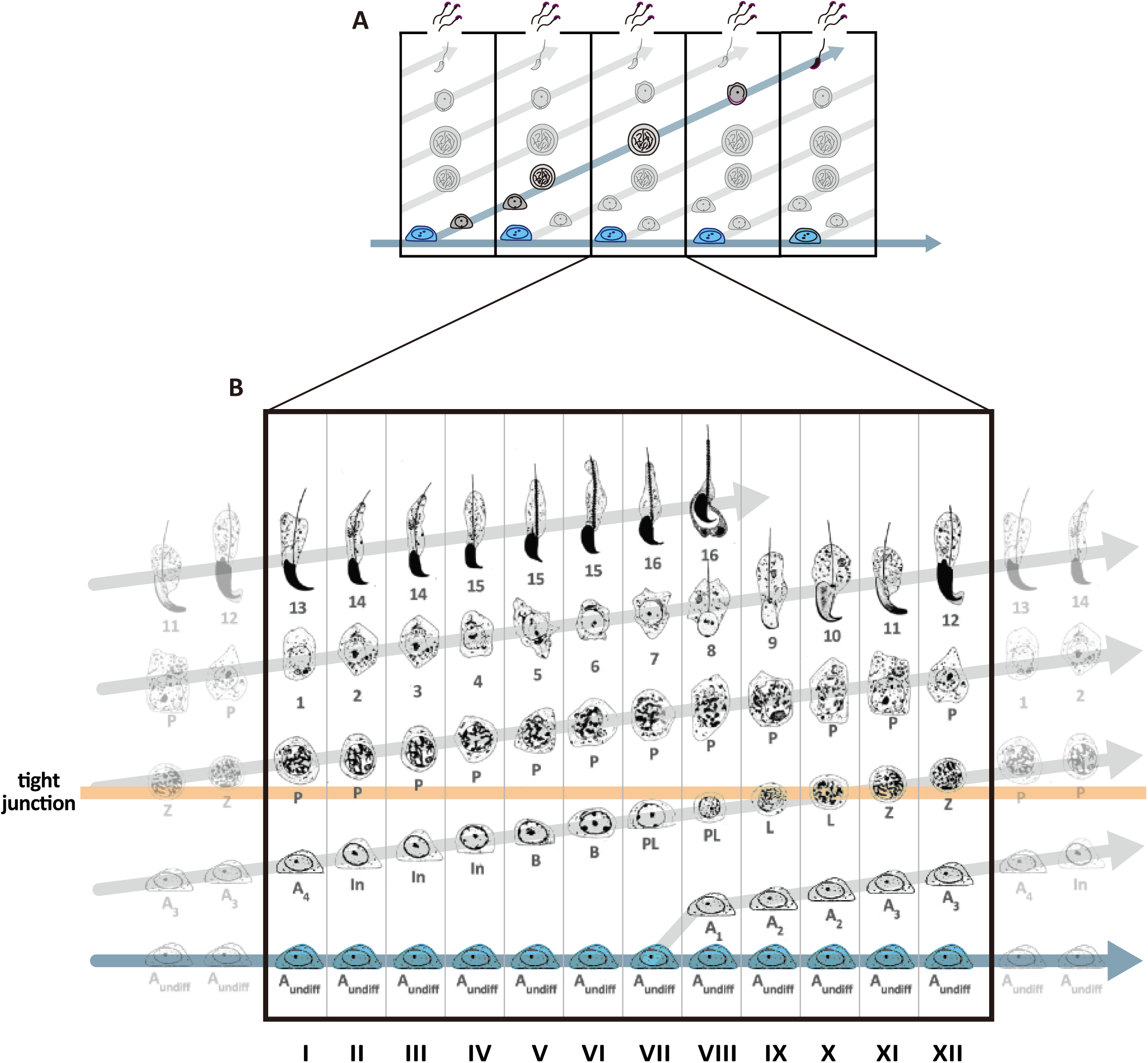
Spermatogenic cycle and germ cell turnover in seminiferous epithelium (related to Figure 1) **(A)** A scheme of periodic turnover of differentiating germ cells within the seminiferous epithelium, reproduced from Figure 1A. A_undiff_ (blue) periodically give rise to differentiating cells, so four generations of differentiating germ cells undergo tightly coordinated turnover to form the layered architecture of seminiferous epithelium. Each rectangle indicates one spermatogenic cycle, showing periodic changes in the associations of differentiating cells. **(B)** A detailed illustration of a spermatogenic cycle, i.e., one of the rectangles in (A). Due to the periodic and coordinated differentiation, a cycle is classified into 12 stages (I to XII) based on the associations of four germ cell generations. Note that A_undiff_ (bottom) give rise to the first committed spermatogonia (A_1_) in stage VII, which subsequently undergo differentiation through differentiating spermatogonia (A_1_, A_2_, A_3_, A_4_, In, B), spermatocytes (PL, preleptotene; L, leptotene; Z, zygotene; P, pachytene; D, diplotene), round spermatids (step 1 to 8), and elongating spermatids (step 9-16) which leave the tissue at stage VIII. During stages IX to XI, early spermatocytes (L and Z) translocate across tight junctions (schematically shown by an orange bar). As germ cells turn over by one generation, the stage returns to the initial one. Modified from Russell, 1990.^1^

**Figure S2.**
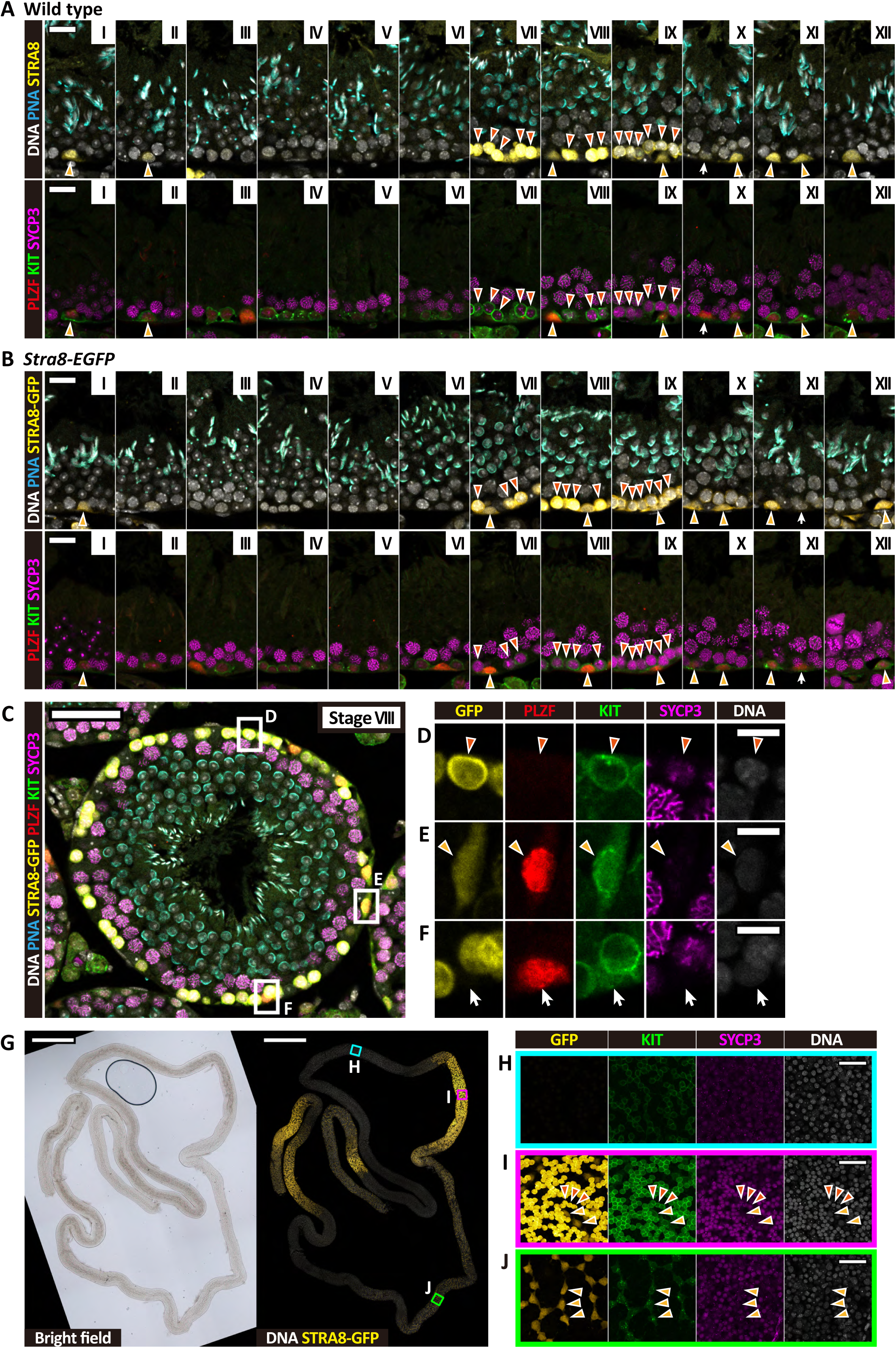
Stra8-GFP Expression during the spermatogenic cycle (related to Figure 2) **(A-B)** Cross-section images of multi-color-stained seminiferous epithelia of wild-type (A) and *Stra8-EGFP* (B) mice at indicated stages (I to XII). Stages and cell types were determined based on PNA-lectin (cyan)/DNA (gray) and PLZF (red)/KIT (green)/SYCP3 (magenta), respectively. Light and dark orange arrowheads indicate Stra8^+^ or Stra8-EGFP^+^ early differentiating spermatogonia and early spermatocytes, respectively. White arrows indicate undifferentiated spermatogonia (A_undiff_). The staining patterns used to determine these cell types are shown in (C-F). Scale bars, 20 µm in (A-B). Upper panels in (B) are identical to Figure 2B. **(C-F)** A cross-section of a *Stra8-EGFP* mouse seminiferous tubule stained as above (C), which includes different Stra8-GFP^+^ cell types as shown in higher magnification in (D-F). (D) An early spermatocyte which is Stra8-GFP^+^/PLZF^−^/KIT^+^/SYCP3^+^. (E) A differentiating spermatogonia which is Stra8-GFP^+^/PLZF^+^/KIT^+^/SYCP3^−^. Note that early differentiating spermatogonia retain PLZF protein.^85^ (F) An A_undiff_ which is Stra8-GFP^−^/PLZF^+^/KIT^−^/SYCP3^−^; the faint green nuclear signal is a leakage from the PLZF channel due to our 6-color staining methods, while the genuine surface KIT signal is absent. Scale bars, 50 µm in (C) and 10 µm in (D-F). **(G-J)** Whole-mount seminiferous tubule of a *Stra8-EGFP* mice, stained for Stra8-GFP, KIT, and SYCP3. (G) Low-magnification bright (left) and fluorescent (right) images, showing variable Stra8-GFP staining between different regions. (H-J) High-magnification images of the indicated areas in (G). (H) An area harboring no Stra8-GFP^+^ cells (cyan); (I) An area with Stra8-GFP^+^ spermatocytes and spermatogonia (magenta); (J) Stra8-GFP^+^ spermatogonia only (green). Light and dark orange arrowheads indicate Stra8-EGFP^+^ early differentiating spermatogonia and early spermatocytes, respectively. Scale bars indicate 1000 µm in (G) and 50 µm in (H-J). Some images are also shown in Figure 2C.

**Figure S3.**
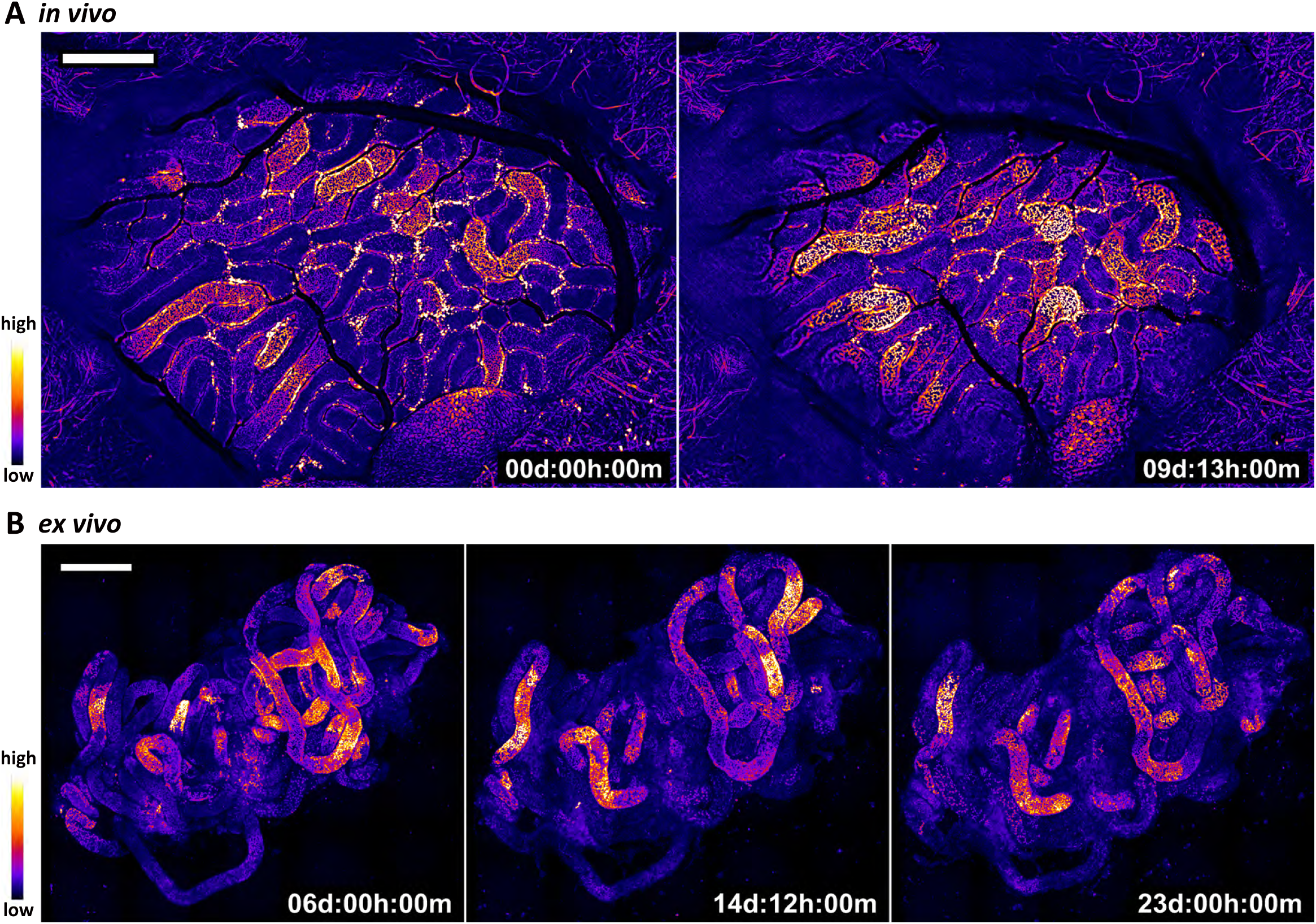
*in vivo* and *ex vivo* live imaging of *Stra8-EGFP* mouse testes (related to Figure 2, Video S1, and Video S3) Time-lapse images selected roughly once every spermatogenic cycle from *in vivo* **(A)** and *ex vivo* **(B)** filming of *Stra8-EGFP* mouse testes. The full image series are provided in Videos S1 and S3, respectively. GFP signal intensity is presented by pseudo color as indicated. Note that Stra8-EGFP signals came back in almost the same tubule segments; scale bars indicate 1000 µm in (A) and 500 µm in (B).

**Figure S4.**
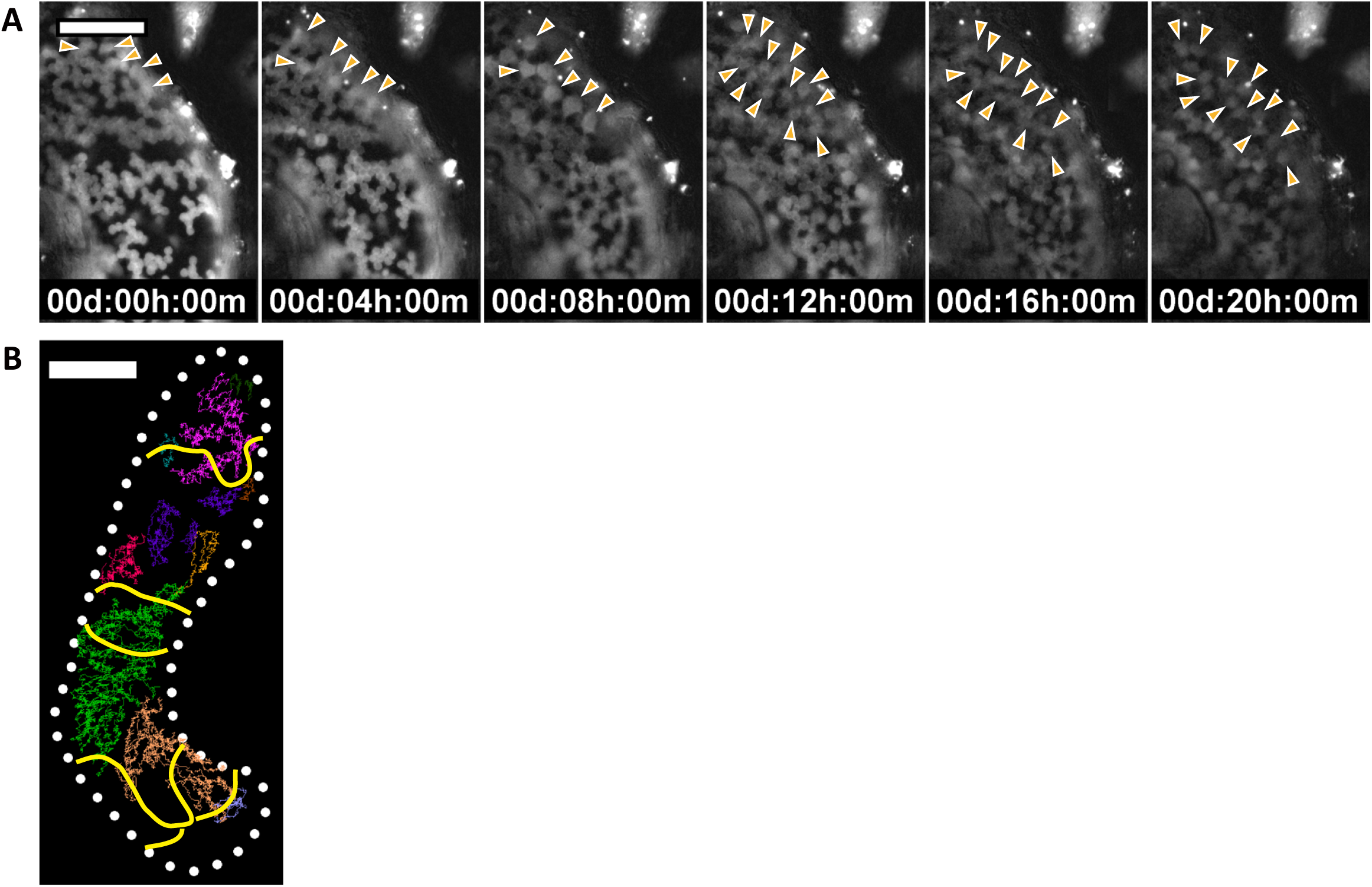
Supplementary data for *in vivo* live imaging of *Stra8-EGFP* mouse testes (related to Figure 3, Figure 4, Video S5, and Video S6) **(A)** Selected time-lapse images acquired by *in vivo* live imaging of a *Stra8-EGFP* mouse testis using a 30x objective (N.A. 1.05). The full image series is shown in Video S5. Light orange arrowheads indicate Stra8-GFP^+^ spermatogonia in a syncytium, collectively migrating from the vasculature-proximal region and synchronously dividing between 8h and 12h. Note that individual spermatocytes are visualized more clearly than in our routine, less harsh live-imaging setting using lower-power optics optimized for long-term continuous filming. Scale bar, 100 µm. **(B)** Overlay of the boundaries of synchronous clusters of Stra8-GFP^+^ spermatocytes (yellow lines; see Figure 4A) with the trajectories of Stra8-GFP^+^ spermatogonia belonging to multiple syncytia (indicated by the same colors as Figure 4C and Video S6). Note that the boundaries between territories of spermatogonial syncytia do not show apparent correlations with those between the synchronous clusters of spermatocytes. Scale bar, 200 µm.

**Figure S5.**
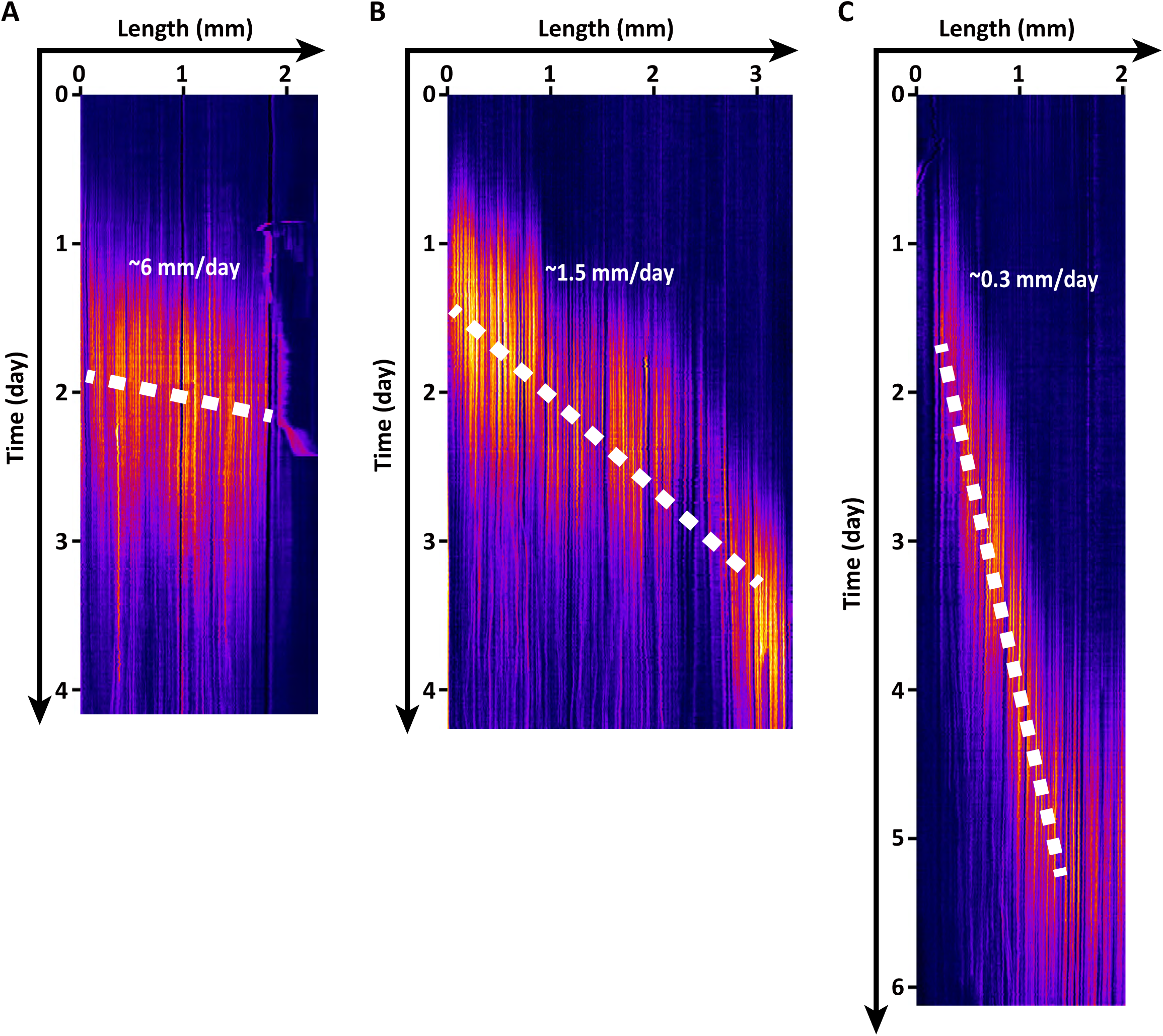
Velocity variance of the spermatogenic wave among seminiferous tubule regions (related to Figure 5) **(A-C)** Kymographs of the spermatogenic wave in three separate seminiferous tubule segments within a single testis, observed *in vivo*, based on the Stra8-GFP signal intensity. The averaged velocities along the dashed lines are roughly 6, 1.5, and 0.3 mm/day in (A), (B), and (C), respectively. (B) is reproduced from Figure 5B.

**Figure S6.**
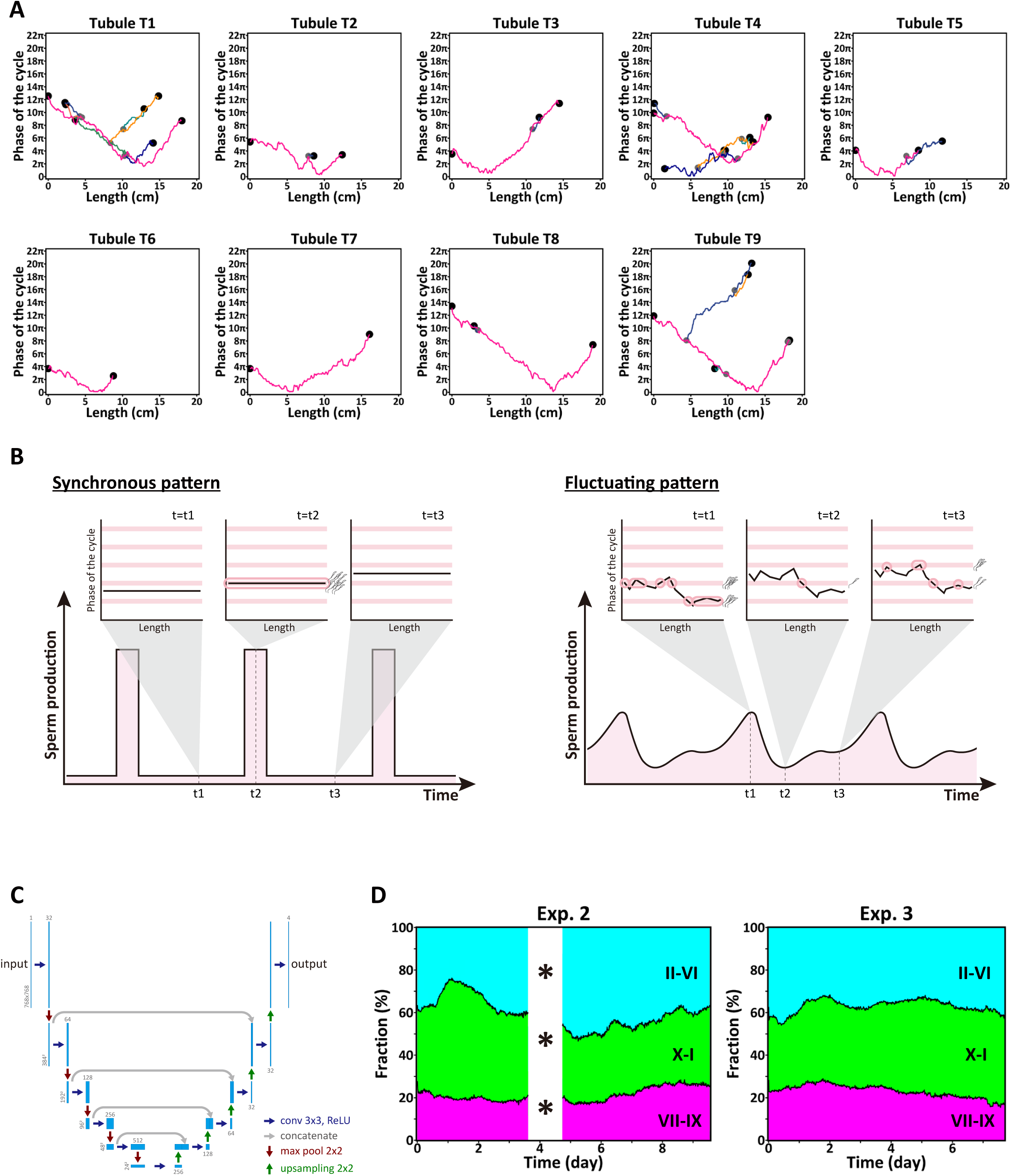
Organ-scale orders of the spermatogenic wave (related to Figure 6) **(A)** Phase distribution of the spermatogenic wave over the full lengths of seminiferous tubule loops, based on the precise stage mapping by 3D-reconstruction of serial sections of a testis, as described in Figure 6A and Method. Stage information (I to XII) was converted into phase value (0 to 2π) and plotted against the tubule length. Given the phase continuity between adjacent segments, local phases are shown by values ranging beyond 0-2π. All tubule loops included in a testis that had been reconstructed previously (tubule ID T1-T9; Nakata et al. 2017) were re-analyzed at a higher resolution and presented by phase plots. The boundaries of the tubule loops, i.e., junctions with the rete testis, are indicated with black dots. In branched loops (all but T6 and T7), branches are shown in different colors, with the gray dots marking the branching points. Note that a grand “jagged V-shape” pattern occurs frequently regardless of the branching pattern. An identical plot of tubule T7 is also presented in Figure 6C with a different color code. **(B)** Schemes explaining time variation of the summed sperm production in hypothetical seminiferous tubule loops, showing non-V-shaped phase profiles, namely, a perfect phase-locked synchrony (left) and highly fluctuating phase waves (right). Note that, compared with those showing a clear V-shape pattern (Figure 6D), the summed sperm production shows significantly larger temporal fluctuations. **(C)** A Scheme of the U-Net architecture used to categorize the tubule regions based on Stra8-GFP images in this study. **(D)** Kinetics of the proportion of the three phases of the spermatogenic cycle (GFP^+^ spermatocytes and spermatogonia shown in magenta; GFP^+^ spermatogonia only shown in green; and that without GFP^+^ cells shown in cyan), over the entire surface area of the testes live-imaged *in vivo*. The results of one of three experiments are shown in Figure 6E-F; that of the others are shown here. In Exp. 2, image data were not acquired fully due to a tentative technical problem (asterisks). Note the largely constant composition.

**Figure S7.**
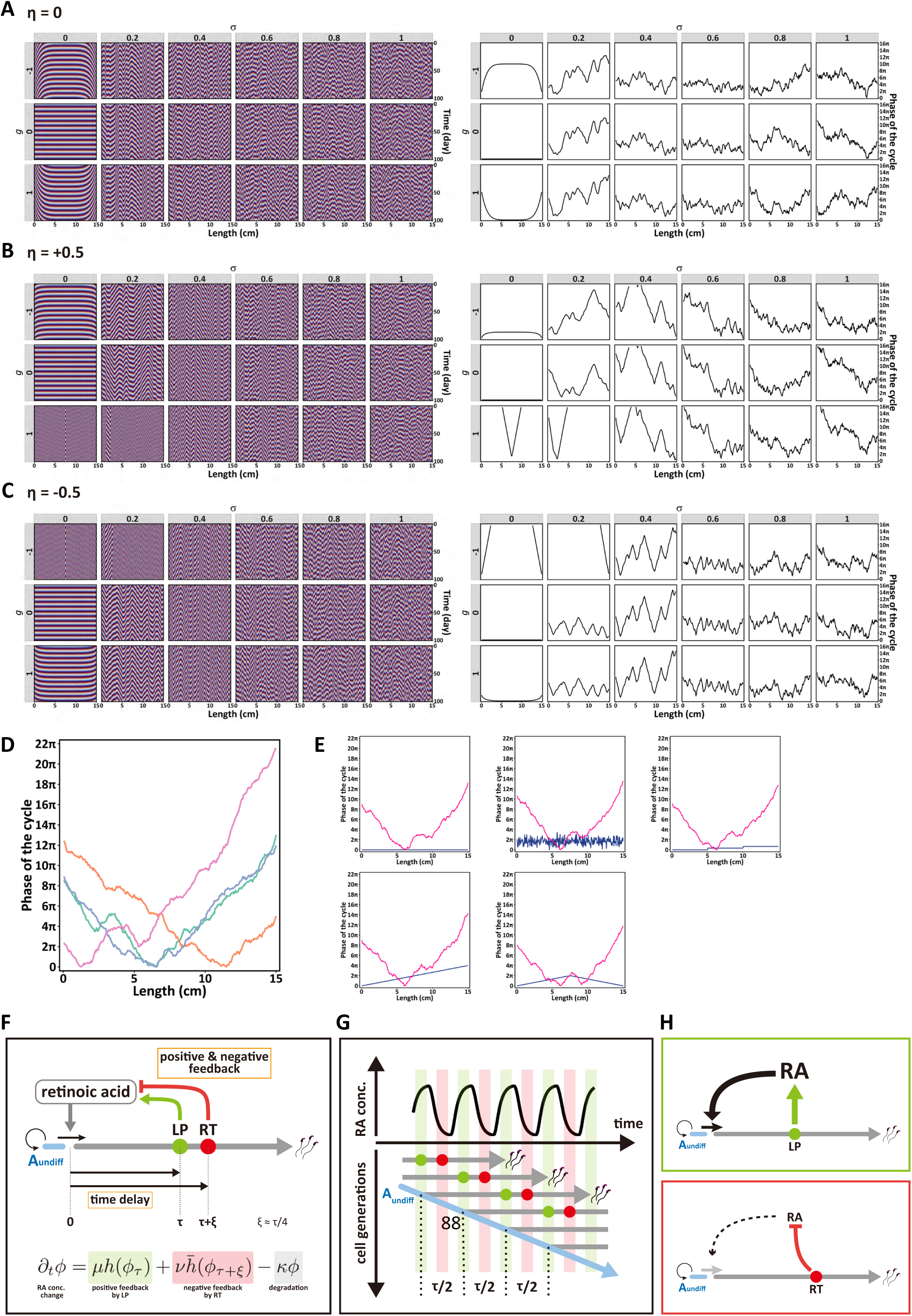
Deconstructing the dynamics of the spermatogenic wave as a model of locally coupled oscillators (related to Figure 7) **(A-C)** Parameter dependence of the classic Sakaguchi-Kuramoto model of the spermatogenic wave based on a one-dimensional array of coupled oscillators. Kymographs (left) and phase maps in long-term steady state (right) are shown when the asymmetry parameter *η* is set as 0 (A), +0.5 (B), and −0.5 (C) while *σ* and *g* are systematically changed (For details of the model and parameter notation, we refer to the main text and Supplementary Theory). **(D)** Representative phase maps of multiple simulations of stochastic Sakaguchi-Kuramoto mode, in which different colors indicate simulations from independent quenched spatial noises, demonstrating the robustness of the jagged V-shape pattern. **(E)** Representative phase maps (magenta lines) of stochastic Sakaguchi-Kuramoto model starting from various initial phase distributions (blue lines; uniform, random, stem-wise, slope, and inverted V-shape). Note that the Jagged V-shape is robustly generated independent of the initial condition albeit not perfectly identical. **(F-H)** Synthesis of RA pathway-based coupled oscillator model and emergence of phase-wave dynamics (see Supplemental Theory for details). (F) A detailed scheme of time-delayed feedback regulation of the RA model shown in Figure 7G. (G) Persistent oscillations of RA concentration occur in concert with the A_undiff_-to-A_1_ commitment and subsequent differentiation of germ cells. Note that the interval is equal to half the duration from SSC commitment to becoming LPs (tau). (H) Explanatory schematics of the state of RA metabolism and its impact on germ cell differentiation, in opposing phases of the oscillation colored in green (top) and red (bottom) in (G).

## Supplemental videos

**Video S1:** A compiled movie of an *in vivo* live-imaged adult *Stra8-EGFP* mouse testis, presenting the entire observed area of the testis surface. Scale bar, 1000 µm. Related to Figure 2E and S3A.

**Video S2:** An *in vivo*-imaged movie showing a small region of a *Stra8-EGFP* mouse testis, covering the entire spermatogenic cycle. Scale bar, 100 µm. Related to Figure 2G.

**Video S3:** A compiled movie of an *ex vivo* live-imaged *Stra8-EGFP* mouse seminiferous tubules shown in Figure 2I. Inset shows the magnified image in large yellow rectangle shown in the inset of Figure 2I; small yellow rectangle indicates the area shown in Figure 2J and used for GFP signal intensity measurement in Figure 2K. Scale bar, 500 µm.

**Video S4:** An *in vivo*-imaged movie showing a small region of a testis, highlighting the spatiotemporal coordination between Stra8-GFP^+^ spermatocytes and spermatogonia. Scale bar, 100 µm. Related to Figure 3A.

**Video S5:** A high-resolution *in vivo*-imaged movie showing a small region of a testis, acquired using a 30x objective lens. Scale bar, 100 µm. Related to Figure S4A.

**Video S6:** An *in vivo*-imaged movie covering a stretch of a seminiferous tubule (left) and traces of Stra8-GFP^+^ spermatogonia, in which spermatogonia belonging to the same syncytium are shown in the same color (right). Scale bar, 200 µm. Related to Figure 4 and S4B.

**Video S7:** An *in vivo*-imaged movie covering an extended segment of a seminiferous tubule (left), and computationally linearized images of the outlined region (right). Scale bar, 200 µm. Related to Figure 5A-B and S5B.

**Video S8:** An *ex vivo*-imaged movie covering a segment of a seminiferous tubule (left, indicated by a yellowish-gray line) and computationally linearized images of the indicated segment (right). Note the three rounds of largely regular traveling waves. Scale bar, 200 µm. Related to Figure 5C-F.

**Video S9:** An *ex vivo*-imaged movie covering an extended segment of a seminiferous tubule (left, indicated by a yellowish-gray line) and computationally linearized images of the indicated segment (right). Note the propagation of three rounds of traveling waves showing significant irregularities explained in Figure 5G-I and the text. Scale bar, 200 µm.

**Video S10:** 3D-reconstructed seminiferous tubules from serial testis sections, highlighting T7 tubules with the phase map. Related to Figure 6B.

**Video S11:** A compiled movie of an *in vivo* live-imaged adult *Stra8-EGFP* mouse testis (left) and segmentation into three phases (magenta, green and cyan) using machine learning (right). Scale bar, 1000 µm. Related to Figure 6E-F.

## Supplementary Theory

Here, we develop in further detail aspects of the modeling-based analysis of the spatiotemporal organization of spermatogenic differentiation within testicular seminiferous tubules presented in the main text. Specifically, in the first section, we start with the classic Sakaguchi-Kuramoto model and explore various extensions to the model in search of a minimal framework to explore the dynamics of locally coupled phase oscillators. We demonstrate that, under general conditions, the stochastic extension of the classic Sakaguchi-Kuramoto model can explain the qualitative properties of the observed spermatogenic wave as a self-organization phenomenon. Based on this model, we then use the results of stochastic simulation to infer the best-fit parameters of the model from experimental data obtained from measurements of the stage dependence of fixed tissue. The best-fit parameters provided the model with a high predictive capacity for the observed intricate phase-wave behavior of mouse spermatogenesis involving a range of irregularities and global phase patterning over the loops of seminiferous tubules (descent of segmental order). Next, we present a refined model tailored to predict the cellular and molecular mechanism of the local spermatogenic cycle, potentially constituting the elemental components of the overarching macroscopic stochastic Sakaguchi-Kuramoto model. This microscopic model incorporates delayed feedback mechanisms between the retinoic acid (RA) metabolism in differentiating germ cells and the commitment of spermatogenic stem cells for differentiation promoted by RA. Finally, by extending this microscopic model of the spermatogenic cycle to the one-dimensional geometry of the seminiferous tubules, we show that the dynamics of this more complex system mirror the behavior of the minimal stochastic Sakaguchi-Kuramoto model.

### T1. SAKAGUCHI-KURAMOTO-TYPE MODELS OF THE SPERMATOGENIC WAVE WITHIN SEMINIFEROUS TUBULES

To begin, we first consider the collective dynamics of the local spermatogenic cycles at the length scale of the seminiferous tubule. Measurements based on intravital imaging of Stra8-GFP expression show that the progression of the spermatogenic wave along the tubule length proceeds in a step-like manner, with local clusters of germ cells becoming synchronized in their phase of expression with small but definite phase shifts between neighbors (see main text and Figure 4), suggesting the existence of local interactions of the phase of the spermatogenic cycles between neighboring sites (locally synchronized cell clusters). While there are a number of candidate cellular or molecular mechanisms that could mediate local coupling of the phases of clusters, such as RA diffusion along the length of the tubule or cell-cell contact between germ cell syncytia and/or supporting somatic cells called Sertoli cells, we place emphasis on the experimental phenomenology, setting aside the question of the precise underlying molecular mechanisms that support such interactions. Instead, we take a minimalist approach by taking the local spermatogenic cycles as given and explore whether local coupling alone is sufficient to give rise to the experimentally observed phenomena, including the spontaneous appearance of phase waves, stretches of reversed velocity (modulations), and the characteristic phase distribution over the seminiferous tubule loops in the form of jagged V-shape pattern, as described in the main text. To this end, we first introduce the classic Sakaguchi-Kuramoto coupled oscillator model as the simplest theory to explore the dynamics of the spermatogenic wave. While this simple theory qualitatively captures the basic phenomena observed in experiments, the parameter regime that gives rise to consistent global V-shapes is not compatible with the number of local variations observed along the tubules. To improve upon the minimal model, we explore various extensions, including quasi-1D, next-nearest-neighbor, and stochastic Sakaguchi-Kuramoto models, and find that the stochastic Sakaguchi-Kuramoto model provides the best quantitative match with experimental data, as detailed below.

To begin, we first questioned whether the range of experimental behavior could be captured by a minimal phenomenological description. Noting that the phase varies little in the direction perpendicular to the axis of the tubule (Figure 4), we began by modeling individual seminiferous tubules as a one-dimensional array of discrete lattice sites, 1 ≤ *i* ≤ *N*. At each lattice site, the phase of the spermatogenic cycle is indexed by the angular coordinate *θ_i_*(*t*), which varies periodically over time on the internal [0, 2*π*] and is driven at a frequency *ω_i_*. To accommodate potential spatial heterogeneities along the length of the tubule, we proposed that the frequencies are defined by a constant average *ω̅* with static fluctuations, Λ*_i_*drew, for simplicity, from a normal (Gaussian) distribution with zero mean and variance ⟨Λ*_i_*Λ*_j_*⟩ = *σ*^2^*δ_ij_*. Later, we will consider the impact of dynamic fluctuations in the local frequencies. We further hypothesized the existence of a weak local nonlinear coupling of the phases as the simplest non-reciprocal interaction between neighboring models, which we represent in the minimal form of the classic Sakaguchi-Kuramoto model [1],

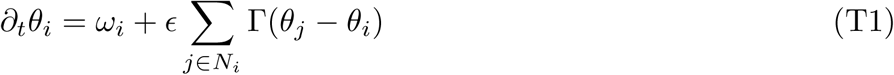

**FIG. T1.**
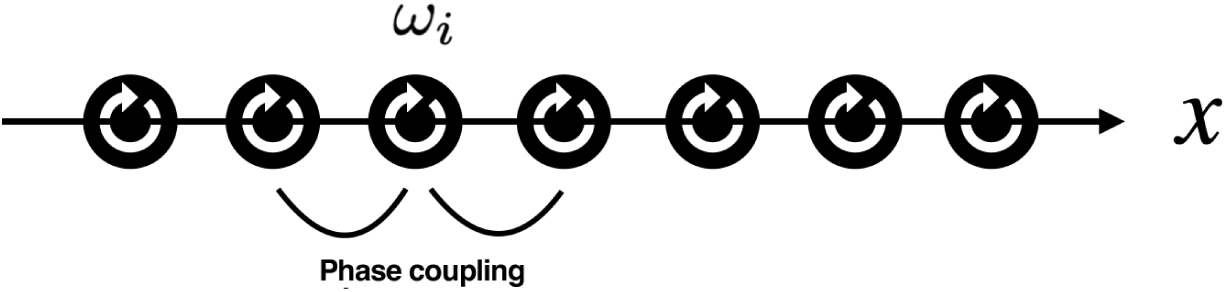
Schematic illustration of the Sakaguchi-Kuramoto type models used to describe the spermatogenic wave. Here, seminiferous tubules are modelled as a one-dimensional array of cell clusters in which the phase of the oscillation, the spermatogenic cycle, is synchronized. At each cluster, or lattice site *i*, the phase *θ_i_* of the oscillation is driven at frequency *ω_i_*, with phases coupled weakly between neighboring lattice sites along the axis *x* of the tubule.

where, for a given lattice site *i*, the sum runs over all nearest neighbors *N_i_*. Here, Γ(*θ*) = sin *θ* + *η*(1 − cos *θ*) denotes the nature of the coupling with *ɛ* its strength. When *η* = 0, the coupling is reciprocal, so that neighbors exert a reciprocal (equal and opposite) force on each other. However, when *η* ≠ 0, it follows that Γ(*ϕ*) ≠ Γ(*ϕ*), implying that forces between a nearest-neighbor pair with phase difference *ϕ* are not reciprocal, driving the system out of equilibrium [2]. Indeed, there are a number of candidate biological mechanisms that can drive non-reciprocity, such as check-pointing in cell cycles and saturation of response (see section T3). However, for now, we remain agnostic to the precise mechanisms that underpin the form of the phase interaction and focus instead on the macroscopic behavior of the model system.

To explore the dynamics of the system, we first noted that, without loss of generality, we may set the parameter *ɛ* to unity by rescaling the time coordinate and frequency scale, leaving three adjustable parameters: *η*, *ω̅*, and *σ*. Since both ends of individual tubules are connected to the rete testis, and may experience a singular microenvironment that slows down or speeds up the cycles, we adopted fixed gradient boundary conditions, setting *θ*_1_ − *θ*_0_ = *g*_1_ and *θ_N_*_+1_ − *θ_N_* = *g*_2_, where *g*_1_ and *g*_2_ are constants. Based on the observed antisymmetry of the two ends of a tubule (manifest in the V-shaped pattern of measured phase velocities, as well as the symmetric anatomy of the tissue), we set *g*_2_ = − *g*_1_ = *g*. The fixed gradient boundary condition allows us to perform a Galilean transformation 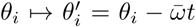 that enables the average frequency *ω̅* to be gauged away, i.e., without loss of generality, we may set *ω̅* = 0 with *θ_i_* now representing spatial and temporal fluctuations of the phase on top of an overall constant cycle at frequency *ω̅*. We will see below that, for a large system size *N* and non-vanishing disorder strength *σ*, the boundary conditions do not play a key role in determining the overall *θ*-profile.

For certain parameter ranges (see below), the oscillators eventually become entrained into a synchronized state, where the phases at all sites oscillate at the same frequency Ω with a constant spatial phase profile,

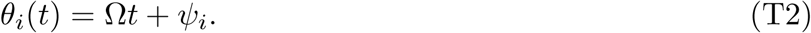

Moreover, it has been shown that in addition to reducing the strength of the disorder *σ*, increasing the asymmetry parameter *η* also facilitates synchronization across large length scales [2, 3]. (Technically, this requires the distribution of *ω_i_*to be bounded from above by some high frequency; but in realistic scenarios, the frequency of oscillators must remain finite.) In cases where the dynamics do not reach system-wide synchronization, the steady state comprises locally synchronized patches that decrease in length as the strength of disorder *σ* is increased [2, 4].

#### A. The general profile of the phase wave, *θ_i_*

To gain insight into the general behaviors of the system, we first assumed that the system is positioned in a parameter regime where the phases become fully synchronized. Substituting Eq. (T2) into (T1), and defining *ϕ_i_* = *ψ_i_*_+1_ − *ψ_i_*, we obtain

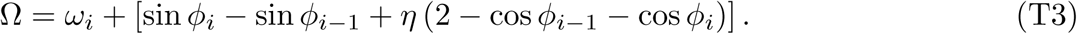

Summing over sites *i* and dividing by *N*, we obtain the constraint

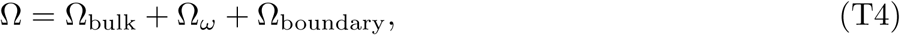

where

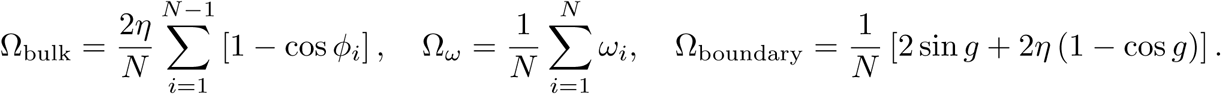

Here, Ω_bulk_ ~ *O*(1) represents the bulk contribution to the overall frequency, 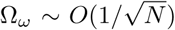 arises from the (Gaussian) disorder in the drive frequencies, and Ω_boundary_ ~ *O*(1*/N*) stems from the fixed-gradient boundary conditions. If we focus on the large-*N* system, the leading contribution arises from the bulk term, meaning that Ω has the same sign as the asymmetry parameter *η*.

To understand qualitatively the form of the steady-state phase profile *θ_i_*, we then considered the continuum limit, keeping terms that are lowest order in the lattice spacing (i.e., the lowest gradients of the phase field *ϕ*). Defining the continuum fields *ψ_i_ → ψ*(*x*), *ϕ*(*x*) = *∂_x_ → ψ*(*x*) and *ω_i_ → ω*(*x*), and rearranging the terms, the stationary part of the classic Sakaguchi-Kuramoto model (T1) takes the form

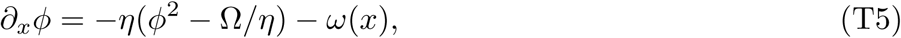

where the lattice spacing is rescaled away. Here, *ω*(*x*) represents a *δ*-correlated white-noise field with ⟨*ω*(*x*)*ω*(*y*)⟩ = *σ*^2^*δ*(*x* − *y*), where ⟨· · · ⟩ denotes the quenched disorder average.

Note that, if we read the spatial coordinate *x* as an effective “time” variable, Eq. (T5) is mathematically equivalent to the equation of motion of an overdamped particle moving in a potential *U* (*ϕ*) connected to a thermal bath of temperature *σ/*2, where *∂_ϕ_U* (*ϕ*) = *η*(*ϕ*^2^ – Ω*/η*). The shape of the potential for negative and positive *η* are depicted in Figure T2. Although the potential is unbounded from below, the constraints given by fixed gradient boundary conditions dictate that the particle cannot descend down the slope indefinitely.

Based on the stage reconstruction of the spermatogenic wave, there is evidence from experiments that the spatial profile of *θ* is generally characterized by a downward “V-shaped” dependence as observed experimentally (see Figure 6, main text, and below). In the absence of disorder (*σ* = 0), a prominent V-shape profile calls for a positive value of the asymmetry parameter *η* and *g >* 0. Self-consistently, we must have Ω*/η ≈ g*^2^, placing the two stationary points of *U* (*ϕ*) at ± *g*. The V-shape profile then corresponds to the “particle” *ϕ* staying at the peak at − *g* before sliding down the slope to arrive at +*g* on the other side. Any asymmetry in the boundary conditions (i.e., *g*_1_ ≠ *g*_2_) leads to one of them not translating to a stationary point, and hence a strongly lopsided V-shape.

However, this picture changes dramatically in the context of non-vanishing spatial disorder *ω*(*x*), as shown in Figure T3. With the possibility of noise-driven transitions between the peaks, the V-shape pattern can also occur for negative *η*, as there can be an “instanton-like” path from the trough towards the peak on the right, facilitated by appropriate noise configurations. The disorder is also responsible for the generation of smaller-scale chance reversals in the direction of the phase wave (defined in literature as modulations as described in the main text), superposed on the back of the overall V-shape (see Figures 6C, S6A, and T5 below). This occurs when there are multiple noise-driven transitions, or when the transition does not fully reach the other stationary point, as shown in the last three panels of the third rows of Figure T3. (For a more comprehensive discussion of the long-term stability of the model as a function of the parameters, we refer to Ref. [4].)

**FIG. T2.**
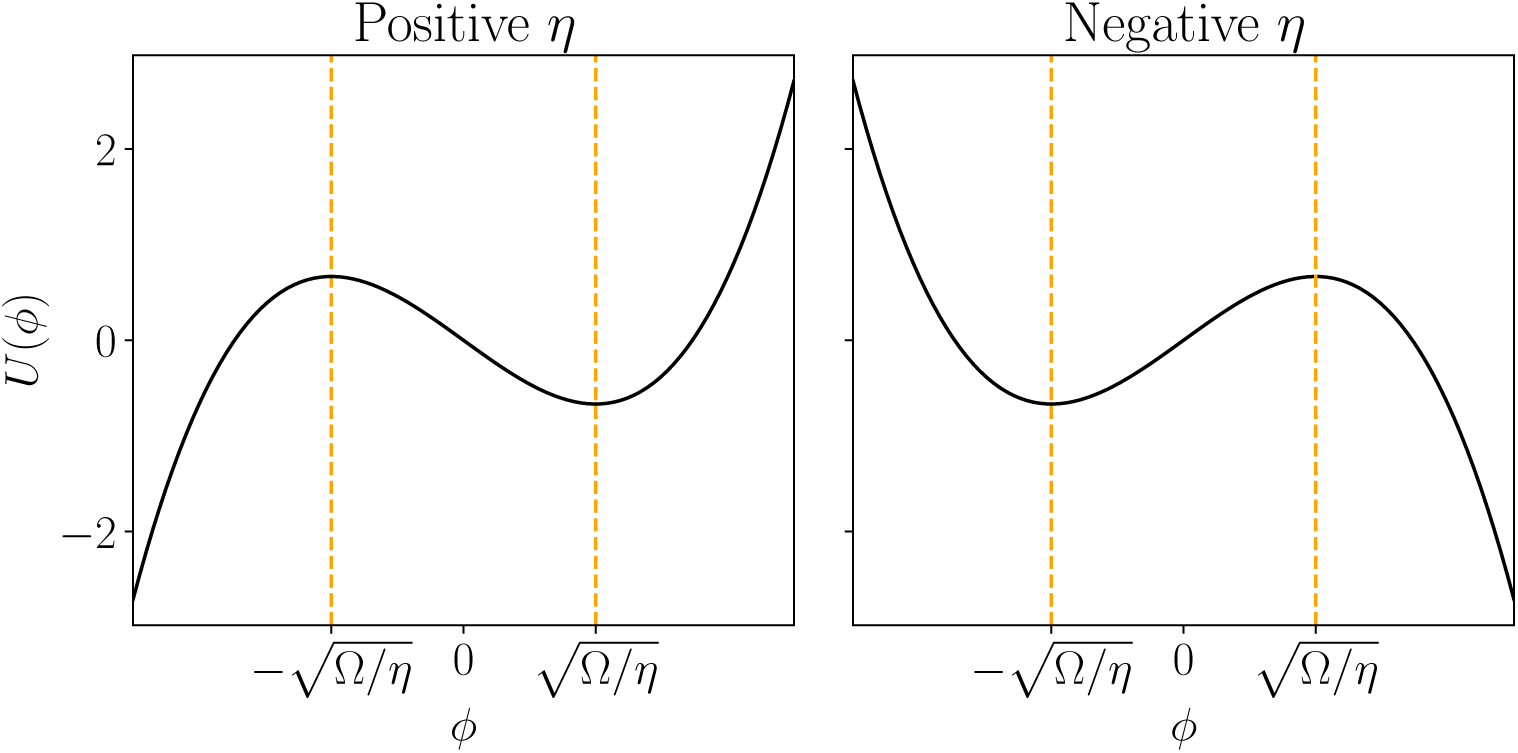
Effective potential for the phase for the coupled oscillator model. At steady-state, the spatial profile of the phase wave acquires a stationary solution defined by Eq. (T5). Here, we show the shape of the effective potential *U* (*ϕ*) for positive and negative *η*. For positive *η*, the local minimum lies at positive *ϕ* while the local maximum lies at negative *ϕ*. The positions of the stationary points are the opposite for negative *η*.

Based on these findings, we conclude that the classic Sakaguchi-Kuramoto model can qualitatively reproduce phase waves that manifest the characteristic irregularities (Figure 5) and the global V-shape pattern over the tubule loops (Figure 6) observed in experiments. To gain deeper insight into the suitability of the model, we then turned to a more quantitative evaluation of the validity of the model using the Maximum Likelihood Estimation.

#### B. Maximum Likelihood Estimation for parameter inference

In general, the inference of parameter values from non-equilibrium stochastic models is highly non-trivial since the probability distribution of the steady-state solution is usually not known. In the case of Sakaguchi-Kuramoto-type models, this is the situation that prevails. However, in the case where the steady-state solution is phase synchronized across the length of the system (i.e., *∂_t_ϕ_i_*= 0 for all sites *i*), it is possible to write down the probability distribution of *ϕ_i_ exactly* using Eq. (T3). Rearranging Eqs. (T3) and (T4), we have for each site *i*

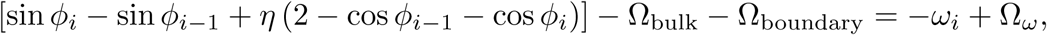

where we have moved Ω*_ω_* to the right-hand side as it is a sum of *ω_i_*. Then, summing over the first *n* sites, we have

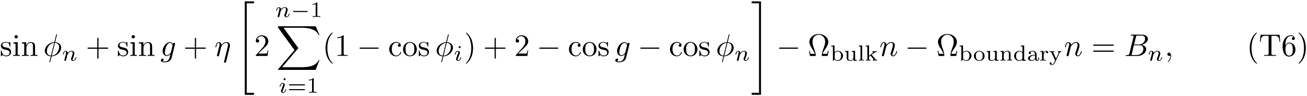

where *B_n_* is a “Brownian Bridge” type process [5] defined as *B_n_* = *W_n_*−*nW_N_ /N* with 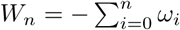 representing a discrete Wiener process. The Brownian Bridge is a multivariate Gaussian process with covariance *C_nm_*= *B_n_, B_m_* = *σ*^2^*n*(*N m*) (for *m > n*) and zero mean [5]. This allows us to write down the probability of a realization of a Brownian Bridge exactly. Denoting the Brownian Bridge process as vector **B** and the covariance as matrix **C**,

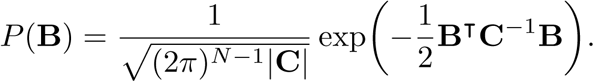

**FIG. T3.**
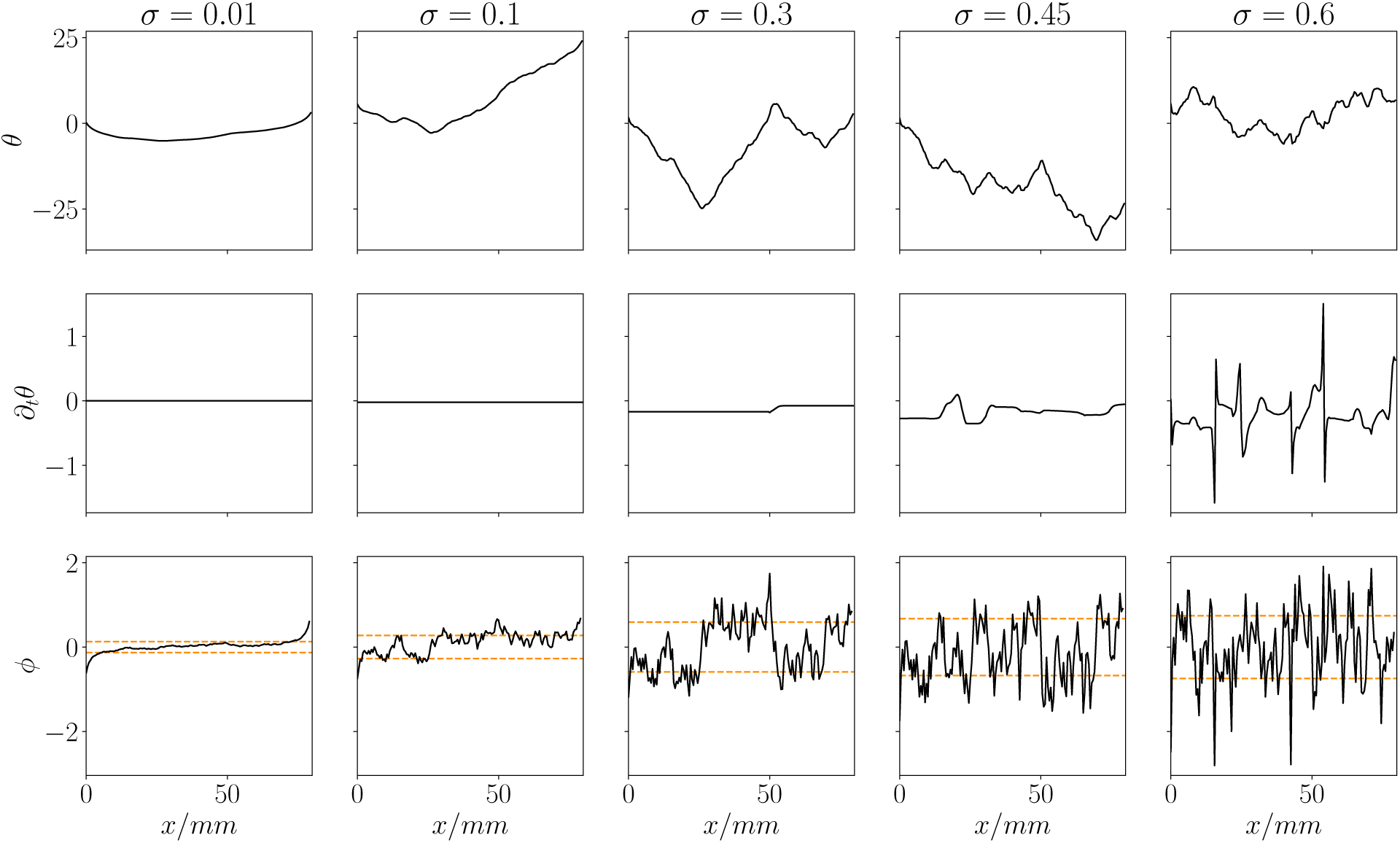
Solutions of the phase field *θ*(*x*), *∂_t_θ*, and *ϕ*(*x*) = *∂_x_θ* from the numerical integration of Eq. (T5) at long times for increasing values of the disorder strength, *σ*. The orange dashed line in the final row corresponds to the stationary points of the effective potential *U* (*ϕ*), calculated retrospectively from the *ϕ* profile. The model is solved with *η* = −0.44 and the fixed gradient boundary condition with *g*_1_ = − 1.3. The *ω_i_* distribution is the same for all simulations.

The probability of observing a realization of *ϕ* is the same as the probability of the corresponding Brownian Bridge from Eq. (T6) multiplied by the Jacobian determinant,

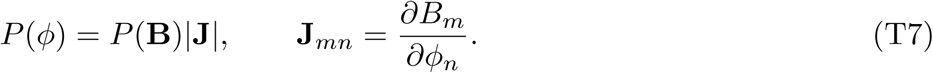

These calculations give the probability of observing a data set given the parameters. In other words, one can calculate the likelihood exactly (so long as we assume the system is fully synchronized). Hence, we can adapt the Maximum Likelihood Estimation (MLE) to infer the parameters. Overall, assuming a flat prior for all parameters, the fitting procedure is as follows:

1. For a set of parameters *η* and *g*, compute the corresponding Brownian Bridge **B** of *ϕ_i_* using Eq. (T6) and the Jacobian determinant.
2. Obtain the MLE for *σ* as 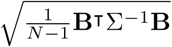, where Σ*_nm_* = *n*(*N* − *m*) (equivalent to the correlation matrix with *σ* = 1).
3. Calculate the log-likelihood of {*ϕ_i_*} using Eq. (T7).
4. Minimize the log posterior (which is the same as the log-likelihood in the current situation since we assume a flat prior) over values of *η* and *g* to obtain the Maximum Likelihood estimate for *η, g*.

**FIG. T4.**
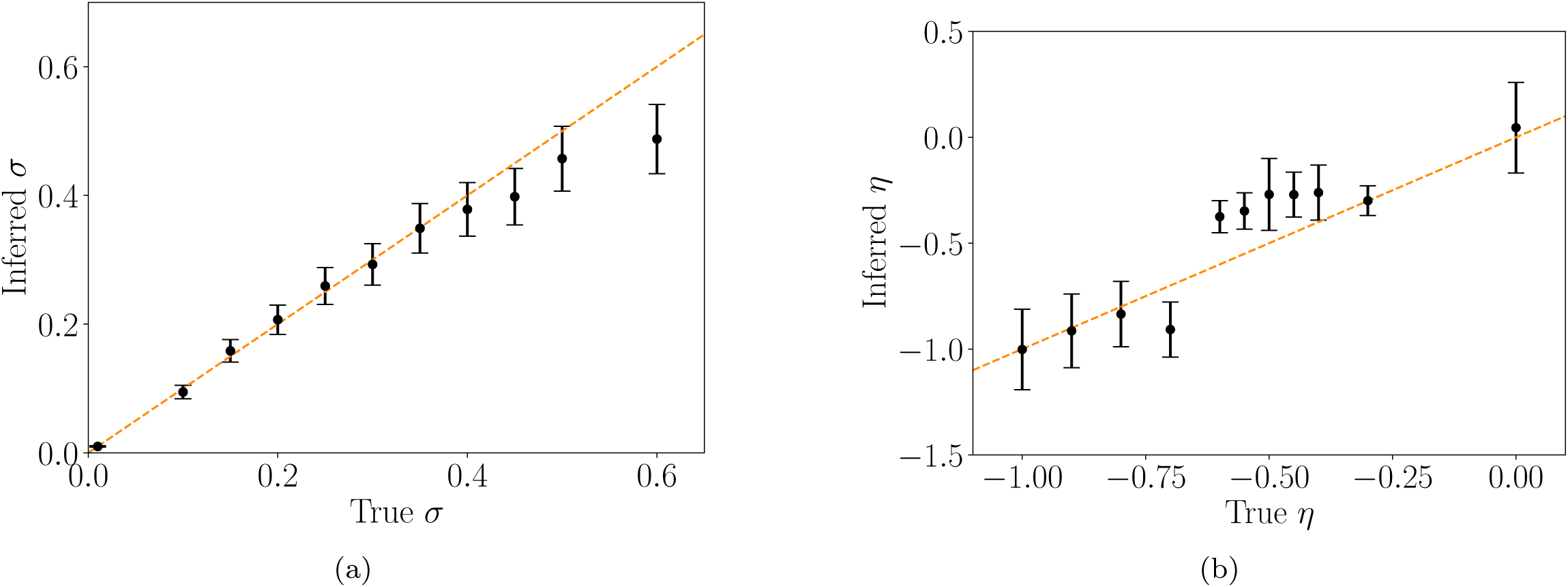
Comparison between the true parameters of the classic Sakaguchi-Kuramoto model and those inferred from a Maximum Likelihood(ML) Estimate based on a stochastic simulation of the model. The orange dashed lines in both panels represent perfect inference, and the black dots represent the ML estimates with the associated error bars using the ML method. (a) Inference of *σ* for *η* = −0.45 and *g* = 1. (b) Inference of *η* for *σ* = 0.44 and *g* = 1.

Once ML estimates are obtained, we can numerically calculate the error bounds by computing ratios of likelihoods at ML values and deviations from the ML, and then select the cutoff as the commonly used value of 1.92 to obtain the 95% confidence intervals.

To gain confidence in the method, we first assessed the ML method using simulation data by comparing the true parameter values against inferred ones for *η* and *σ* in Figure T4. (Note that the *g* comparison is not shown as the estimated error bar of *g* is typically large – note that varying *g* does not significantly affect the outcome. See also Figure T6 below.) As shown in the previous numerical simulations of the model (see Figure T3), for *η* = −0.45 (used in panel (a) of Figure T4), the system stops becoming fully synchronized at a disorder strength of around *σ* ≥ 0.3. Further numerical investigation also showed that, for a fixed *σ* of 0.44 (used in panel (b) of Figure T4), the system de-synchronizes at an asymmetry parameter of around *η* ≥ −0.8.

As expected, when applied to the computationally generated mock data sets, the numerical algorithm for the fitting procedure performs extremely well in the region of parameter space where the system becomes fully synchronized. Here, the ML procedure gives accurate parameter estimates, as demonstrated towards the left-hand side of both panels of Figure T4. Further, the ML algorithm is still able to estimate *σ* well beyond these limits: the right portions of both panels show reasonable agreement between the inferred values and true values, taking into account error bars. On the other hand, the stochastic simulations show that, when the system does not reach phase synchronization across the tubule, the algorithm tends to underestimate the magnitude of *η* and *σ*, a feature that should be kept in mind when we infer parameters from the experimental data.

Finally, we emphasize that only two assumptions have been made in the inference scheme, while the rest of the analysis is formally exact:

1. The system is fully phase synchronized at steady-state.
2. The distribution of the drive frequencies *ω_i_* is well described by a normal distribution.

**FIG. T5.**
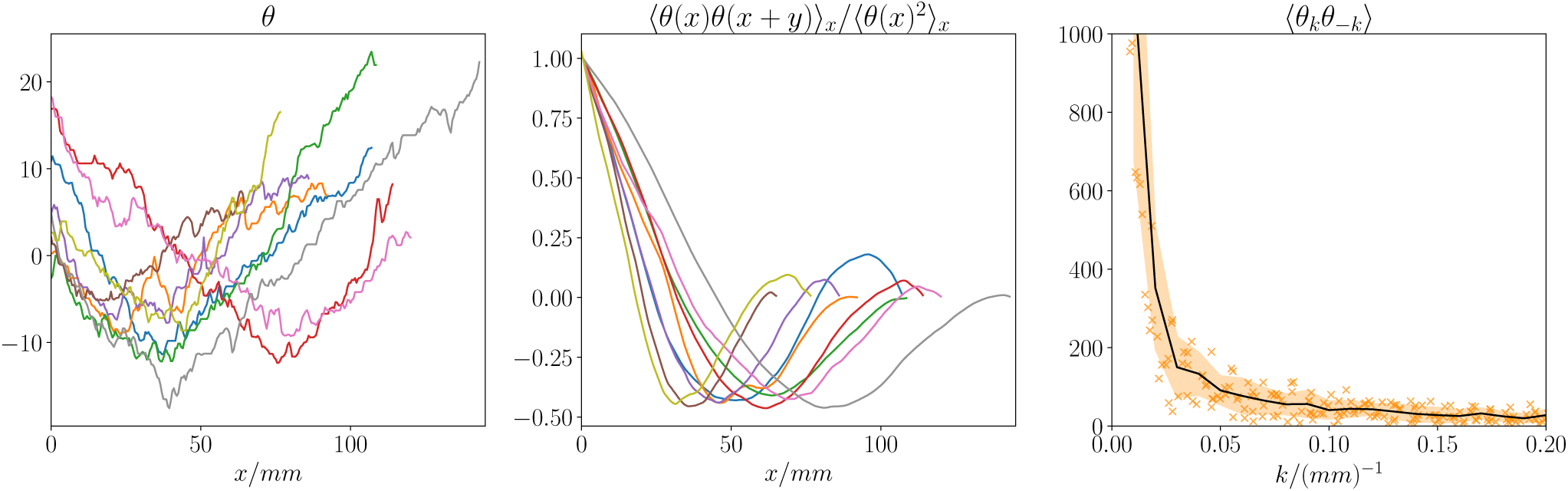
Histological stage measurements of 9 tubules (left) along with the autocorrelation (middle) and power spectrum (right) of *θ*. The corresponding phase values *θ_i_* were first shifted (by ±2*π*) to be continuous and then smoothed by averaging over every 50 measurements, with a final lattice spacing of approximately 500 *µm*. In the right panel, data from individual tubules are represented by orange crosses. The black line is the average and the orange area marks the standard errors. Experimental measures were obtained by revising the published data in Ref. [6], to gain a higher-resolution dataset (see STAR Method for detailed procedure).

**FIG. T6.**
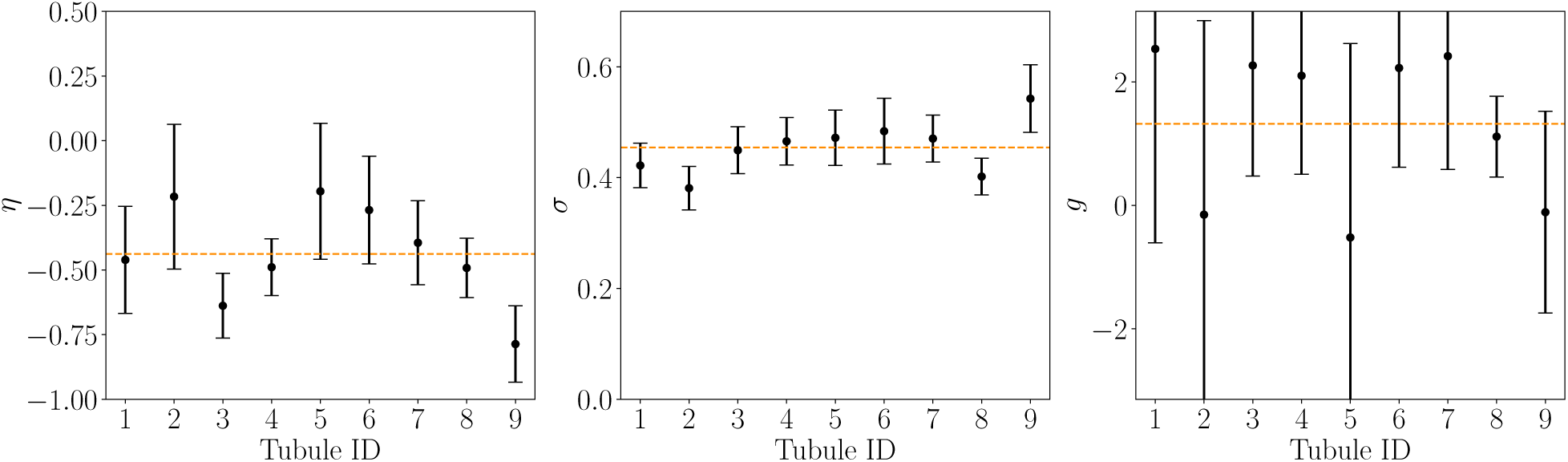
Parameter estimation of the asymmetry parameter *η*, the disorder strength *σ*, and boundary gradient *g* from measurements taken from 9 tubules (see Figure T5). For details of the fit procedure, see main text of Supplementary Theory. The orange dashed lines indicate the mean of the inferred values from all tubules. The error bars indicate 95% confidence intervals.

#### C. Assessing the quality of fit of the classic SK model

With these preliminaries, we then turned to infer the values of *η*, *σ*, and *g*, as well as the associated error bars, from the experimental data using the ML method outlined above. Here, to assign the local phase of the spermatogenic cycle, we made use of morphological classifications of stages from a total of 9 tubules from Ref. [6]. The histological identification classifies the cycle into 12 stages, with durations of 22.0, 18.1, 8.7, 18.6, 11.3, 18.1, 20.6, 20.8, 15.2, 11.3, 21.4, and 20.8 hrs, respectively [7]. After converting the discrete stage number to the midpoint of the stage times, we then divided the time by the sum of stage times to arrive at a phase variable within the range [0, 2*π*]. This procedure, compounded with measurement errors of (±1 stage), gives an error of approximately ±*π/*6 for each data point. The phases of each tubule were then averaged over every 50 measurements to reduce the measurement error to ±0.1. The smoothed data used in the subsequent inference scheme are shown in Figures 6C, S6A, and T5.

The ML estimates and errors for *η*, *σ*, and *g* are plotted in Figure T6. The error bars (95% confidence intervals) overlap significantly for all three parameters, implying that the best-fit Sakaguchi-Kuramoto models for all tubules are the same within tolerance. We note that the error bars for *g* are especially large, sometimes encompassing significant proportions of the [0, 2*π*] range, confirming our previous statement that the boundary conditions do not play a significant role in determining the steady state for moderate disorder.

**FIG. T7.**
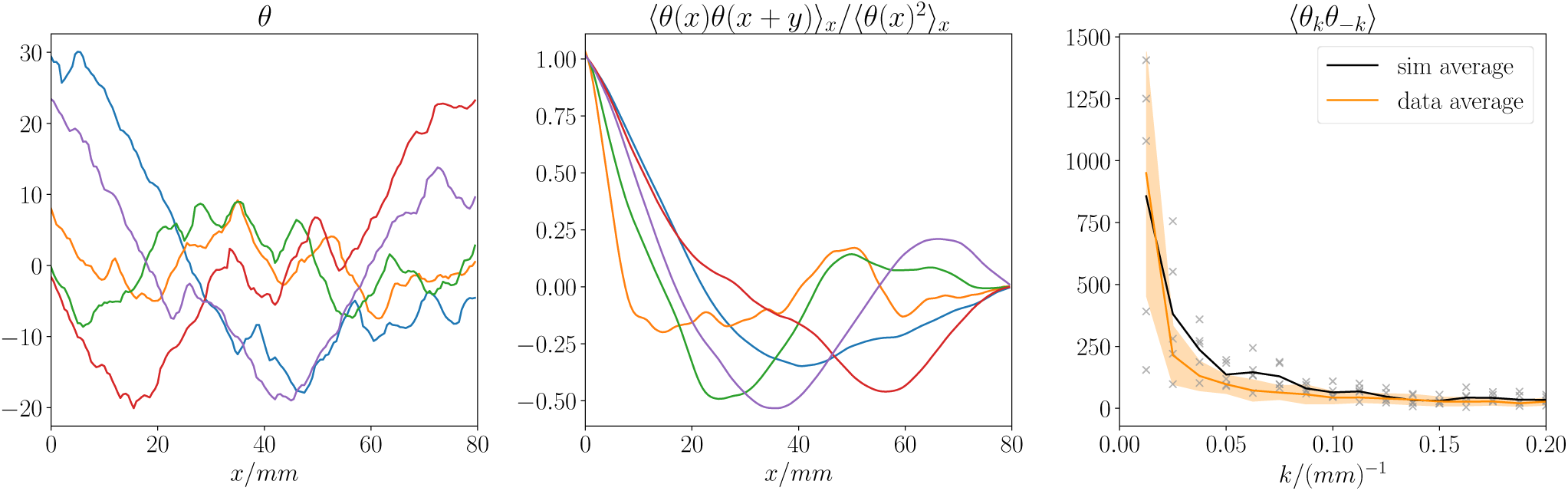
Phase variation (left) of the classic Sakaguchi-Kuramoto model obtained from five simulations using fit parameters obtained from the MLE values of *η* = −0.44 and *σ* = 0.45 with different realizations of static disorder. The corresponding autocorrelation function (middle) and power spectrum (right) are also shown. In the right panel, data from each simulation run are represented by grey crosses. The solid lines mark the average of data and simulations while the orange area marks the standard errors of data.

Based on these findings, we then questioned whether we could reproduce the experimental-like dynamics from simulations. With the ML values *η* = −0.44 and *σ* = 0.45, and fixed gradients of (−1.3, 1.3) at the boundaries, five steady-state *θ* profiles are shown in Figure T7. All simulated steady states show irregularities and reversals, qualitatively similar to features seen in the experimental data in Figure T5. However, only three out of the five simulations in Figure T7 display prominent V-shape patterns and the short length-scale characteristics of the *θ* variations do not closely resemble the experimental data.

Before closing this section, it is prudent to reflect on possible sources of error in the parameter estimations. One potential source of error is in the phase measurements themselves. As noted before, each measurement has an error of around 0.1 in the phase variable, which would only change the *σ* estimate by around 0.02. Thus we do not expect measurement error to bring about major changes in the dynamics. Second, the lack of synchronization may break the inference scheme, compromising the ability to obtain the *true* best-fit parameters. However, as seen in Figure T4, the inference scheme performs moderately well even in regimes that are not fully synchronized. While the lack of synchronization makes the inference scheme less accurate, we would not expect it to alter significantly the parameter estimations.

Thus, the discrepancy between the simulations at MLE parameters and the experimental data indicates that the simplest possible model – the classic Sakaguchi-Kuramoto model – is unable to simultaneously capture the overall V-shape and the high degree of local phase variation (jaggedness). As we can see from Figure T3, with the same *η* value, while the V-shape is more prominent at low *σ*, the local variations in *θ* are much reduced. The MLE estimates represent the “best” compromise between these two opposing factors within the constraint of the classic Sakaguchi-Kuramoto model. We therefore questioned whether potential refinements of the model could resolve the apparent discrepancy between theory and experiment.

#### D. Refinements of the Sakaguchi-Kuramoto model

In arriving at the classic Sakaguchi-Kuramoto model, at least two simplifying assumptions were made: (1) the dynamics are effectively one-dimensional, and (2) the spatio-temporal variations of the natural frequencies are much smaller than the quenched disorder. We therefore explored the consequences of relaxing these assumptions.

**FIG. T8.**
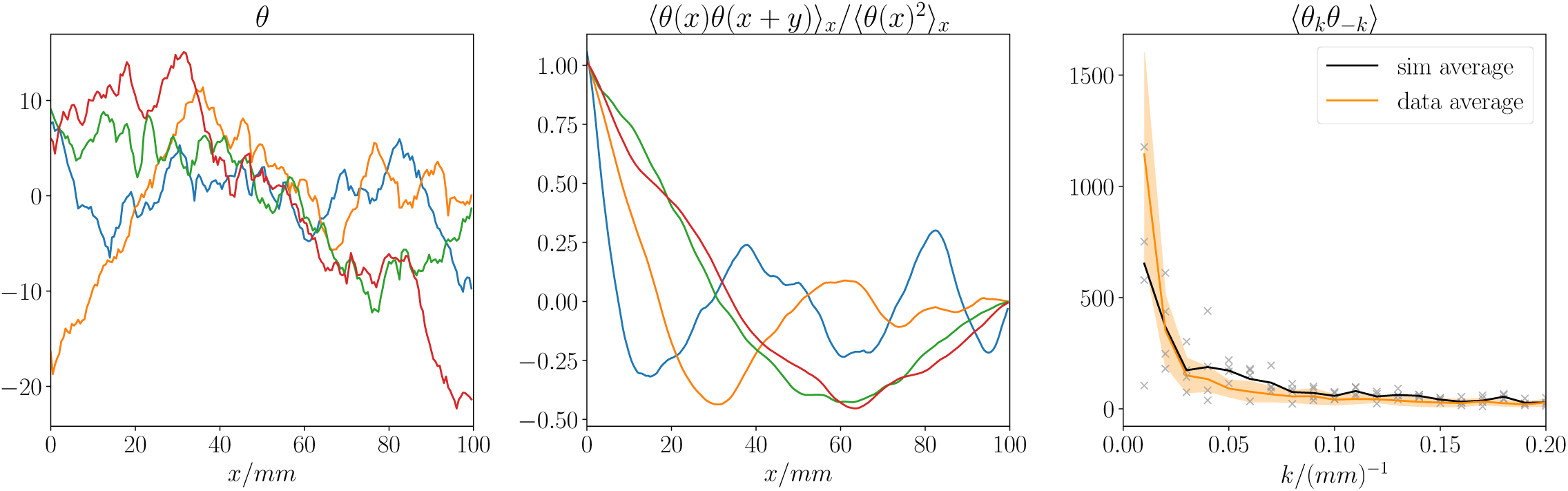
Simulations of the classic Sakaguchi-Kuramoto model with next-nearest-neighbor coupling, mimicking the 2nd dimension of the quasi 1D seminiferous tubules. at the MLE values of *σ* = 0.93, *η* = −0.59, and *α* = 0.3, where *α* denotes the strength of the next-nearest-neighbor coupling as a ratio of coupling. The corresponding autocorrelation function (middle) and power spectrum (right) are also shown. In the right panel, data from each simulation run are represented by grey crosses. The solid lines mark the average of data and simulations while the orange area marks the standard errors of data.

##### 1. The classic Sakaguchi-Kuramoto model in quasi-1D

With the coarse-graining chosen for the data in Figure T5, the lattice spacing was around 500 *µm* in physical units. In comparison, the circumference of the tubule is approximately 600 *µm*, meaning that the extent of the second dimension is comparable to the lattice spacing. To examine the potential impact of the second dimension on the phase dynamics, we performed simulations in a quasi-one-dimensional lattice of size 200 × 2 units with periodic boundary conditions in the second dimension and fixed-gradient boundary conditions in the first dimension, as before. The addition of the extra dimension effectively enhances the interactions between the oscillators and hence leads to a more prominent V-shape.

Since histological measurements of the phase data were only available along a one-dimensional path along the tubules, to fit the model, we chose to mimic the effect of the second dimension using next-nearest-neighbor interactions, reducing the effective dynamics back to 1D. We devised an inference scheme similar to the one in section T1 B and simulated the steady state at MLE parameters, as shown in Figure T8. While the new model captured the variations better than the classic Sakaguchi-Kuramoto model, it was still unable to capture the global V-shape at the optimal fit parameters. Thus, we concluded that the apparent discrepancy between the classic Sakaguchi-Kuramoto model and the data could not be explained by the two-dimensionality of the biological system.

##### 2. Stochastic Sakaguchi-Kuramoto model

Previously, to explore the dynamics of the Sakaguchi-Kuramoto-type models, we focused on a model of static disorder in which local variations in the natural oscillator frequencies do not change over time. However, we reasoned that, due to the changing environment of the seminiferous tubules, the natural frequencies *ω_i_* may be dynamic, preventing the system from ever reaching a true equilibrium steady-state (regardless of whether the system is in a parameter regime when phase synchronization of the static model is possible). To model the potential impact of dynamic fluctuations in the local frequencies, we added a spatio-temporal noise term to Eq. (T1), setting

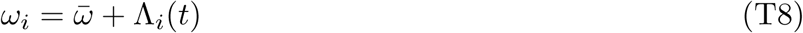

where 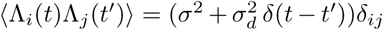 with *σ_d_* denoting the amplitude of the dynamic component of the noise. We note that due to the delta function in time, *σ_d_* does not have the same unit as *σ*: *σ* has a unit of 1/time, whereas *σ_d_* has a unit of 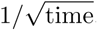.

**FIG. T9.**
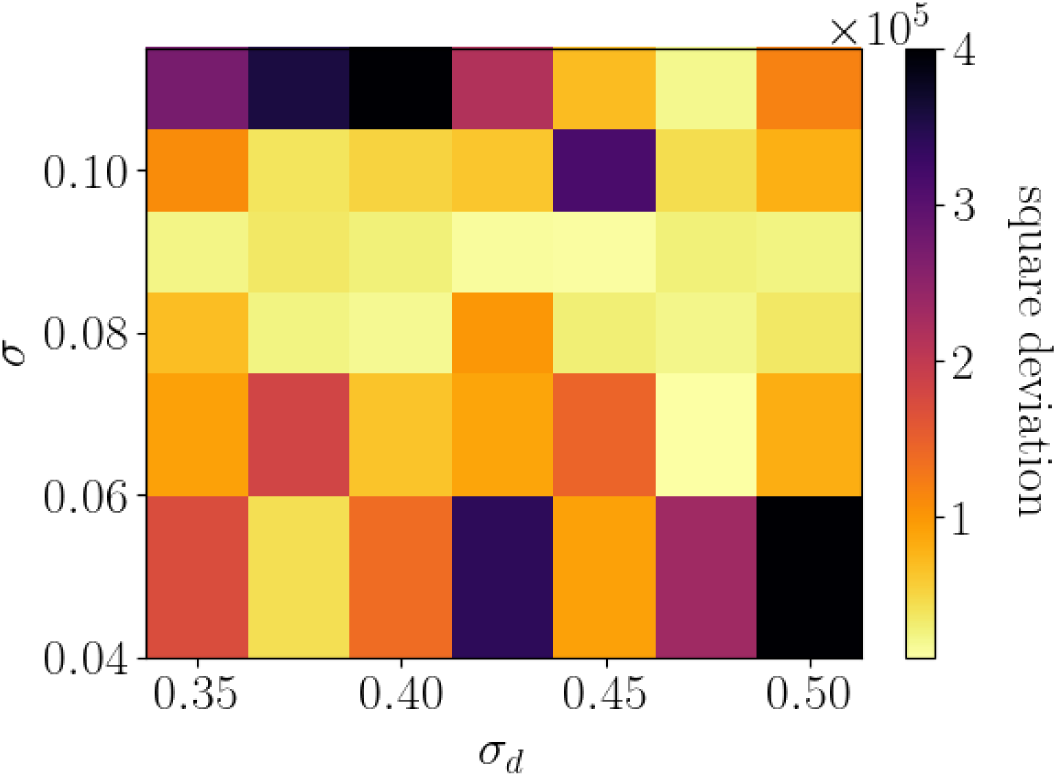
Heatmap in (*σ, σ_d_*)-space of the square deviation of the power spectrum ⟨*θ*(−*k*)*θ*(*k*)⟩ of simulated trajectories from experimental data. *η* = −0.44*, g* = 1.3 for all simulations.

Since the presence of dynamic noise rules out global phase synchronization even at the longest time scales, we can no longer perform maximum likelihood inference to fit the parameters of the model. In fact, inference of such stochastic systems without spatio-temporal data is nearly impossible without exact solutions of the underlying Fokker-Planck equation associated with the stochastic partial differential equation. Thus, we instead performed a two-dimensional scan of the (*σ, σ_d_*) parameter space while restricting *η* = *η*_SK_ where the SK subscript denotes the MLE value obtained from MLE fit to the classic Sakaguchi-Kuramoto model above. To assess the goodness of fit, we used the power spectrum of *θ*, ⟨*θ*(*k*)*θ*(−*k*)⟩, as a measure of the closeness between simulations and data. For each point in (*σ, σ_d_*) parameter space, we compute the total square deviation between the power spectrum of five simulated trajectories and experimental data, as shown in Figure T9. The minimum of the square deviation yields the best-fit values of *σ* = 0.09±0.02*, σ_d_* = 0.44 0.10. Note here that the mean values and error bars are judged by eye from the heatmap to represent the regime of parameters that agree reasonably well with experimental observations and should not be taken as a statistically rigorous estimate.

Simulations using the best-fit parameters are shown in Figure T10. These results show that, with the addition of dynamic disorder, we can consistently capture both the scale of local variations in the phase and the global V-shape pattern observed in experimental data. We note here that, although the best-fit quenched noise (*σ*) value of 0.09 is much smaller than the estimated dynamic noise *σ_d_*, the quenched noise is indispensable in creating the overall V-shape. Without the quenched noise, the overall profile would look more like a wide “U” rather than a “V” (similar to the left-most panel of Figure T3). We further note that the V-shape displayed here is robust to changes in initial conditions, as shown in Figures S7E and T11.

#### E. Temporal dynamics

Finally, to directly compare the phase model with *in vivo* and *ex vivo* kymographs in Figures 5 and S5, we must restore the time dimension by fitting the remaining parameters *ω̅* and *ɛ*, where we recall that *ω̅* is the average of the natural frequencies of the oscillators and *ɛ* was initially set to 1 by rescaling the time coordinate. The total average frequency takes the form

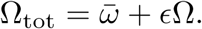

Here Ω_tot_ is the observed net frequency expressed as the sum of the two components – the natural frequency of the bare oscillators and the mean value of the adjustment due to the effect of the collective synchronization of the chain. According to the literature [7, 8], the observed spermatogenic cycles have a period of approximately 8.6 days, implying that Ω_tot_ = 2*π/*(8.6 days). From dimensional analysis (or assuming that *ψ_i_* = *i* × *ϕ* in Eq. (T2)), we have

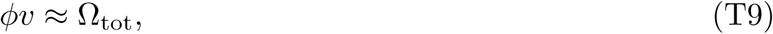

**FIG. T10.**
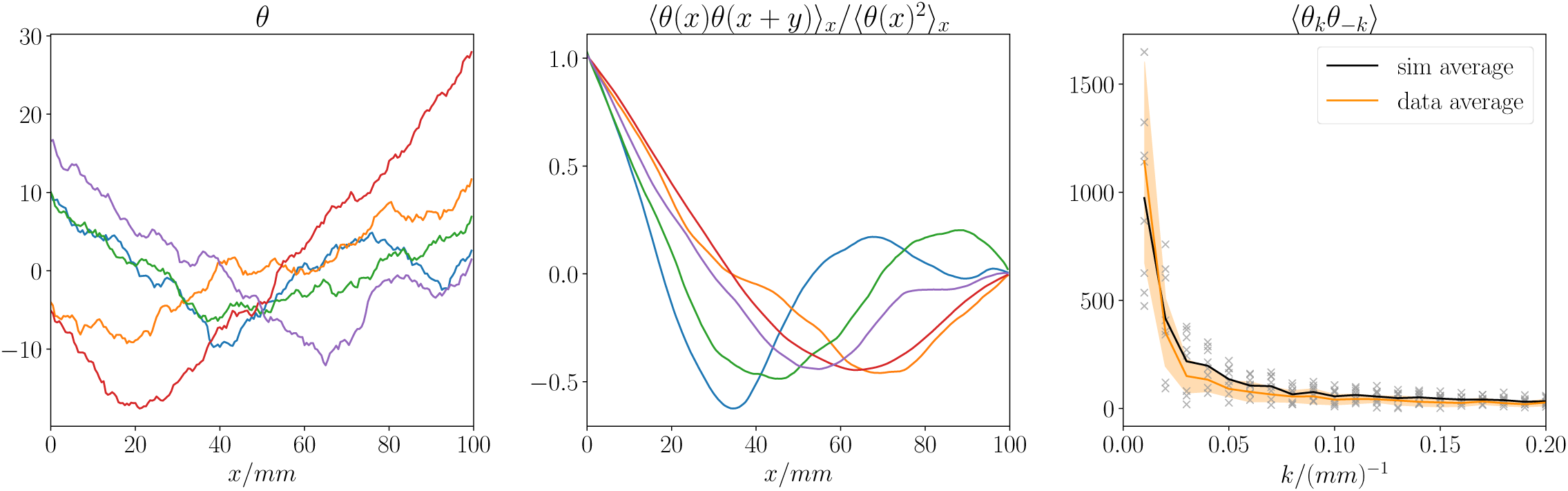
Numerical simulations of the stochastic Sakaguchi-Kuramoto model with spatio-temporal disorder in the local frequencies, with the best-fit parameters of *η* = −0.44, *σ* = 0.09, and *σ_d_* = 0.44. The results show the phase variable (left), its autocorrelation (middle), and power spectrum (right). In the right panel, data from each simulation run are represented by grey crosses. The solid lines mark the average of data and simulations while the orange area marks the standard errors of data. Note that the average power spectrum of simulation runs falls within the error bounds of experimental measurements.

**FIG. T11.**
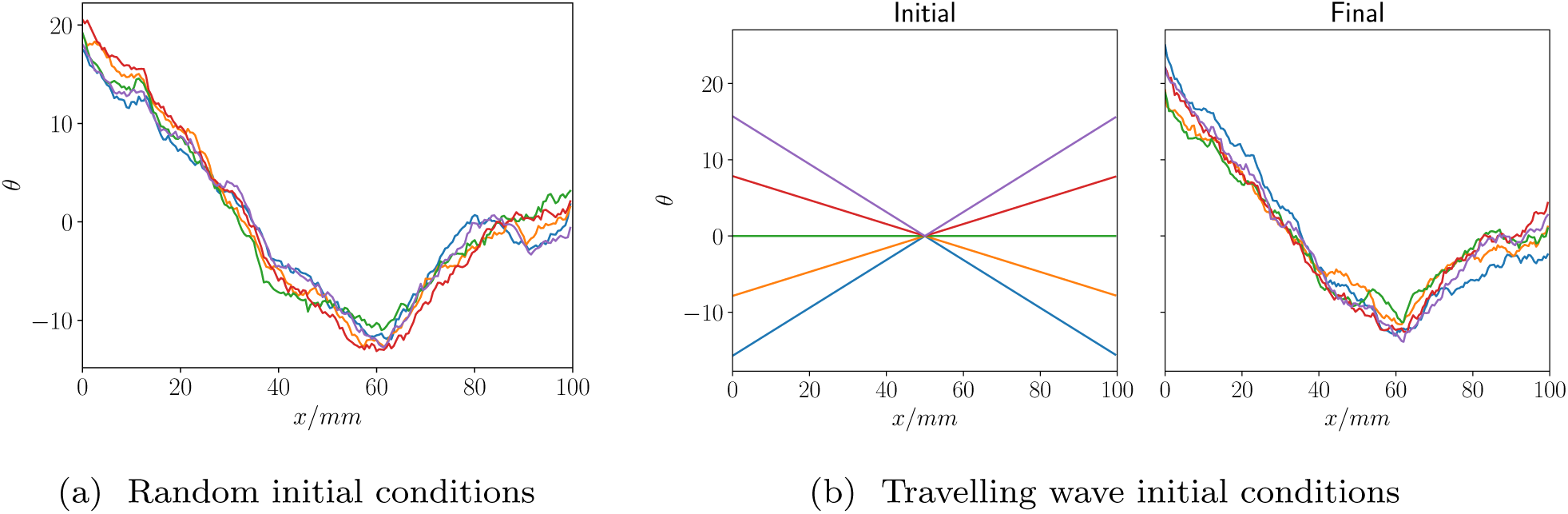
Simulations testing the robustness of the V-shape against initial conditions. All simulations are performed with the same random seed for the static randomness in oscillator frequencies. (a) 10 simulations with random initial conditions; (b) 5 simulations with traveling wave initial conditions, with different velocities of the traveling wave.

where *v* is the wave speed in the kymographs. Measuring the *t*-*x* slope from both *in vivo* and *ex vivo* kymographs (Figures 5 and S5), we arrived at a typical wave speed of approximately (600 *µ*m)/day albeit showing significant variation. The typical *ϕ* value can be measured from the histological data as 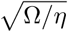, where 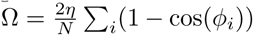, as previously discussed in Section II (the additional spatial-temporal noise does not change the average value of Ω). From the data obtained from the 9 tubules, we found that *ϕ* ≈ 0.5*/*(500 *µm*). Thus, in Eq. (T9), the left-hand side ∼ 0.6 while the right-hand side ∼ 0.7, confirming a remarkable degree of consistency across the experimental datasets.

From the analysis above, it is apparent that the average wave speed alone are not sufficient to fit both *ω̅* and *ɛ* parameters since they do not provide additional information beyond the histological measurements that we have already used. The simplest assumption (without further temporal data) is that *ω̅* = Ω_tot_ and that *ɛ* is sufficiently small to not alter the overall frequency significantly. An example of a simulated spatial-temporal kymograph is shown in Figure T12. As discussed in the main text, the simulated kymograph shows qualitative agreements with the experimental data based on live imaging. We additionally note that the disappearance of modulation can happen for two reasons: first, as a transient before a dynamic long-term steady-state is reached or, second, at a position where there is a significant change in the natural oscillator frequency due to the dynamic disorder.

**FIG. T12.**
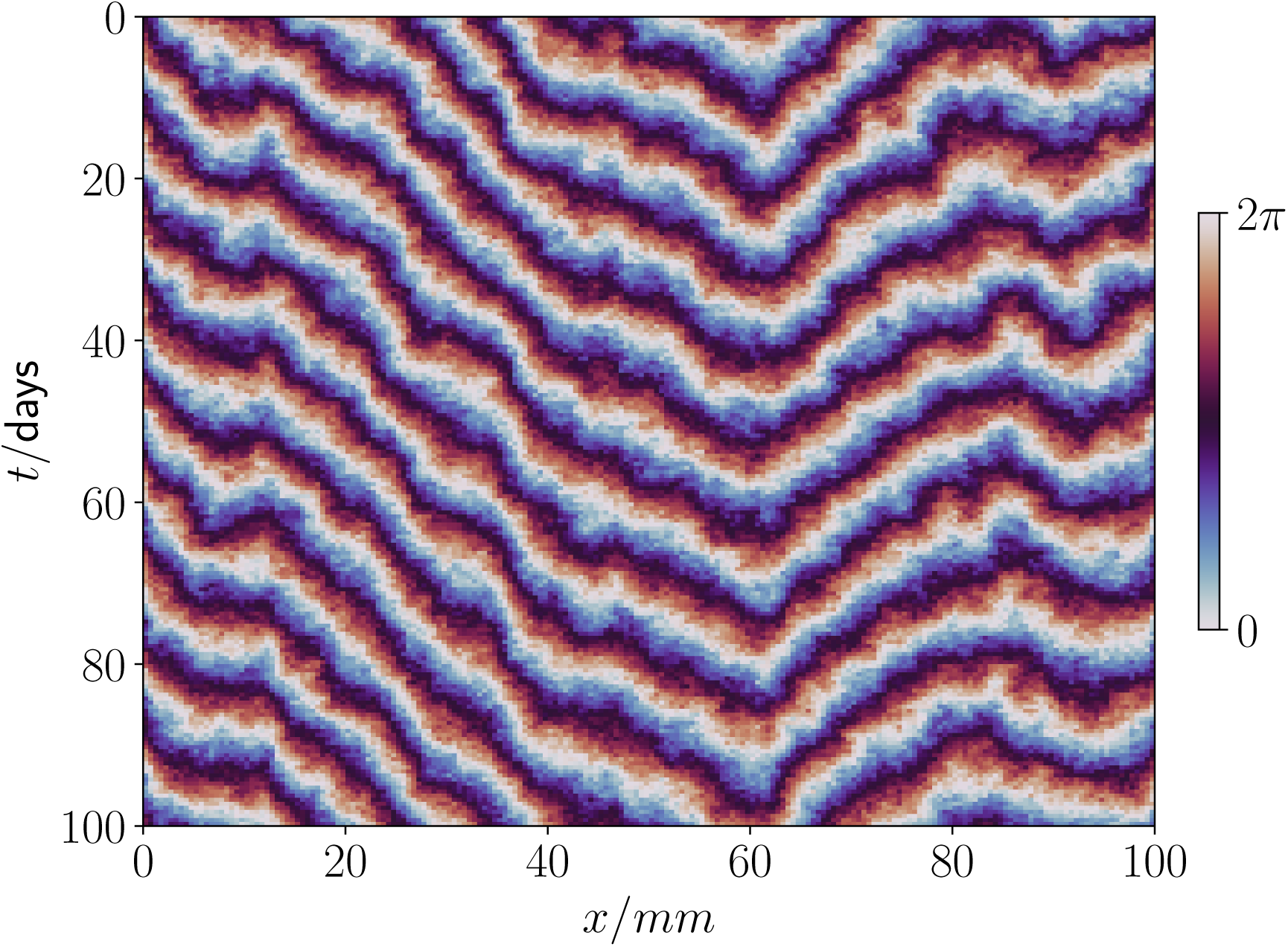
Kymograph of a simulation of the Sakaguchi-Kuramoto model with both static and dynamic disorder. Discrete lattice sites with lattice spacing 500 *µm* were used along the *x* axis, in line with the histological measurements shown in Figure T5. The model parameters are *η* = 0.44, *σ/ɛ* = 0.09, 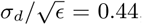, *ω̅* = 2*π/*(8.6 days), and *ɛ* = 1*/*(5 days).

Overall, based on our analyses, the results show that the Sakaguchi-Kuramoto model with stochastically varying frequencies and asymmetric coupling provides a plausible description of the experimental data, both qualitatively and quantitatively. Importantly, this minimal model can explain robust features of the data, including the emergence of phase waves and the sporadic appearance of reversals of the phase velocity that overlay the hallmark V-shape pattern of the phase profile along the tubule.

### T2. MINIMAL THEORY OF THE SPERMATOGENIC CYCLE DRIVEN BY RA PATHWAY

In section T1, we demonstrated that the properties of the spermatogenic waves observed at the length scale of the seminiferous tubules can emerge through the nearest-neighbor phase-coupling of locally formed autonomous oscillators. To do so, based on the stochastic Sakaguchi-Kuramoto model, we considered a phenomenological description, setting aside potential mechanisms that could mediate the genesis of the phase oscillator and its coupling. Here, we explore one plausible candidate for the origin of the robust spermatogenic cycles forming in the seminiferous epithelium. To this end, motivated by insights gained from experimental observations, we will synthesize a minimal model that incorporates feedback regulations between the spermatogenic stem cells and their differentiating progeny, mediated by the RA metabolism and function, as illustrated in Figure T13.

As detailed in the main text, it has been established that the RA signal plays a crucial role in generating and regulating the spermatogenic cycle and wave. However, the underlying mechanisms remain elusive from a systems biology perspective. The primary role of RA is to induce commitment of the spermatogenic stem cells for differentiation towards matured sperm. At the same time, the production of RA is promoted by the late pachytene spermatocyte (LP) stage of germ cell differentiation (which occurs after a maturation time *τ* representing the time taken to transition from the stem cell compartment to the LP stage), and inhibited by cells at the round spermatid (RT) stage (after a maturation time of *τ* + *ξ* to become RT) [9, 10]. Together, these processes comprise time-delayed feedback loops for the RA concentration, which creates the conditions to induce autonomous oscillations in the form of a limit cycle [11]. At the mean-field level, the dynamics of the RA concentration can be written as

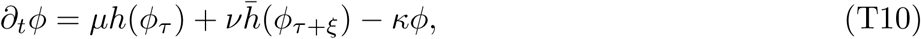

where *h*(*x*) = *x^n^/*(1 + *x^n^*) denotes a Hill-type function with *h̅*(*x*) = 1 − *h*(*x*), *µ*, *ν*, and *κ* are constants, and *ϕ_τ_* and *ϕ_τ_*_+*ξ*_ indicate the RA concentration of the field at time *t − τ* and *t* − (*τ* +*ξ*), respectively. The first term corresponds to positive feedback from the LP stage (with the time-delay *τ* from when stem cells enter into a differentiation program) with dose-response properties represented by the Hill-type function. Similarly, the second term indicates the negative feedback from the RT stage occurring at time *τ* + *ξ*, while the third term represents the natural decay of RA. Note that, here, we have rescaled *ϕ* by the half-maximum concentration of the Hill functions, with which differentiation-committed cells that regulate the RA production after time delays are generated.

**FIG. T13.**
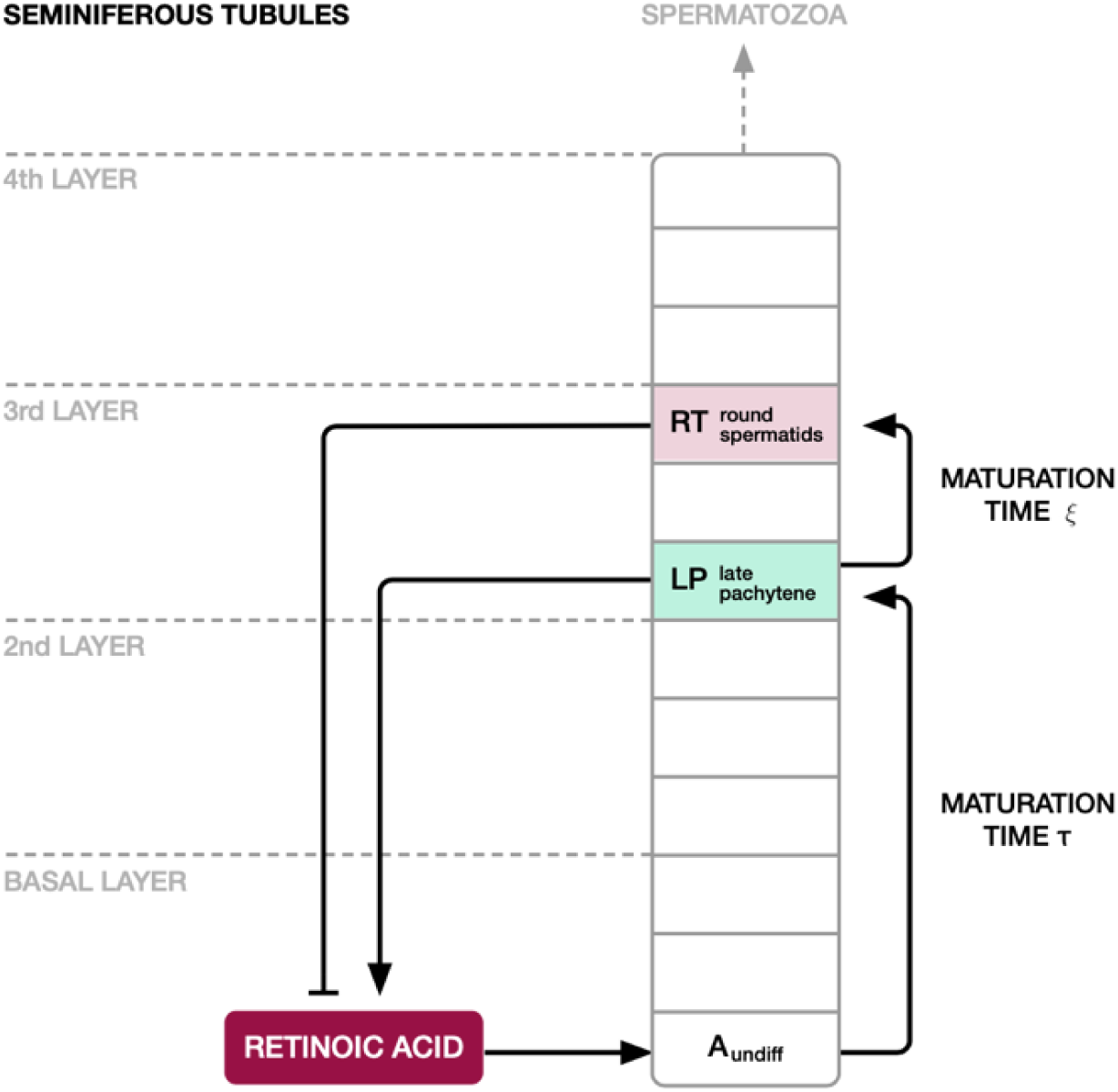
Diagrammatic illustration of the key regulatory circuit that controls the RA concentration in the mouse seminiferous epithelium. For further details, see the main text and main text of Supplementary Theory. The commitment of stem cells (Aundiff) to transit to the differentiation toward spermatozoa is coordinated tightly with the strength of the RA signal (i.e., the local RA concentration). During the differentiation processes, cells experience the steps of LP (late pachytene spermatocyte) and RT (round spermatid), which respectively up- and down-regulate the RA production. These regulations form positive and negative feedback loops with time delay.

To begin, since the formation of time-delayed feedback loops fulfills the requisite of forming a limit cycle oscillator, we question whether this modeling scheme can support limit cycle oscillations. We first investigate analytically the potential onset of an instability towards oscillatory dynamics of the model using a bifurcation analysis. Suppose that the system has a uniform steady state *ϕ*(*t*) = *ϕ̅*. The uniform state must satisfy the relation

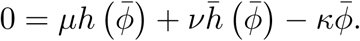

Hereafter, we set *f* (*ϕ*) = *µh* (*ϕ̅*)+ *νh̅* (*ϕ̅ κϕ̅*), and use *f* ^′^ as a shorthand for *∂_ϕ_f _ϕ_*_=*ϕ̅*_. For simplicity, we will focus on the case with one unique uniform steady state, corresponding to negative *f* ^′^, which can be achieved if we set *µ* ≤ *ν*.

Now consider a small perturbation around the uniform steady state, *ϕ*(*t*) = *ϕ̅* + *s*(*t*). Expanding the dynamical equation (T10) to leading order in *s*(*t*),

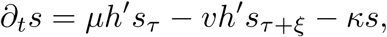

where *h*^′^ ≡ *h*^′^(*ϕ̅*), and substituting the ansatz *s*(*t*) = *s*_0_*e^λt^*, we obtain the relation

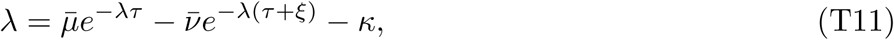

where *µ̅* = *µh*^′^ and *ν̅* = *νh*^′^. If there is a solution for *λ* that is real and negative, the uniform state is stable and there are no oscillations. Since when *λ* = 0, the right-hand side is negative (as discussed previously, the uniform state *ϕ̅* is stable against temporally uniform perturbations), for the right-hand and left-hand side to be equal at negative *λ*, the exponential growth for negative *λ* must be extremely slow, which requires *τ, ξ* ≪ 1. However, this is not the case we are interested in – we expect the maturation times *τ* and *ξ* to be much larger than the typical degradation time. Therefore, we turn to complex solutions for *λ*.

Setting *λ* = *λ_R_* + *iλ_I_*,

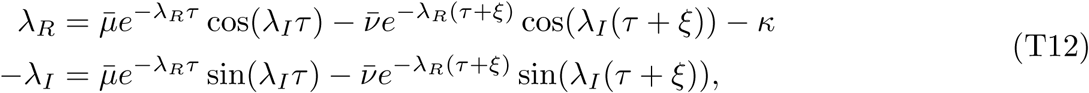

one may note that when *ξ* = 0 and *τ* = 0, *λ* is real and negative. Additionally, it has been shown that along either axis, *λ* is a continuous function (in the complex plane), and that *λ_R_* follows the uppermost branch of a family of Lambert W-functions, which means that it either goes from negative to positive with a single intersection or stays negative throughout [11]. Although no analytical results are known for the shape of *λ_R_* in both variables, we can reasonably assume that the *λ_R_*(*τ, ξ*) is well-behaved and continuous, and there is a continuous line of thresholds that marks the onset of instability. From numerical simulations, we find that this is indeed the case, as shown in Figure T14. Thus, to determine the true line at which there is an onset of instability, we need to find the family of solutions of (*τ, ξ*) at which *λ_R_* = 0.

Setting *λ_R_* = 0 in Eq. (T12), we obtain a much simpler complex equation,

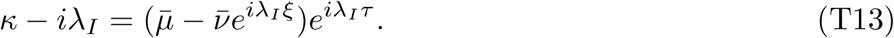

Equating the modulus,

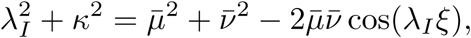

we find that there are multiple possible values of *λ_I_*for a given *ξ* due to the cosine term on the right-hand side. Nonetheless, we can numerically solve for the roots and then substitute back into the following equation of the complex argument to obtain *τ*,

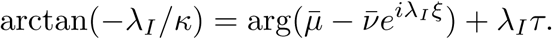

The multiple solutions of *λ_I_* correspond to multiple solutions of *τ* for each *ξ*. All the solutions are plotted in Figure T14. We conclude that the numerical solutions of the much simpler equation (T13) can capture the onset of oscillations in the (*τ, ξ*) plane.

**FIG. T14.**
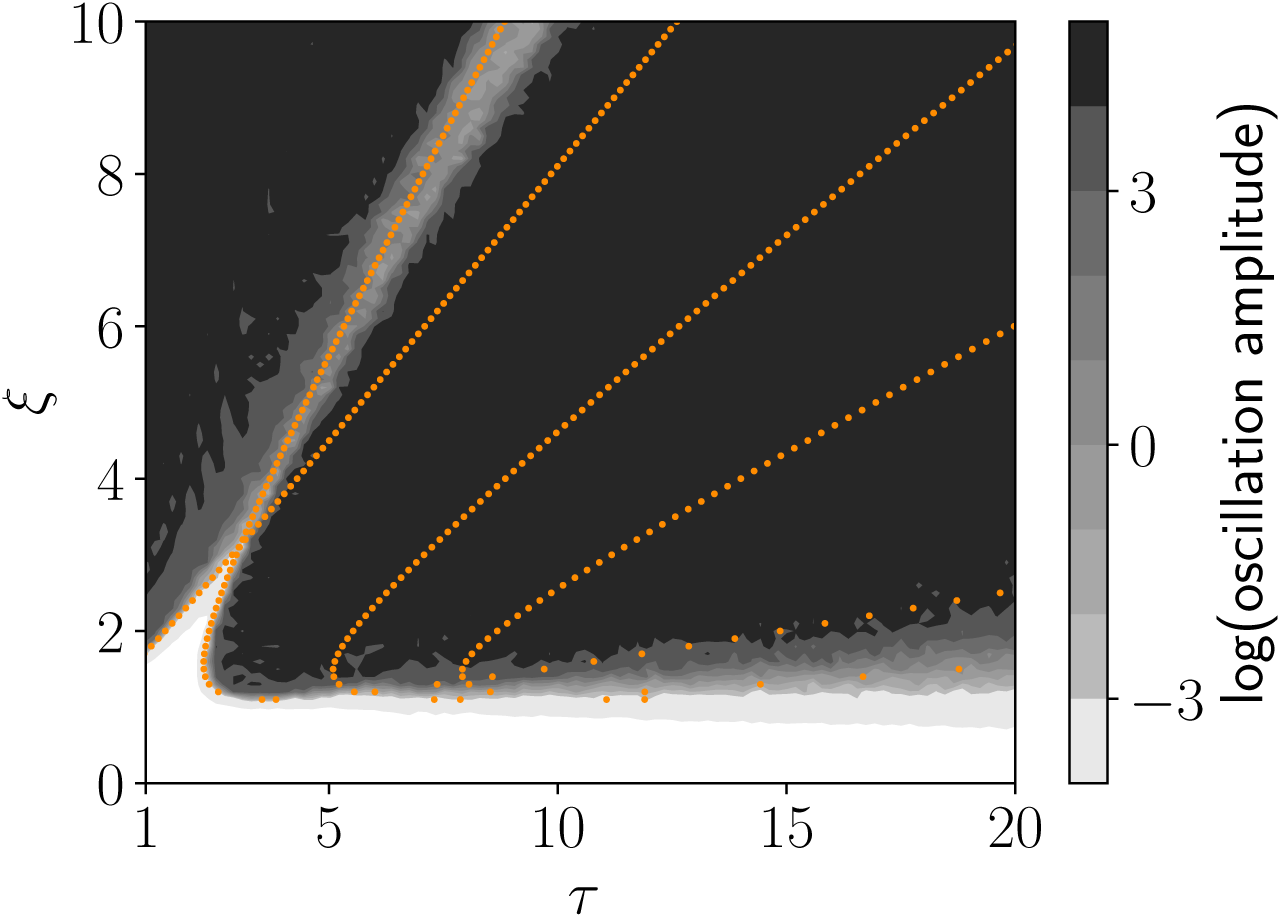
Contour map showing the amplitude of oscillations obtain from the numerical solution of the dynamical equation (T10) for each (*τ, ξ*) value. The orange dotted lines are all numerical solutions of Eq. (T13). The true line of onset of oscillations corresponds to where the patch extending from the origin meets the various solutions. The remaining parameters are *ν* = *µ* = 1.2*, κ* = 1*, n* = 8

**FIG. T15.**
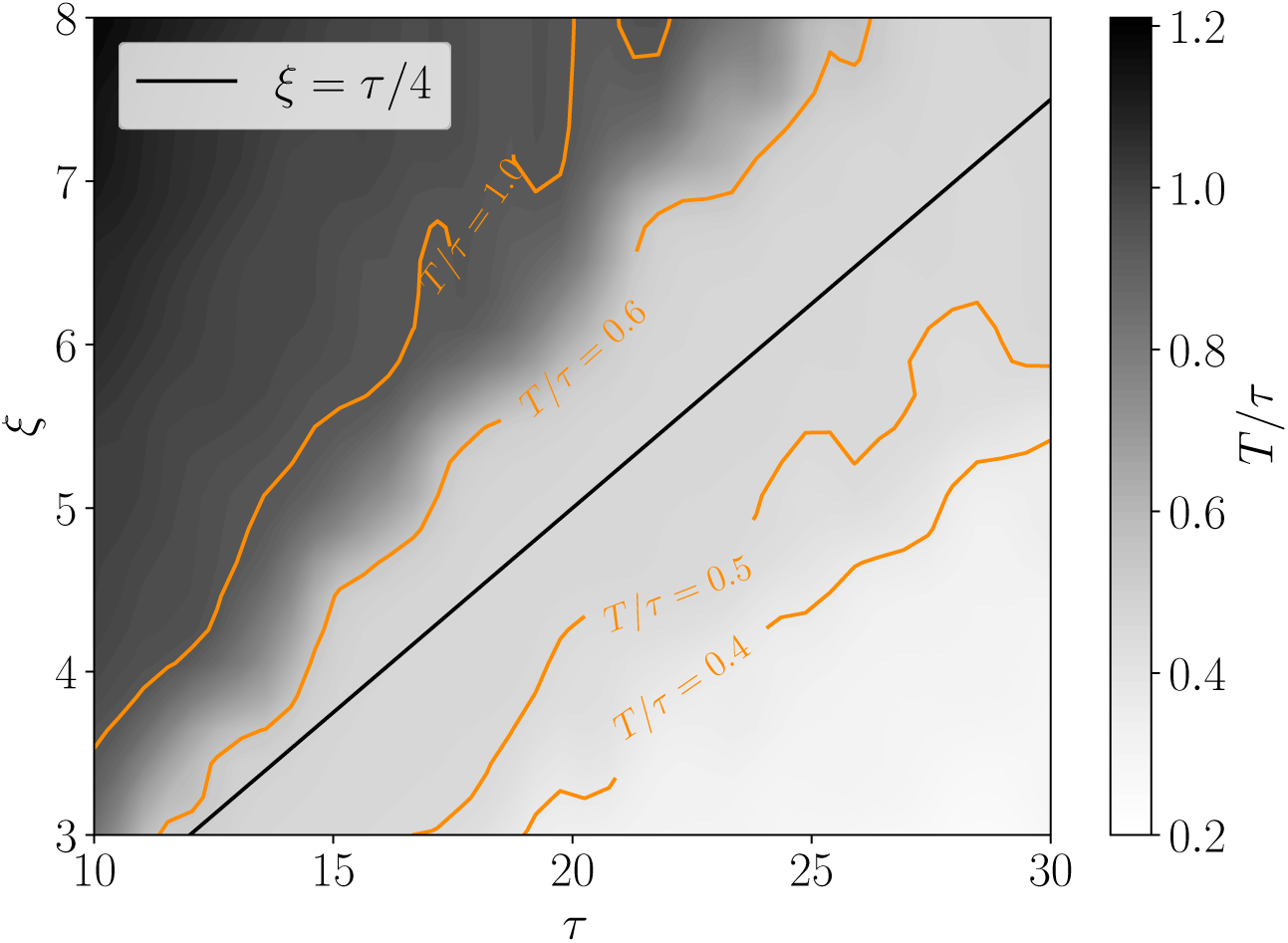
The period of the oscillation cycles of the dynamical equation (T10) in the *ξ − τ* plane. For details of the shading and color coding, we refer to the text. The remaining parameters are *ν* = *µ* = 1.2*, κ* = 1*, n* = 8

Having established that cyclic behaviors emerge naturally from the delayed feedback model of RA concentration, we next pinpoint the operating parameter space for the actual dimension of the spermatogenic cycle within the seminiferous epithelium. Experimental observations based on staging the spermatogenic cycle indicate that it takes around 2 cycles for stem cells (Aundiff) to differentiate into LP expressing a high level of RA synthesizing enzyme (Aldh1a2), and approximately 2.5 cycles for stem cells to differentiate into RT expressing enzymes inhibiting the RA synthesis, corresponding to *τ* and *τ* +*ξ*, respectively [9, 10]. Thus, the ratio of *ξ* to *τ* is approximately 0.25, with the ratio of the cycle period *T* to *τ* at around 0.5, as illustrated in Figure T13. The former constraints are outlined as the solid straight line and the latter constraints are represented by the orange contours (showing the area with ratios of 0.4, 0.5, and 0.6) in Figure T15. The overlap between the two sets of constraints is the operating parameter space for generating limit cycle oscillators with observed cycle periodicity of the spermatogenic cycle (*T* = *τ/*2).

In summary, the analysis of the onset of oscillatory solutions of the delayed-feedback dynamics of the RA concentration model confirms the existence of a parameter subspace that gives rise to biologically relevant cycle lengths.

### T3. FROM THE RA CONCENTRATION THEORY TO A SAKAGUCHI-KURAMOTO-TYPE MODEL

In section T2, we saw that the delayed feedback regulation by RA signals is capable of generating a limit cycle oscillator, in a manner compatible with the key observed properties of the spermatogenic cycle. Then, we ask whether this RA-based oscillation model, when extended in one-dimensional geometry, can lead to the emergence of the dynamics of phase-coupled oscillators that we observed in section T1 based on the stochastic Sakaguchi-Kuramoto model.

While, as emphasized in the main text, this analysis serves only as a proof of principle, showing the potential of the RA pathway to mediate oscillator coupling between neighboring regions, without pretending to comprise a complete mechanistic theory of the coupling and local dynamics. Indeed, the microscopic theory is presented here to demonstrate that a more biologically inspired theory can give rise to the same qualitative behavior as the stochastic Sakaguchi-Kuramoto Model.

A natural way of extending the local oscillator theory to the phase-coupled one-dimension array is to include feedback from both the LP and RT stages of the neighboring sites with time delay. Within the framework of the feedback model, such behavior can be encapsulated by the dynamics

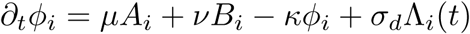

Where

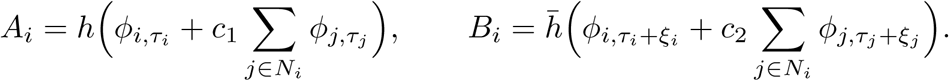

and Λ*_i_*(*t*) are independent spatio-temporal noises at each site, which play the role of dynamic noises in the stochastic SK model. Here *c*_1_ and *c*_2_ denote the coupling strength between neighboring sites, representing the fact that the RA concentration at a certain location is not only affected by feedback from germ cells within the particular regions but also from those at neighboring sites. We also denote the standard deviations of {*τ_i_*} as *σ_τ_* and the standard deviation of {*ξ_i_*} as *σ_ξ_* to represent the scale of static fluctuations in the natural frequencies, similar to the role of the static noise *σ* in the stochastic Sakaguchi-Kuramoto model. We emphasize that the *σ_τ_, σ_ξ_* parameterize static noises as the {*τ_i_*} and {*ξ_i_*} distributions are fixed over time, as opposed to Λ*_i_*(*t*), which vary dynamically over time.

Within the context of the RA-based 1D model, it is difficult to disentangle the amount of nonreciprocity (characterized by the strength of *η* in Sakaguchi-Kuramoto-type models) from the interaction strength (*ɛ*, as defined in section T1), as the non-reciprocity in the coupling is naturally encoded in the saturation effect of the Hill functions. However, we can trace the origin of the non-reciprocity *qualitatively* and determine the sign of *η*. For example, consider the scenario where only negative feedbacks are coupled (*c*_1_ = 0, *c*_2_ ≠ 0): if there is strong negative feedback locally, adding more negative contributions from neighbors does not have an effect due to the saturation of the Hill function whereas, if there is no negative feedback locally, adding negative feedback from neighbors has a strong effect. This creates an asymmetry between the interactions of neighboring oscillators – the lagging cycle has a larger effect on the leading cycle, as the negative feedback of the lagging cycle is able to inhibit the entrance of the leading cycle, equivalent to setting *η* negative in the Sakaguchi-Kuramoto-type models. Note that we do not conclude that this model exclusively explains the mechanism of non-reciprocal coupling, but there are other possible mechanisms such as check-pointing of cell differentiation progression and spermatogenic stage-related fluctuation in the number of differentiation-primed stem cells [12]. This issue goes beyond the scope of this paper.

**FIG. T16.**
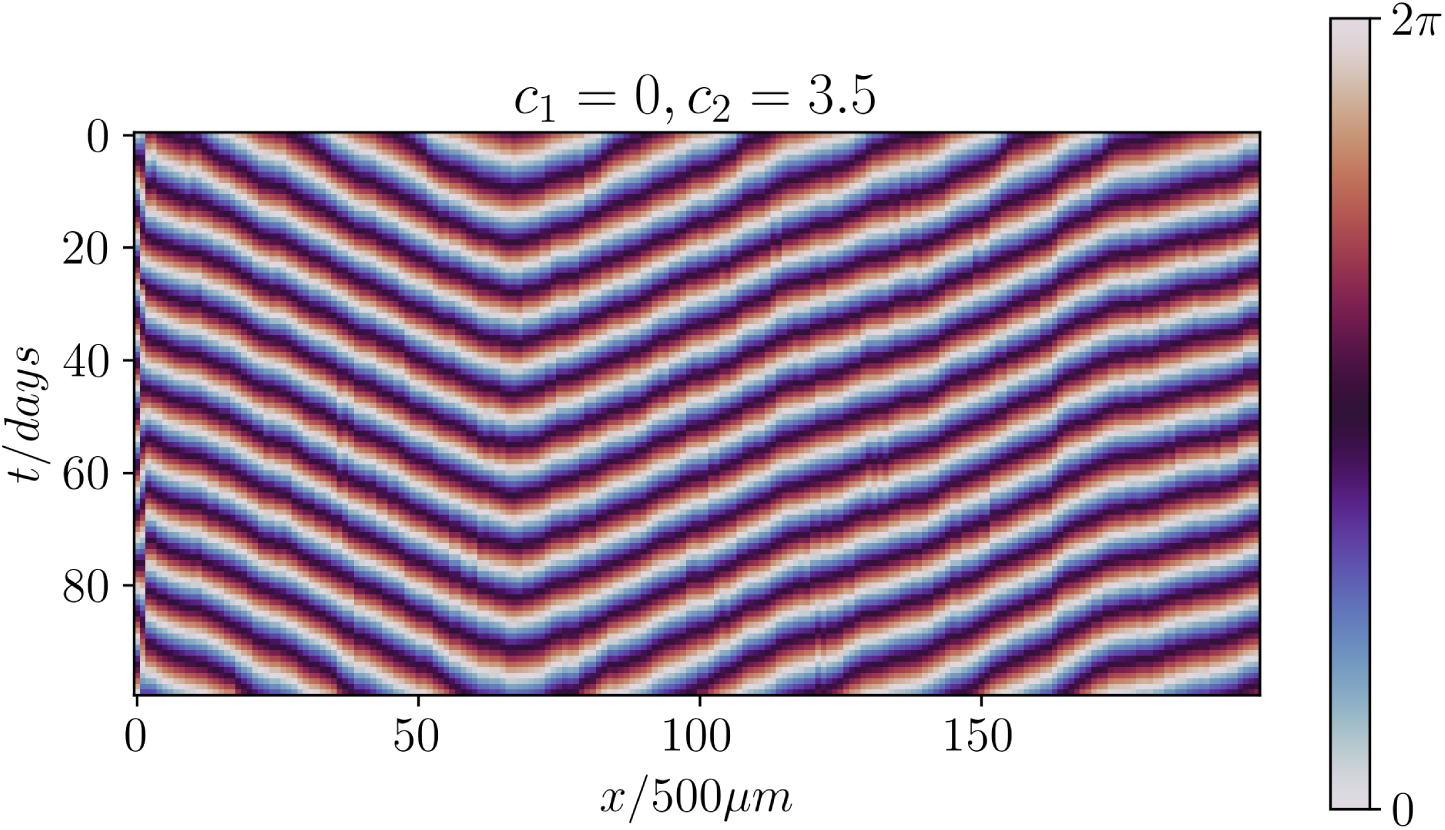
Kymograph of the spatial-temporal evolution of the extended RA concentration model in one-dimension with *µ* = *ν* = 1, *κ* = 1, *c*_1_ = 0, *c*_2_ = 3.5, *τ* = 17.2 days, *ξ* = 4.3 days, *σ_τ_* = 0.01, *σ_ξ_* = 0 and *σ_d_* = 0.1. Note here *µ, ν* are smaller than the local RA concentration model to account for the feedback of neighboring cycles.

As shown in Figure T16, there is clear evidence of phase waves present in the system. Additionally, we found that the phase waves are robust to variations in the dynamic noise *σ_d_*within reasonable limits, similar to the observations in the stochastic Sakaguchi-Kuramoto model. By scanning the parameters of the model, we obtained a regime in parameter space where the phase profile also exhibits an overall rugged V-shape consistently (see Figure T16). We note that, due to the high dimensionality of the parameter space and the lack of a direct handle on the amount of non-reciprocity, the parameter search here is tedious and time-consuming, compared to the simplicity of the stochastic Sakaguchi-Kuramoto Model. This supports our initial decision to investigate minimal rather than microscopically-inspired complex models.

In conclusion, motivated by experimental observations indicating the key roles of RA in forming the spermatogenic wave, we could present that a minimal microscopic model explains the phase-wave dynamics of mouse spermatogenesis. The model is a natural extension of the local delayed-feedback model for RA concentration, involving non-reciprocity encoded in the saturation effect of the Hill functions that govern nearest-neighbor interactions. Further, through a comprehensive parameter scan, we found an area in parameter space that is capable of robustly producing the macroscopic V-shaped phase organization, similar to that of the stochastic Sakaguchi-Kuramoto model and well explaining the measured data from experiments.

## References

1. Russell, L.D. (1990). Histological and histopathological evaluation of the testis (Cache River Press).

2. Yoshida, S. (2019). Chapter Seven - Heterogeneous, dynamic, and stochastic nature of mammalian spermatogenic stem cells. In Current Topics in Developmental Biology The Immortal Germline., R. Lehmann, ed. (Academic Press), pp. 245–285. 10.1016/bs.ctdb.2019.04.008.

3. Cordeiro, D.A.J., Costa, G.M.J., and França, L.R. de (2021). Testis structure, duration of spermatogenesis and daily sperm production in four wild cricetid rodent species (A. cursor, A. montensis, N. lasiurus, and O. nigripes). PLOS ONE 16, e0251256. 10.1371/journal.pone.0251256.

4. Russell, L.D. (1977). Movement of spermatocytes from the basal to the adluminal compartment of the rat testis. Am. J. Anat. 148, 313–328. 10.1002/aja.1001480303.

5. Smith, B.E., and Braun, R.E. (2012). Germ Cell Migration Across Sertoli Cell Tight Junctions. Science 338, 798–802. 10.1126/science.1219969.

6. Moens, P.B., and Go, V.L.W. (1972). Intercellular bridges and division patterns of rat spermatogonia. Z.Zellforsch 127, 201–208. 10.1007/BF00306802.

7. Ren, H.P., and Russell, L.D. (1991). Clonal development of interconnected germ cells in the rat and its relationship to the segmental and subsegmental organization of spermatogenesis. Am. J. Anat. 192, 121–128. 10.1002/aja.1001920203.

8. Ebner, V. von (1871). Untersuchungen über den Bau der Samencanälchen und die Entwicklung der Spermatozoiden bei den Säugethieren und beim Menschen Electronic ed. (Engelmann).

9. Leblond, C.P., and Clermont, Y. (1952). Definition of the Stages of the Cycle of the Seminiferous Epithelium in the Rat. Annals of the New York Academy of Sciences 55, 548–573. 10.1111/j.1749-6632.1952.tb26576.x.

10. Oakberg, E.F. (1956). A description of spermiogenesis in the mouse and its use in analysis of the cycle of the seminiferous epithelium and germ cell renewal. Am. J. Anat. 99, 391–413. 10.1002/aja.1000990303.

11. Oakberg, E.F. (1956). Duration of spermatogenesis in the mouse and timing of stages of the cycle of the seminiferous epithelium. Am. J. Anat. 99, 507–516. 10.1002/aja.1000990307.

12. Clermont, Y., and Trott, M. (1969). Duration of the Cycle of the Seminiferous Epithelium in the Mouse and Hamster Determined by Means of 3H-Thymidine and Radioautography. Fertility and Sterility 20, 805–817. 10.1016/S0015-0282(16)37153-9.

13. Elftman, H. (1950). The Sertoli cell cycle in the mouse. The Anatomical Record 106, 381–393. 10.1002/ar.1091060306.

14. Ueno, H., and Mori, H. (1990). Morphometrical Analysis of Sertoli Cell Ultrastructure during the Seminiferous Epithelial Cycle in Rats. Biol Reprod 43, 769–776. 10.1095/biolreprod43.5.769.

15. França, L.R. de, Ghosh, S., Ye, S.-J., and Russell, L.D. (1993). Surface and Surface-to-Volume Relationships of the Sertoli Cell during the Cycle of the Seminiferous Epithelium in the Rat. Biology of Reproduction 49, 1215–1228. 10.1095/biolreprod49.6.1215.

16. Ye, S.-J., Ying, L., Ghosh, S., França, L.R. de, and Russell, L.D. (1993). Sertoli cell cycle: A re-examination of the structural changes during the cycle of the seminiferous epithelium of the rat: SERTOLI CELL CYCLE. Anat. Rec. 237, 187–198. 10.1002/ar.1092370206.

17. Sugimoto, R., Nabeshima, Y., and Yoshida, S. (2012). Retinoic acid metabolism links the periodical differentiation of germ cells with the cycle of Sertoli cells in mouse seminiferous epithelium. Mechanisms of Development 128, 610–624. 10.1016/j.mod.2011.12.003.

18. Chakravorty, A., Simons, B.D., Yoshida, S., and Cai, L. (2024). Spatial Transcriptomics Reveals the Temporal Architecture of the Seminiferous Epithelial Cycle and Precise Sertoli-Germ Synchronization. Preprint at bioRxiv, 10.1101/2024.10.28.620681

19. Regaud, C. (1900). Direction helicoidale du mouvement spermatogenetique dans les tubes seminiferes du rat. Comptes rendus des séances de la Société de biologie et de ses filiales, 1042.

20. Perey, B., Clermont, Y., and Leblond, C.P. (1961). The wave of the seminiferous epithelium in the rat. Am. J. Anat. 108, 47–77. 10.1002/aja.1001080105.

21. Hogarth, C.A., and Griswold, M.D. (2010). The key role of vitamin A in spermatogenesis. J. Clin. Invest. 120, 956–962. 10.1172/JCI41303.

22. Yoshida, S. (2016). From cyst to tubule: innovations in vertebrate spermatogenesis: Innovations in vertebrate spermatogenesis. WIREs Dev Biol 5, 119–131. 10.1002/wdev.204.

23. Nakata, H., Wakayama, T., Sonomura, T., Honma, S., Hatta, T., and Iseki, S. (2015). Three-dimensional structure of seminiferous tubules in the adult mouse. J. Anat. 227, 686–694. 10.1111/joa.12375.

24. Ebner, V. von (1888). Zur Spermatogenese bei den Säugethieren. Archiv f. mikrosk. Anatomie 31, 236–292. 10.1007/BF02955709.

25. Curtis, G.M. (1918). The morphology of the mammalian seminiferous tubule. Am. J. Anat. 24, 339–394. 10.1002/aja.1000240303.

26. Nakata, H., Sonomura, T., and Iseki, S. (2017). Three-dimensional analysis of seminiferous tubules and spermatogenic waves in mice. Reproduction 154, 569–579. 10.1530/REP-17-0391.

27. Griswold, M.D. (2016). Spermatogenesis: The Commitment to Meiosis. Physiological Reviews 96, 1–17. 10.1152/physrev.00013.2015.

28. Yoshida, S. (2018). Open niche regulation of mouse spermatogenic stem cells. Develop. Growth Differ. 60, 542–552. 10.1111/dgd.12574.

29. Hogarth, C.A., Arnold, S., Kent, T., Mitchell, D., Isoherranen, N., and Griswold, M.D. (2015). Processive Pulses of Retinoic Acid Propel Asynchronous and Continuous Murine Sperm Production. Biology of Reproduction 92, 1–11. 10.1095/biolreprod.114.126326.

30. Endo, T., Freinkman, E., de Rooij, D.G., and Page, D.C. (2017). Periodic production of retinoic acid by meiotic and somatic cells coordinates four transitions in mouse spermatogenesis. Proc Natl Acad Sci USA 114, E10132–E10141. 10.1073/pnas.1710837114.

31. Endo, T., Romer, K.A., Anderson, E.L., Baltus, A.E., de Rooij, D.G., and Page, D.C. (2015). Periodic retinoic acid–STRA8 signaling intersects with periodic germ-cell competencies to regulate spermatogenesis. Proc Natl Acad Sci USA 112, E2347–E2356. 10.1073/pnas.1505683112.

32. Morales, C., and Griswold, M.D. (1987). Retinol-Induced Stage Synchronization in Seminiferous Tubules of the Rat. Endocrinology 121, 432–434. 10.1210/endo-121-1-432.

33. Bartlett, J.M.S., Weinbauer, G.F., and Nieschlag, E. (1990). Stability of Spermatogenic Synchronization Achieved by Depletion and Restoration of Vitamin A in Rats. Biol Reprod 42, 603–612. 10.1095/biolreprod42.4.603.

34. Hogarth, C.A., Evanoff, R., Mitchell, D., Kent, T., Small, C., Amory, J.K., and Griswold, M.D. (2013). Turning a Spermatogenic Wave into a Tsunami: Synchronizing Murine Spermatogenesis Using WIN 18,446. Biology of Reproduction 88, 1–9. 10.1095/biolreprod.112.105346.

35. França, L.R. de, Ogawa, T., Avarbock, M.R., Brinster, R.L., and Russell, L.D. (1998). Germ Cell Genotype Controls Cell Cycle during Spermatogenesis in the Rat. Biology of Reproduction 59, 1371–1377. 10.1095/biolreprod59.6.1371.

36. Yoshida, S., Sukeno, M., and Nabeshima, Y. -i. (2007). A Vasculature-Associated Niche for Undifferentiated Spermatogonia in the Mouse Testis. Science 317, 1722–1726. 10.1126/science.1144885.

37. Komeya, M., Kimura, H., Nakamura, H., Yokonishi, T., Sato, T., Kojima, K., Hayashi, K., Katagiri, K., Yamanaka, H., Sanjo, H., et al. (2016). Long-term *ex vivo* maintenance of testis tissues producing fertile sperm in a microfluidic device. Scientific Reports 6, 21472. 10.1038/srep21472.

38. Chihara, M., Ikebuchi, R., Otsuka, S., Ichii, O., Hashimoto, Y., Suzuki, A., Saga, Y., and Kon, Y. (2013). Mice Stage-Specific Claudin 3 Expression Regulates Progression of Meiosis in Early Stage Spermatocytes1. Biology of Reproduction 89, 1–12. 10.1095/biolreprod.113.107847.

39. Mark, M., Teletin, M., Vernet, N., and Ghyselinck, N.B. (2015). Role of retinoic acid receptor (RAR) signaling in post-natal male germ cell differentiation. Biochimica et Biophysica Acta (BBA) - Gene Regulatory Mechanisms 1849, 84–93. 10.1016/j.bbagrm.2014.05.019.

40. Nebel, B.R., Amarose, A.P., and Hackett, E.M. (1961). Calendar of Gametogenic Development in the Prepuberal Male Mouse. Science 134, 832–833. 10.1126/science.134.3482.832.

41. Drumond, A.L., Meistrich, M.L., and Chiarini-Garcia, H. (2011). Spermatogonial morphology and kinetics during testis development in mice: a high-resolution light microscopy approach. Reproduction 142, 145–155. 10.1530/REP-10-0431.

42. Huckins, C., and Oakberg, E.F. (1978). Morphological and quantitative analysis of spermatogonia in mouse testes using whole mounted seminiferous tubules. I. The normal testes. Anat. Rec. 192, 519–527. 10.1002/ar.1091920406.

43. Hara, K., Nakagawa, T., Enomoto, H., Suzuki, M., Yamamoto, M., Simons, B.D., and Yoshida, S. (2014). Mouse Spermatogenic Stem Cells Continually Interconvert between Equipotent Singly Isolated and Syncytial States. Cell Stem Cell 14, 658–672. 10.1016/j.stem.2014.01.019.

44. Benda, C. (1887). Untersuchungen über den Bau des funktionirenden Samenkanälchens einiger Säugethiere und Folgerungen für die Spermatogenese dieser Wirbelthierklasse. Archiv f. mikrosk. Anatomie 30, 49–110. 10.1007/BF02955609.

45. Ronneberger, O., Fischer, P., and Brox, T. (2015). U-Net: Convolutional Networks for Biomedical Image Segmentation. In Medical Image Computing and Computer-Assisted Intervention – MICCAI 2015 Lecture Notes in Computer Science., N. Navab, J. Hornegger, W. M. Wells, and A. F. Frangi, eds. (Springer International Publishing), pp. 234–241. 10.1007/978-3-319-24574-4_28.

46. Kuramoto, Y. (1984). Chemical Oscillations, Waves, and Turbulence (Springer-Verlag) 10.1007/978-3-642-69689-3.

47. Sakaguchi, H., Shinomoto, S., and Kuramoto, Y. (1988). Mutual Entrainment in Oscillator Lattices with Nonvariational Type Interaction. Progress of Theoretical Physics 79, 1069–1079. 10.1143/PTP.79.1069.

48. Novák, B., and Tyson, J.J. (2008). Design principles of biochemical oscillators. Nat Rev Mol Cell Biol 9, 981–991. 10.1038/nrm2530.

49. Vernet, N., Dennefeld, C., Rochette-Egly, C., Oulad-Abdelghani, M., Chambon, P., Ghyselinck, N.B., and Mark, M. (2006). Retinoic Acid Metabolism and Signaling Pathways in the Adult and Developing Mouse Testis. Endocrinology 147, 96–110. 10.1210/en.2005-0953.

50. Morrison, S.J., and Spradling, A.C. (2008). Stem Cells and Niches: Mechanisms That Promote Stem Cell Maintenance throughout Life. Cell 132, 598–611. 10.1016/j.cell.2008.01.038.

51. Turing, A.M. (1952). The chemical basis of morphogenesis. Philosophical Transactions of the Royal Society of London. Series B, Biological Sciences 237, 37–72. 10.1098/rstb.1952.0012.

52. Sasai, Y. (2013). Cytosystems dynamics in self-organization of tissue architecture. Nature 493, 318–326. 10.1038/nature11859.

53. Palmeirim, I., Henrique, D., Ish-Horowicz, D., and Pourquié, O. (1997). Avian hairy Gene Expression Identifies a Molecular Clock Linked to Vertebrate Segmentation and Somitogenesis. Cell 91, 639–648. 10.1016/S0092-8674(00)80451-1.

54. Tsiairis, C.D., and Aulehla, A. (2016). Self-Organization of Embryonic Genetic Oscillators into Spatiotemporal Wave Patterns. Cell 164, 656–667. 10.1016/j.cell.2016.01.028.

55. Biga, V., Hawley, J., Soto, X., Johns, E., Han, D., Bennett, H., Adamson, A.D., Kursawe, J., Glendinning, P., Manning, C.S., et al. (2021). A dynamic, spatially periodic, micro-pattern of HES5 underlies neurogenesis in the mouse spinal cord. Molecular Systems Biology 17, e9902. 10.15252/msb.20209902.

56. Tomchik, K.J., and Devreotes, P.N. (1981). Adenosine 3ʹ,5ʹ-Monophosphate Waves in Dictyostelium discoideum: a Demonstration by Isotope Dilution—Fluorography. Science 212, 443–446. 10.1126/science.6259734.

57. Sawai, S., Thomason, P.A., and Cox, E.C. (2005). An autoregulatory circuit for long-range self-organization in Dictyostelium cell populations. Nature 433, 323–326. 10.1038/nature03228.

58. Tam, P.P.L. (1981). The control of somitogenesis in mouse embryos. Development 65, 103–128. 10.1242/dev.65.Supplement.103.

59. Kageyama, R., Niwa, Y., Isomura, A., González, A., and Harima, Y. (2012). Oscillatory gene expression and somitogenesis. WIREs Developmental Biology 1, 629–641. 10.1002/wdev.46.

60. Ikami, K., Tokue, M., Sugimoto, R., Noda, C., Kobayashi, S., Hara, K., and Yoshida, S. (2015). Hierarchical differentiation competence in response to retinoic acid ensures stem cell maintenance during mouse spermatogenesis. Development 142, 1582–1592. 10.1242/dev.118695.

61. Ebner, V. von (1902). Männliche Geschlechtsorgane. In Handbuch der Gewebelehre des Menschen, A. Kölliker and V. von Ebner, eds. (Leipzig : W. Engelmann), pp. 402–505.

62. Roosen-Runge, E.C., and Barlow, F.D. (1953). Quantitative studies on human spermatogenesis. I. Spermatogonia. American Journal of Anatomy 93, 143–169. 10.1002/aja.1000930202.

63. Schulze, W., and Rehder, U. (1984). Organization and morphogenesis of the human seminiferous epithelium. Cell Tissue Res. 237. 10.1007/BF00228424.

64. Johnson, L., Mckenzie, K.S., and Snell, J.R. (1996). Partial wave in human seminiferous tubules appears to be a random occurrence. Tissue and Cell 28, 127–136. 10.1016/S0040-8166(96)80001-2.

65. Lin, M., and Jones, R.C. (1990). Spatial arrangement of the stages of the cycle of the seminiferous epithelium in the Japanese quail, Coturnix coturnix japonica. Reproduction 90, 361–367. 10.1530/jrf.0.0900361.

66. Stenn, K.S., and Paus, R. (2001). Controls of Hair Follicle Cycling. Physiological Reviews 81, 449–494. 10.1152/physrev.2001.81.1.449.

67. Suzuki, N., Hirata, M., and Kondo, S. (2003). Traveling stripes on the skin of a mutant mouse. Proc Natl Acad Sci USA 100, 9680–9685. 10.1073/pnas.1731184100.

68. Ma, L., Liu, J., Wu, T., Plikus, M., Jiang, T.-X., Bi, Q., Liu, Y.-H., Müller-Röver, S., Peters, H., Sundberg, J.P., et al. (2003). “Cyclic alopecia” in Msx2 mutants: defects in hair cycling and hair shaft differentiation. Development 130, 379–389. 10.1242/dev.00201.

69. Plikus, M.V., Widelitz, R.B., Maxson, R., and Chuong, C.-M. (2009). Analyses of regenerative wave patterns in adult hair follicle populations reveal macroenvironmental regulation of stem cell activity. Int. J. Dev. Biol. 53, 857–868. 10.1387/ijdb.072564mp.

70. Nakagawa, T., Jörg, D.J., Watanabe, H., Mizuno, S., Han, S., Ikeda, T., Omatsu, Y., Nishimura, K., Fujita, M., Takahashi, S., et al. (2021). A multistate stem cell dynamics maintains homeostasis in mouse spermatogenesis. Cell Reports 37. 10.1016/j.celrep.2021.109875.

71. Schindelin, J., Arganda-Carreras, I., Frise, E., Kaynig, V., Longair, M., Pietzsch, T., Preibisch, S., Rueden, C., Saalfeld, S., Schmid, B., et al. (2012). Fiji: an open-source platform for biological-image analysis. Nat Methods 9, 676–682. 10.1038/nmeth.2019.

72. Hörl, D., Rusak, F.R., Preusser, F., Tillberg, P., Randel, N., Chhetri, R.K., Cardona, A., Keller, P.J., Harz, H., Leonhardt, H., et al. (2019). BigStitcher: reconstructing high-resolution image datasets of cleared and expanded samples. Nat Methods 16, 870–874. 10.1038/s41592-019-0501-0.

73. Miura, K. (2020). Bleach correction ImageJ plugin for compensating the photobleaching of time-lapse sequences. F1000Res 9, 1494. 10.12688/f1000research.27171.1.

74. Thevenaz, P., Ruttimann, U.E., and Unser, M. (1998). A pyramid approach to subpixel registration based on intensity. IEEE Transactions on Image Processing 7, 27–41. 10.1109/83.650848.

75. Ahmed, E.A., and de Rooij, D.G. (2009). Staging of Mouse Seminiferous Tubule Cross-Sections. In Meiosis, S. Keeney, ed. (Humana Press), pp. 263–277. 10.1007/978-1-60761-103-5_16.

76. Meistrich, M.L., and Hess, R.A. (2013). Assessment of Spermatogenesis Through Staging of Seminiferous Tubules. In Spermatogenesis, D. T. Carrell and K. I. Aston, eds. (Humana Press), pp. 299–307. 10.1007/978-1-62703-038-0_27.

77. Nakata, H., Wakayama, T., Takai, Y., and Iseki, S. (2015). Quantitative Analysis of the Cellular Composition in Seminiferous Tubules in Normal and Genetically Modified Infertile Mice. J Histochem Cytochem. 63, 99–113. 10.1369/0022155414562045.

78. Parvinen, M., and Hecht, N.B. (1981). Identification of living spermatogenic cells of the mouse by transillumination-phase contrast microscopic technique for ‘in situ’ analyses of DNA polymerase activities. Histochemistry 71, 567–579. 10.1007/BF00508382.

79. Kotaja, N., Kimmins, S., Brancorsini, S., Hentsch, D., Vonesch, J.-L., Davidson, I., Parvinen, M., and Sassone-Corsi, P. (2004). Preparation, isolation and characterization of stage-specific spermatogenic cells for cellular and molecular analysis. Nat Methods 1, 249–254. 10.1038/nmeth1204-249.

80. Mäkelä, J.-A., Cisneros-Montalvo, S., Lehtiniemi, T., Olotu, O., La, H.M., Toppari, J., Hobbs, R.M., Parvinen, M., and Kotaja, N. (2020). Transillumination-Assisted Dissection of Specific Stages of the Mouse Seminiferous Epithelial Cycle for Downstream Immunostaining Analyses. JoVE (Journal of Visualized Experiments), e61800. 10.3791/61800.

81. Tinevez, J.-Y., Perry, N., Schindelin, J., Hoopes, G.M., Reynolds, G.D., Laplantine, E., Bednarek, S.Y., Shorte, S.L., and Eliceiri, K.W. (2017). TrackMate: An open and extensible platform for single-particle tracking. Methods 115, 80–90. 10.1016/j.ymeth.2016.09.016.

82. Huckins, C. (1978). Spermatogonial intercellular bridges in whole-mounted seminiferous tubules from normal and irradiated rodent testes. Am. J. Anat. 153, 97–121. 10.1002/aja.1001530107.

83. Yoshida, S., Takakura, A., Ohbo, K., Abe, K., Wakabayashi, J., Yamamoto, M., Suda, T., and Nabeshima, Y. (2004). Neurogenin3 delineates the earliest stages of spermatogenesis in the mouse testis. Developmental Biology 269, 447–458. 10.1016/j.ydbio.2004.01.036.

84. Soroldoni, D., Jörg, D.J., Morelli, L.G., Richmond, D.L., Schindelin, J., Jülicher, F., and Oates, A.C. (2014). A Doppler effect in embryonic pattern formation. Science 345, 222–225. 10.1126/science.1253089.

85. La, H.M., Mäkelä, J.-A., Chan, A.-L., Rossello, F.J., Nefzger, C.M., Legrand, J.M.D., De Seram, M., Polo, J.M., and Hobbs, R.M. (2018). Identification of dynamic undifferentiated cell states within the male germline. Nat Commun 9, 2819. 10.1038/s41467-018-04827-z.

## References

[1] Y. Kuramoto. Chemical Oscillations, Waves, and Turbulence. Springer, 1984.

[2] H. Sakaguchi, S. Shinomoto, and Y. Kuramoto. Mutual entrainment in oscillator lattices with nonvariational type interaction. Progress of theoretical physics, 79(5):1069–1079, 1988.

[3] B. Blasius and R. Tonjes. Quasiregular concentric waves in heterogeneous lattices of coupled oscillators. Physical review letters, 95(8):084101, 2005.

[4] J. P. Moroney and P. R. Eastham. Synchronization in disordered oscillator lattices: Nonequilibrium phase transition for driven-dissipative bosons. Physical Review Research, 3(4):043092, 2021.

[5] M. Pinsky and S. Karlin. An introduction to stochastic modeling. Academic press, 2010.

[6] H. Nakata, T. Sonomura, and S. Iseki. Three-dimensional analysis of seminiferous tubules and spermatogenic waves in mice. Reproduction, 154(5):569–579, 2017.

[7] E. F. Oakberg. Duration of spermatogenesis in the mouse and timing of stages of the cycle of the seminiferous epithelium. American Journal of Anatomy, 99(3):507–516, 1956.

[8] Y. Clermont and M. Trott. Duration of the cycle of the seminiferous epithelium in the mouse and hamster determined by means of 3h-thymidine and radioautography. Fertility and Sterility, 20(5):805–817, 1969.

[9] R. Sugimoto, Y. Nabeshima, and S. Yoshida. Retinoic acid metabolism links the periodical differentiation of germ cells with the cycle of sertoli cells in mouse seminiferous epithelium. Mechanisms of Development, 128(11-12):610–624, 2012.

[10] N. Vernet, C. Dennefeld, C. Rochette-Egly, M. Oulad-Abdelghani, P. Chambon, N. B. Ghyselinck, and M. Mark. Retinoic acid metabolism and signaling pathways in the adult and developing mouse testis. Endocrinology, 147(1):96–110, 2006.

[11] J. D. Murray. Mathematical biology: I. An introduction. Springer, 2002.

[12] K. Ikami, M. Tokue, R. Sugimoto, C. Noda, S. Kobayashi, K. Hara, and S. Yoshida. Hierarchical differentiation competence in response to retinoic acid ensures stem cell maintenance during mouse spermatogenesis. Development, 142(9):1582–1592, 2015.

